# Even allocation of benefits stabilizes microbial community engaged in metabolic division of labor

**DOI:** 10.1101/2021.09.10.459869

**Authors:** Miaoxiao Wang, Xiaoli Chen, Xiao-nan Liu, Yuan Fang, Xin Zheng, Ting Huang, Yue-Qin Tang, Yong Nie, Xiao-Lei Wu

## Abstract

Metabolic division of labor (MDOL) is commonly observed in microbial communities, where a metabolic pathway is sequentially implemented by several members similar to an assembly line. Uncovering the assembly rules of the community engaged in MDOL is crucial for understanding its ecological contribution, as well as for engineering high-performance microbial communities. To investigate the assembly of the community engaged in MDOL, we combined mathematical modelling with experimentations using synthetic microbial consortia. We built a theoretical framework predict the assembly of MDOL system, and derived a simple rule: to maintain co-existence of the MDOL members, the populations responsible for former steps should hold a growth advantage (*m*) over the ‘private benefit’ (*n*) of the population responsible for last step, and the steady-state frequency of the last population is determined by the quotient of *n* and *m*. Our experiments further indicated that our theoretical framework accurately predicted the stability and assembly of our engineered synthetic consortia that degrade naphthalene through two-step or multi-step MDOL. Our results demonstrate that the assembly of microbial community engaged in MDOL is determined by a limited number of parameters. This quantitative understanding provides novel insights on designing and managing stable microbial systems to address grand challenges facing human society in agriculture, degradation of the environment, and human health.

## Introduction

Microorganisms colonize all major ecological niches on our planet, from deep terrestrial biosphere miles beneath the land surface^1-4^, to the intestinal tracts of mammalians^5,6^. In order to survive in these ever-changing habitats, microorganisms have evolved highly sophisticated characteristics to accomplish complex metabolic tasks for biochemical transformations, such as converting an unavailable resource into a substrate suitable for cell growth. As a result, their metabolic activities drive global biogeochemical cycles, and thus profoundly influence the health of both ecosystems as well as their inhabitants^7^.

Most of these metabolic tasks are accomplished through long metabolic pathways, which are performed by a single microbial population. Alternatively, these tasks are divided across different interacting populations complementarily at the community level, a phenomenon called metabolic division of labor (MDOL)^8-11^. The former requires a series of enzymes being produced by a single population, and thus create a substantial metabolic burden that limits the productivity and growth of the population carrying out these tasks^9,10^. In contrast, if a long pathway is distributed among different populations in communities engaged in MDOL (simplified as MDOL communities thereafter), each member only needs to specialize for its respective metabolic step. Since each population only contains a subset of genetic components required for the overall pathway, MDOL is thought to be a key evolutionary strategy to reduce metabolic burden^12,13^.

Several studies have shown that numerous ecologically and environmentally important pathways are accomplished through MDOL. In particular, microbial degradation of complex organic compounds is frequently executed through MDOL. For example, the gut communities comprised of microbial symbionts digest plant polysaccharides into either sugars or short chain fatty acids in a MDOL manner, which are then absorbed by the host cells^14,15^. During the Deepwater Horizon oil spill Gulf of Mexico (2010), complete degradation of polycyclic aromatic hydrocarbons (PAHs) required the partitioning of key pathway steps into different bacterial groups^16^. In more specific cases, syringate can be degraded through MDOL between *Acetobacterium woodii* and *Pelobacter acidigallici*^17^; *Marinobacter* is only responsible for a subset of tasks in phenanthrene degradation, while other marine bacteria perform the remaining steps^18^. As the proper functioning of a microbial community is determined by its compositional makeup^19,20^, it is critical to understand and predict the ecological functions of MDOL communities. To this end, uncovering how MDOL systems are stabilized and how they influence community dynamics is of paramount importance.

Inspired by natural MDOL communities, numerous studies have recently explored how to adopt their MDOL strategies for the removal of organic pollutants^21^. For instance, one study engineered a defined consortium composed of an *Escherichia coli* strain and a *Pseudomonas aeruginosa* strain for phenanthrene bio-removal via MDOL^22^. Another study investigated how MDOL in a consortium composed of *Stenotrophomonas* sp. N5 and *Advenella* sp. B9 affects the phenol biodegradation^23^. These studies mainly tested whether MDOL enhances the efficiency of biodegradation compared to relevant monocultures comprised of single species. However, specific strains may not be able to stably co-exist in an artificial co-culture system^24-27^, resulting in the collapse of the community. To fully understand how MDOL systems are stabilized and assembled, it is essential to build quantitative rules that reliably forecast whether a specific combination of strains can successfully assemble into a robust MDOL system. Establishing these rules is vital to rationally adopt MDOL strategies to engineer high-performance microbial communities.

Our recent work combined mathematical modelling and experiments with a synthetic consortium^28^. We found that the traits of the substrate, such as its concentration and toxicity, largely affected the assembly of the MDOL community. Nevertheless, a mass of biotic and abiotic factors, such as metabolic burden of each population, mass transfer rate, as well as the toxicity of substrate or intermediates, may also contribute to the assembly of MDOL community. A general rule combining all of these factors remains absent, rendering any prediction of how MDOL communities assemble challenging. Here, we bridged this gap using a bottom-up approach. We first built an ordinary differential equation (ODE) model to formalize a rule that forecasted how MDOL community assembled. We next tested this rule by performing *in vitro* assays using a series of engineered synthetic consortia implementing naphthalene degradation through two- or multi-step MDOL.

## Results

### Assembly and stability of the microbial community engaged in two-step MDOL

#### Model framework for two-step MDOL

To depict the dynamics of the community engaged in metabolic division of labor (MDOL community), we conceptualized the degradation of an organic compound implemented by a microbial consortium composed of two populations using a simple mathematical model (Figure 1A). In this consortium, the first population (named as [1, 0]) expressed an enzyme (E1) to catalyze the conversion of an organic substrate (S) into an intermediate metabolite (I), while the second population (named as [0, 1]) was responsible for the subsequent conversion of I to a final product (P) by expressing another enzyme (E2). In the basic model, transport of S, I, and P across the cell membrane was assumed to occur via passive diffusion mediated by coefficients *γ*_*s*_, *γ*_*i*_, and *γ*_*p*_. Following these assumptions, we formulated the dynamics of intracellular and extracellular I and P by using seven ordinary differential equations (ODEs) (Eqns. [8]-[13] in Methods section). Consistent with our previous hypotheses and observations^28^, we next assumed that P, which was produced by the second population, was the sole resource for the growth of both populations, while neither S nor I could be directly used for growth. As a consequence, [0, 1] possesses a preferential access to the final product (P), resulted in a ‘private benefit’ derived from the product privatization (Figure 1A). Details about the model construction are described in the Method Section, as well as in the Supplementary Information S1. Meanings of the variables and parameters of the model were listed in Table S1 and S2.

**Figure 1.**
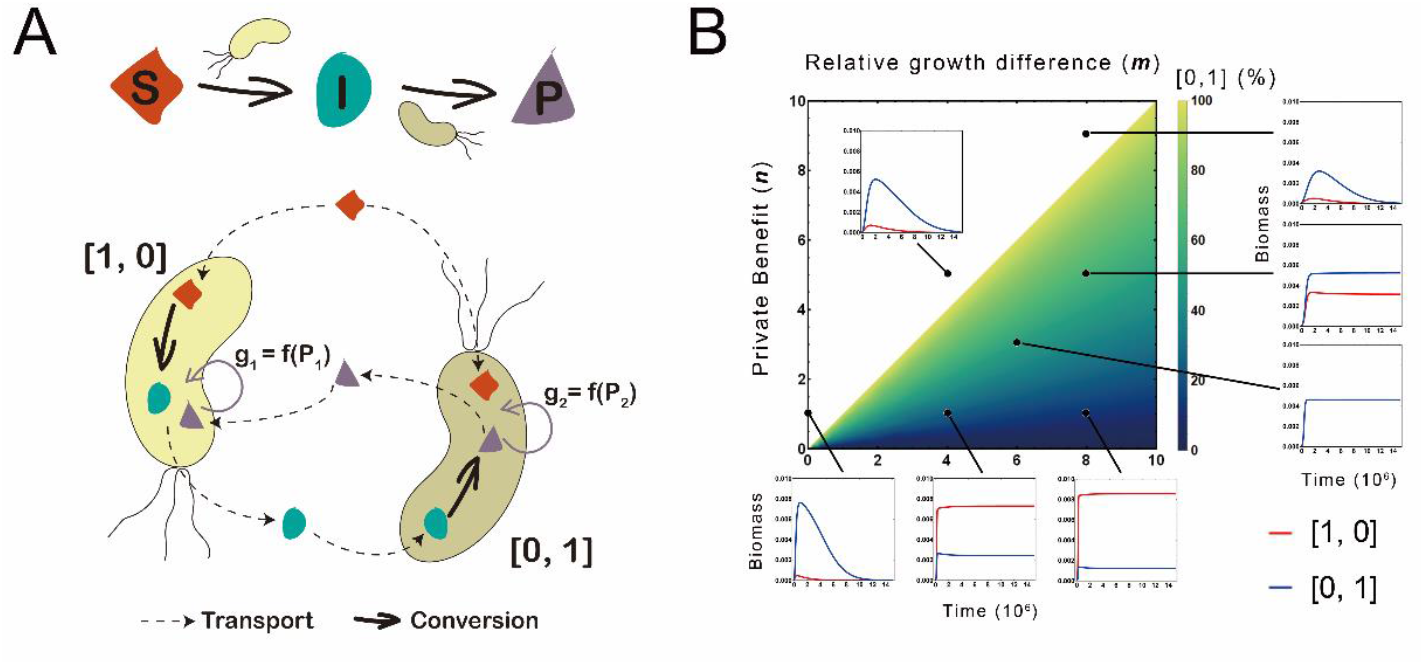
Assembly rule of microbial community engaged in two-step MDOL. (A) Schematic diagram showing the assumptions of our basic model. We assumed a conceptualized organic substrate (S) could be degraded into an intermediate metabolite (I) by a population [1, 0], then to a final product (P) by the other population [0, 1]. All the reactions occurred intracellularly, while S, I, and P passively diffused across the cell membrane. Importantly, the growth of both populations is dependent on the intracellular concentration of P, which is the sole limiting resource of this system. (B) The density map indicates the assembly rule of the two-step MDOL community. The value of privatization benefit (*n*) and relative growth advantage (*m*) determines whether [1, 0] and [0, 1] can stably co-exist (shown by the colorized space), as well as the relative abundance of [0, 1] in the stable community (shown by the color gradient). The corresponding curve graph illustrates the community dynamics of the community under six specific *n* and *m* combinations. In these simulations, the growth cost of the population [0,1] (*c*_*2*_) was adjusted to modify *m*, and the diffusion coffecient of the final product (*γ*_*p*_) was adjusted to to modify *n*. The default values listed in Table S2 were assigned for other parameters.

#### Deriving the criterion for maintaining a stable two-step MDOL community

To derive the criterion for stable two-step MDOL community, we solved steady-state expressions of Eqns. [8]-[17] (Supplementary Information S1.3). We obtained a simple formula, which suggests that if the steady-state of a two-step MDOL community exists, the relative abundance of the [0, 1] (*R*_[0,1]_) should follow:

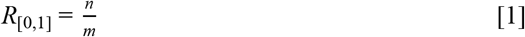

Here *n = Ig*_*1*_*/γ*_*p*_ is the ‘Product demand gap’ of [1, 0], reflecting the ‘private benefit’ of [0, 1] derived from the product privatization, while 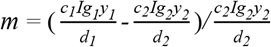, is the normalized difference between the inherent growth rates of the two populations (see Supplementary Information S1.3 for further explanation). Importantly, since *R*_[0,1]_ ranges from 0 to 1, we derived a prerequisite defining the conditions when the two populations were able to stably co-exist:

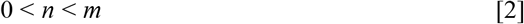

Eqn. [2] represents two meanings. Firstly, the requirement, *m* > 0, suggests that [1, 0] must hold a higher growth advantage than [0, 1]. Secondly, the requirement, *n < m*, indicates that ‘private benefit’ of [0, 1] must lower than the growth advantage of [1, 0]. We visualized these formulations by a two-dimensional density map. As shown in Figure 1B, MDOL community succeeded to a steady-state only when the values of *n* and *m* fell inside the range defined by Eqn. [2], and the community assembly at the steady-state can be directly assessed by Eqn. [1]. We next performed additional analyses to test whether other parameters that are not included in *n* and *m* also affect the proposed rule (Supplementary Information S1.4; Figure S1). Our results indicated that the stability of the MDOL community also requires speed of the first reaction (*a*_*1*_) reaches a threshold (Figure S1A-B).

In summary, our mathematical modelling formulized a rule that defines the condition when the two populations in the MDOL community can stably co-exist, namely when the population [1, 0] possesses a growth advantage that outweighs the ‘private benefit’ of the population [0, 1] (Eqn. [2]). When the steady-state exists, this rule also provides a prediction on the steady-state community structure (Eqn. [1]).

#### Stability of MDOL in response to changing metabolic conditions

Our basic model only considers the effects of 17 basic parameters (Table S2). To investigate how our general rule responds to changing metabolic conditions associated with more specific parameters, we introduced additional assumptions regarding complex pathway mechanisms hitherto excluded into the basic equations. Our model analyses indicated that these additional pathway mechanisms affect the basic assembly rule of MDOL community. In general, the scenarios that benefits [1, 0] relaxed the parameter areas for stable MDOL communities, whereas scenarios that benefits [0, 1] tightened parameter areas (Supplementary information S2).

1. Our basic model assumed that the intermediate (I) and product (P) are transported across the cell membrane by passive diffusion. When we assumed that the final product (P) is actively absorbed by [1, 0], or is actively secreted by [0, 1] (Figure 2A; Figure S2A-B), the MDOL community becomes more likely to reach a steady-state (Supplementary Information 2.1.2-2.1.3; Figure S2C).
2. Both metabolic reactions involved in a MDOL pathway might be performed extracellularly^29^ (Figure 2B; Figure S3A-B). When only the first metabolic step is catalyzed extracellularly, the basic prediction of our rule remains unchanged (Supplementary Information S2.2.1; Figure S3). However, once the second metabolic step is performed extracellularly, the rule change considerably. Our results indicated that the coexistence of the two populations is only present if the two populations exhibit similar relative fitness (as defined by Eqn. [S2.35]). In addition, the assembly of the stable MDOL community is strongly dependent on the initial abundance of the two populations (Supplementary Information S2.2; Figure S3B-C).
3. The intermediates of a MDOL pathway can be converted to chemicals unavailable to microorganisms via spontaneous reactions^30,31^ (Figure 2C; Figure S4A). When we included spontaneous conversion of I, P, or both in our model, we found that achieving stability of MDOL community became more untenable (Supplementary Information S2.3; Figure S3B).
4. Metabolic by-products may be generated from the first-step reaction of MDOL pathway^31-33^ (Figure 2D; Figure S5A). Under this condition, the ‘private benefit’ of [0, 1] was counteracted by the byproduct, strongly favoring [1, 0] and co-existence of the two populations (Supplementary Information S2.7; Figure S4B).
5. Toxic effects of substrate^34-36^, intermediates^34,35,37^, and final product^38,39^ are commonly occur during microbial degradation of organic compounds (Figure 2E-G; Figure S6-8A). While substrate toxicity neutralizes the ‘private benefit’ of [0, 1] (Supplementary Information S2.8.1; Figure S6B), the presence of intermediates toxicity offers an additional benefit to [0, 1], leading to a more rigorous criterion for stabilizing the community (Supplementary Information S2.8.2; Figure S7B-D).

**Figure 2.**
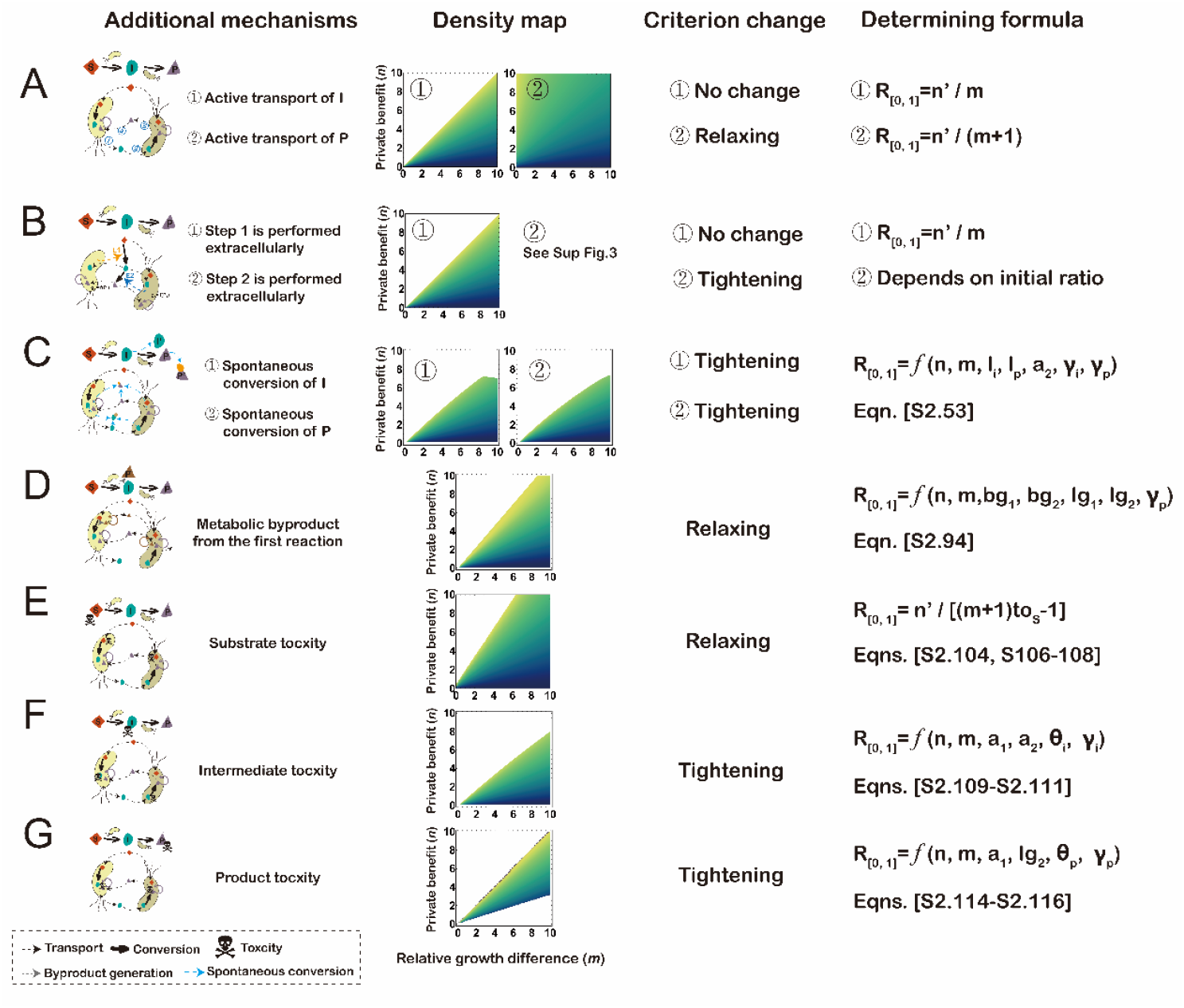
Additional pathway mechanisms influence the assembly of the two-step MDOL community. Seven additional pathway mechanisms were tested: Different configurations for the transport of the intermediate and product (A); Alternative or both metabolic reactions occuring in the extracelluar space (B); Presence of abiotic conversion of intermediate and product (C); Presence of byproduct generated from the first reaction (D); Presence of biotoxicity of substrate (E); Presence of biotoxicity of intermediate (F); Presence of biotoxicity of end product (G). First column: Architectures of two-step MDOL community considering these seven pathway mechanisms. Second column: Representative density maps shows how the additional pathway mechanisms change the assembly rules. Third column: Summary of the how the additional pathway mechanisms change the criterion for the stability of the community. ‘Tightening’ means that the size of the parameter space required for the stability decreases, thus the two-step MDOL community becomes more difficult to maintain stability. In contrast, ‘Relaxing’ means that the size of the parameter space required for the stability increases, thus the two-step MDOL community becomes easier to maintain stability. Fourth column: Summary of the derived formulas that determine the assembly of two-step MDOL community at steady-state in each scenario. For simplicity, the key impacting factors of the scenarios in (C), (D), (F), and (G) are listed. Detailed expressions of these formulas are available in Supplementary Information S2, of which the tracking number are listed.

#### Experimentally testing the rules of community assembly and stability achieved by synthetic microbial consortia

To experimentally test our proposed rule, we engineered three synthetic consortia that degrade naphthalene via two-step MDOL. In these systems, the first *Pseudomonas stutzeri* strain converts naphthalene into its intermediate (i.e., 1, 2-hydroxynaphthalene, salicylate, or catechol), which are exchanged among different cells^33,40^ but cannot be directly used as the carbon source to support bacterial growth. The second strain possesses ability to degrade the intermediate to the final products (pyruvate and acetyl-CoA), which are then partially secreted to the environment and utilized by the consortia as the limiting carbon source (Figure 3A; Figure S9A; Figure S10A; see Supplementary Information S4.1 for the strain construction). To predict the assembly of these synthetic ecosystems, we modified our basic model to include specific pathway mechanisms corresponding to these consortia (see Supplementary Information S4.2 for details about the modifications), and mathematically derived the criteria governing the assembly of these consortia (Figure 3B; Figure S9B; Figure S10B).

**Figure 3.**
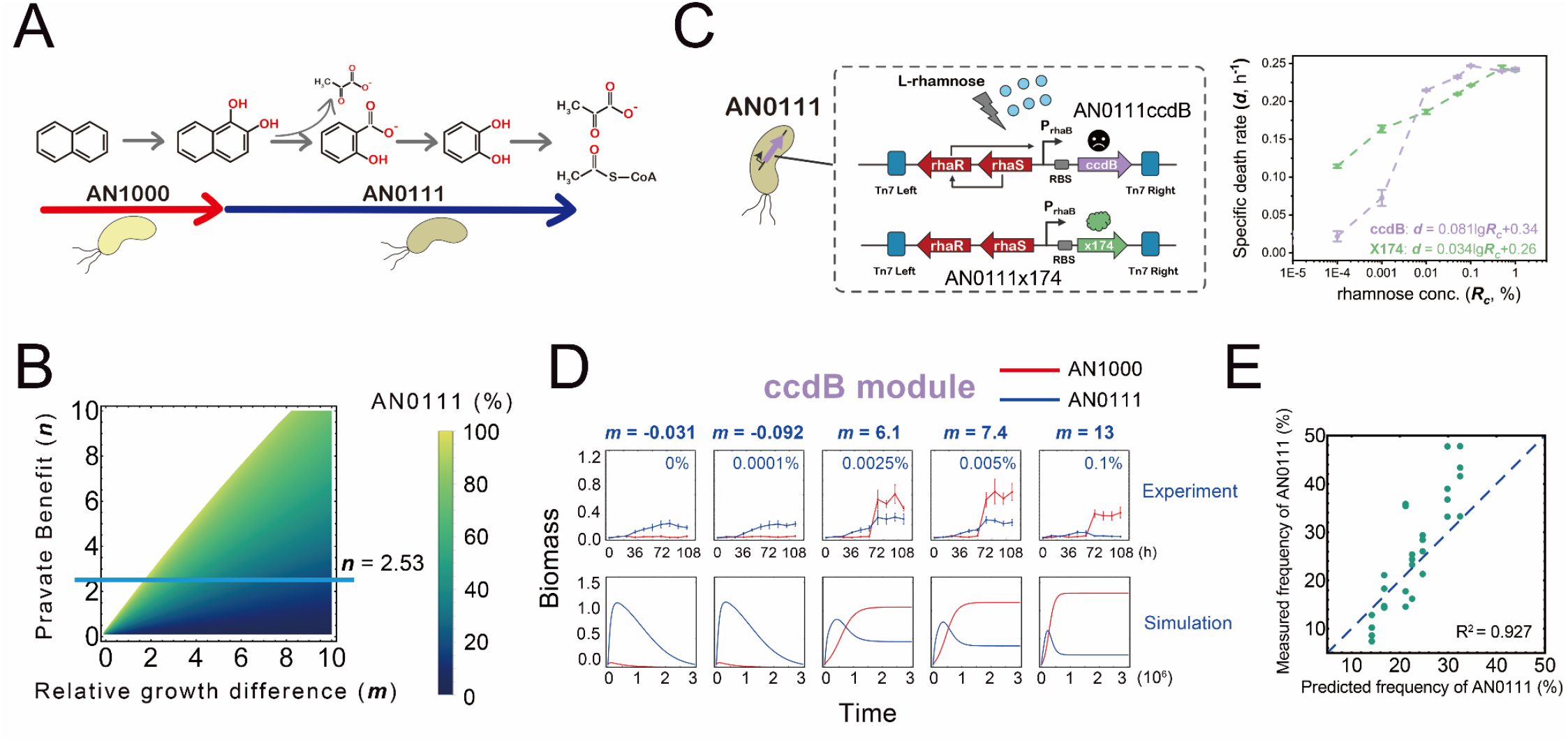
Dynamics of the synthetic community composed of *P. stutzeri* AN1000 and *P. stutzeri* AN0111ccdB. (A) Schematic diagram of the construction of the synthetic consortium. (B) Predicting the assembly of the synthetic community by mathematical modelling. The density map shows the parameter space in which the community can maintain (the colorized space), as well as the relative abundance of *P. stutzeri* AN0111ccdB or *P. stutzeri* AN0111×174 in the stable community (the color gradient). The blue line suggests that in our synthetic community *n* = 2.53, which is the region we performed experimental verification. (C) To experimentally modify *m* value, two genetic modules were introduced into strain *P. stutzeri* AN0111, generating *P. stutzeri* AN0111ccdB and *P. stutzeri* AN0111×174, in which the expression of toxic protein, CcdB or X174, are controllably induced by rhamnose. Therefore, the death rate of the modified strain can be quantitatively modulated by adjusting rhamnose concentration, as shown in the left plot and Table S7. (D) The dynamics of the synthetic community composed of *P. stutzeri* AN1000 and *P. stutzeri* AN0111ccdB from co-culture experiments and mathematical modelling under different rhamnose concentrations (that is, a gradient of *m* values). To measure their relative abundance, the strain performing the first step was labeled with MCherry, while the other strain was labeled with EGFP. (E) Testing the predicting power of our mathematical frameworks. The experimentally measured frequency of the *P. stutzeri* AN0111ccdB (Figure S14) and *P. stutzeri* AN0111×174 (Figure S15) in dilution-transfer experiments are compared with the predicted frequency from our mathematical framework. The values of relative abundance at the end of the third transfer were recorded. Each green dot indicates one experimental replicate. The blue dashed line indicates the line in which the experimental results and predicted results are identical. The adjust R^2^ are aquired from statistical fit using NonlinearModelfit function of *Worfram Mathematica*.

Setting the consortium composed of strain AN1000 and strain AN0111 (Figure 3A) as an example, our modelling analyses suggest that this system is more likely to be stable compared with our basic rule (Figure 1B), mainly due to the toxic effect of naphthalene. To test the predicting power of our theoretical criterion, we cultured the consortium using naphthalene as the sole carbon source. Since the two strains exhibited a similar inherent fitness (*m* = −0.031), the consortia were found to be ecologically unstable (Figure S12A), consistent with the prediction of our model. To obtain a stable MDOL community, we changed the relative inherent fitness of the two populations (the value of *m*) according to our theoretical framework. To this end, we introduced two genetic modules to modify the inherent fitness of the strain AN0111 (see Supplementary Information S4.1.4; Figure 3A and 3C; Table S7). We co-cultured the modified strain AN0111 with strain AN1000, mimicking the community dynamics with a given *n* value of 2.53, (estimated by the experimental measurement of *Ig* divided by previously reported value of *γ*_*p*_^41^; see Supplementary information S4.2.2 for details) and a gradient of *m* values (The blue line in Figure 3B). The results showed that when the value of *m* is lower than the threshold (that is, when *m* < 1.7), at which our models predict that the two populations fail to stably co-exist, our synthetic consortium collapsed (Figure 3D; Figure S13A; Figure S14A; Figure S15A). In contrast, when *m* was set over the threshold, the consortium stabilized (Figure 3D; Figure S13A), even after reaching three passaging cycles (Supplementary Fig 14A; Figure S15A). Further analyses showed that our mathematical modelling accurately predicted the steady-state frequency of strain AN0111 in the consortium (R^2^ = 0.927; Figure 3E).

We also tested our proposed rule using two additional synthetic consortia, and observed similar results. As pyruvate was produced from the conversion of 1, 2-hydroxynaphthalene to salicylate as a byproduct, our models related to these consortia predicted that these two consortia remain stable easier than the consortium composed of AN1000 and AN0111 (Figure S9B; Figure S10B). Nevertheless, our mathematical framework accurately predicted the stability and assembly of these two consortia (Figure S9-15). Together, these results indicated that our mathematical framework reliably guide to construct stable synthetic consortia engaged in MDOL, as well as accurately forecast the assembly of these consortia.

### Assembly and stability of the community engaged in multi-step MDOL

We next investigated the assembly rule of community executing multi-step MDOL, at which a long metabolic pathway is distributed among more than two populations in microbial communities^42^. To this end, we expanded our basic model (based on 2-step MDOL) to build a more complex mathematical framework (Eqns. [18]-[23] in Methods section; Supplementary Information S3), which conceptualizes the dynamics of ***N***-step MDOL community (Figure 4A). We derived a formula based on the analyses of these equations (Supplementary Information S3.2 and S3.4), which defines the steady-state frequency of the population that performs the last step in a community engaged in *N-*step MDOL:

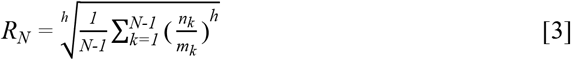

**Figure 4.**
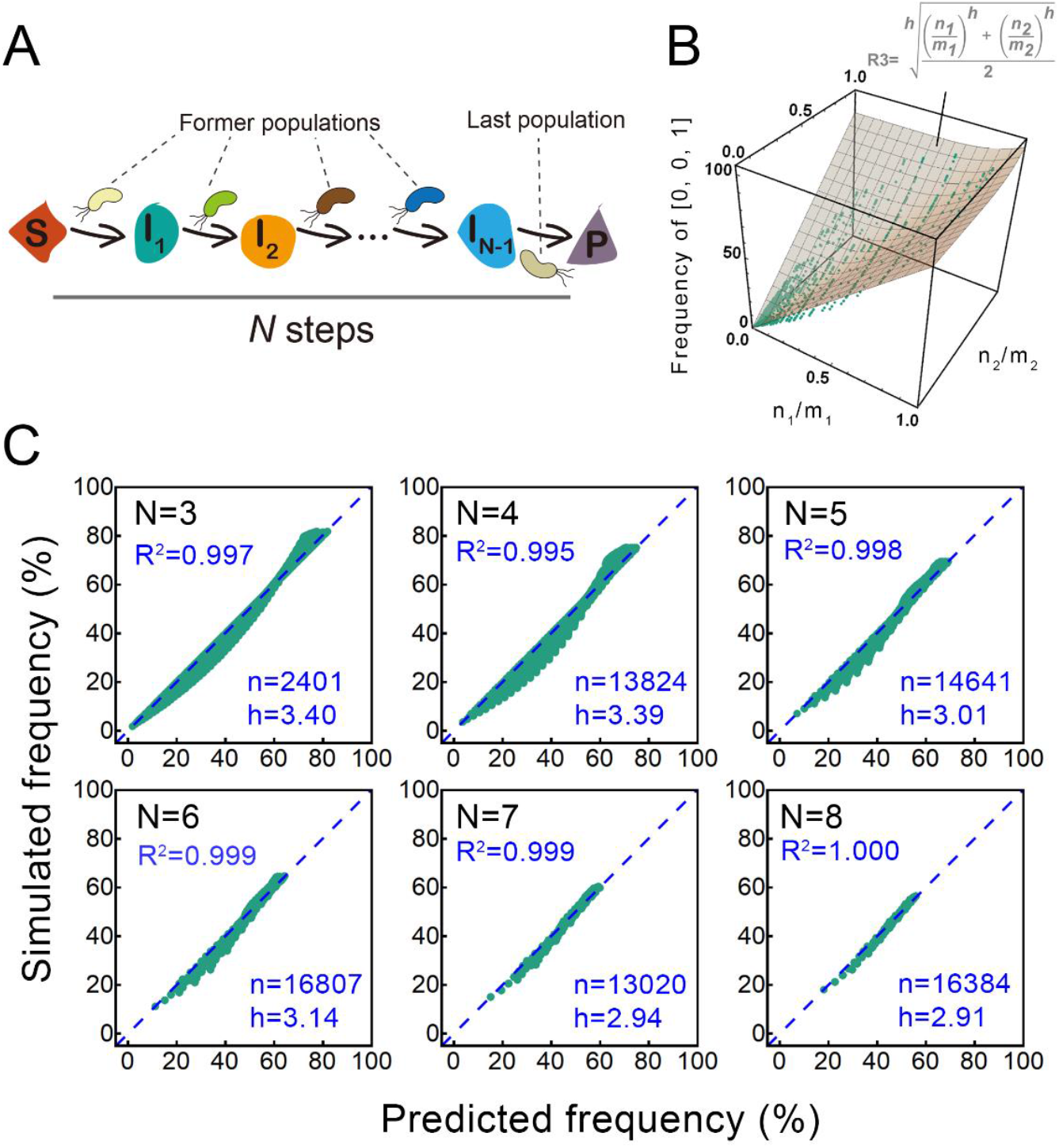
Assembly rule of microbial community engaged in multi-step MDOL. (A) Schematic diagram shows the assumptions of the model regarding the assembly of multi-step MDOL community. We assumed a conceptualized organic substrate (S) was degraded into the end product (P) through *N*-1 intermediates, and the *j*th intermediate was labelled with I_*j*_. Each reaction was carried out by one population and occurred intracellularly. (B) Assembly rule of microbial community engaged in three-step MDOL. The relationship between the relative abundance of the [0, 0, 1] and the ratio *n*_*1*_*/m*_*1*_ and *n*_*2*_*/m*_*2*_ in 890 simulated steady-state communities. Each dot shows the relative abundance of [0, 0, 1] obtained by the simulation parameterized with the corresponding value set. The surface diagram shows distribution of the relative abundance of [0, 0, 1] predicted by Eqn. [3]. Details of these simulations are provided in Supplementary Information S3.2. (C) The linear correlation between the relative abundances of the last population (*R*_*N*_) in those stable *N*-step MDOL communities predicted by Eqn. [3] and those abundances obtained by simulations. Comparisons of three- to eight-step MDOL communities are shown (meaning *N* = 3∼8). The dashed lines show the fitted linear correlation curve. *N* is the number of the simulations included, h is the optimal mean square index *h*. Details of these simulations are described in Supplementary Information S3.4.

Here, *n*_*k*_ *= Ig*_*k*_ */ γ*_*p*_, represents the ‘Product demand gap’ of the *k*th population, reflecting the relative ‘private benefit’ of the population performing the last step (simplified as the last population thereafter; Figure 4A) against the *k*th population, while 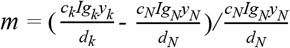, represents the normalized difference between the inherent fitness of the *k*th population and the last population; *h* is a fitted exponent affected by maximum biomass capacity and reaction speed of each metabolic step (Supplementary Information S3.2 and S3.4). Our analysis also identified the two prerequisites that define the co-existence of the populations involved in these complex systems:

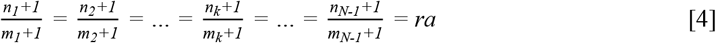

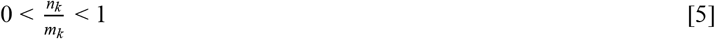

Here, *ra* represents the relative fitness of the population performing the last step to the *k*th population performing the *kth* step (see Supplementary information S3.2.1 and S3.4.1 for the definitions and discussions).

Using three-step MDOL community as an example (Figure S15A; the three population was named as [1, 0, 0], [0, 1, 0], [0, 0, 1]), our simulations showed that the three populations only co-exist when the *ra* value of [1, 0, 0] equals that of the [0, 1, 0] (prerequisite defined by Eqn. [4]; Figure S15B) and the values of *n*_*k*_*/m*_*k*_ belong to the range defined by Eqn. [5] (Figure 4B). This result suggests that, a stable community can only be achieved if the populations performing the former steps (simplified as the former populations thereafter; Figure 4A) exhibit comparable fitness levels (Eqn. [4]). In addition, these populations are required to maintain a growth advantage over that outweighs the ‘private benefit’ of the last population (Eqn. [5]). Under steady-state conditions, we successfully estimated the frequencies of the [0, 0, 1] using Eqn. [3] (Figure 4B-C). Importantly, Eqn. [3] can be expanded to estimate the results of mathematical simulations considering MDOL community with more steps (Figure 4C; up to *N*=8). Remarkably, the rule we proposed about the assembly of two-step MDOL community is a specific case of this rule (when set *N* = 2 in Eqn. [3], we obtain Eqn. [1]). Together, we successfully expanded our mathematical framework to estimate the assembly of the multi-step MDOL community.

To experimentally verify our expanded rule, we separated the naphthalene degradation pathway into four steps and engineered four *P. stutzeri* populations (*P. stutzeri* AN1000, *P. stutzeri* AN0100, *P. stutzeri* AN0010, and *P. stutzeri* AN0001) that possess similar relative fitness (Figure S17; Supplementary information S3.2.1 and S3.4.1) and execute complementary metabolic reactions to degrade naphthalene by exchanging the three intermediates (Figure 5A). The limiting carbon sources of this consortium were mostly supplied by the last population, *P. stutzeri* AN0001, who converts catechol to usable pyruvate and acetyl-CoA (Figure 5A).

**Figure 5.**
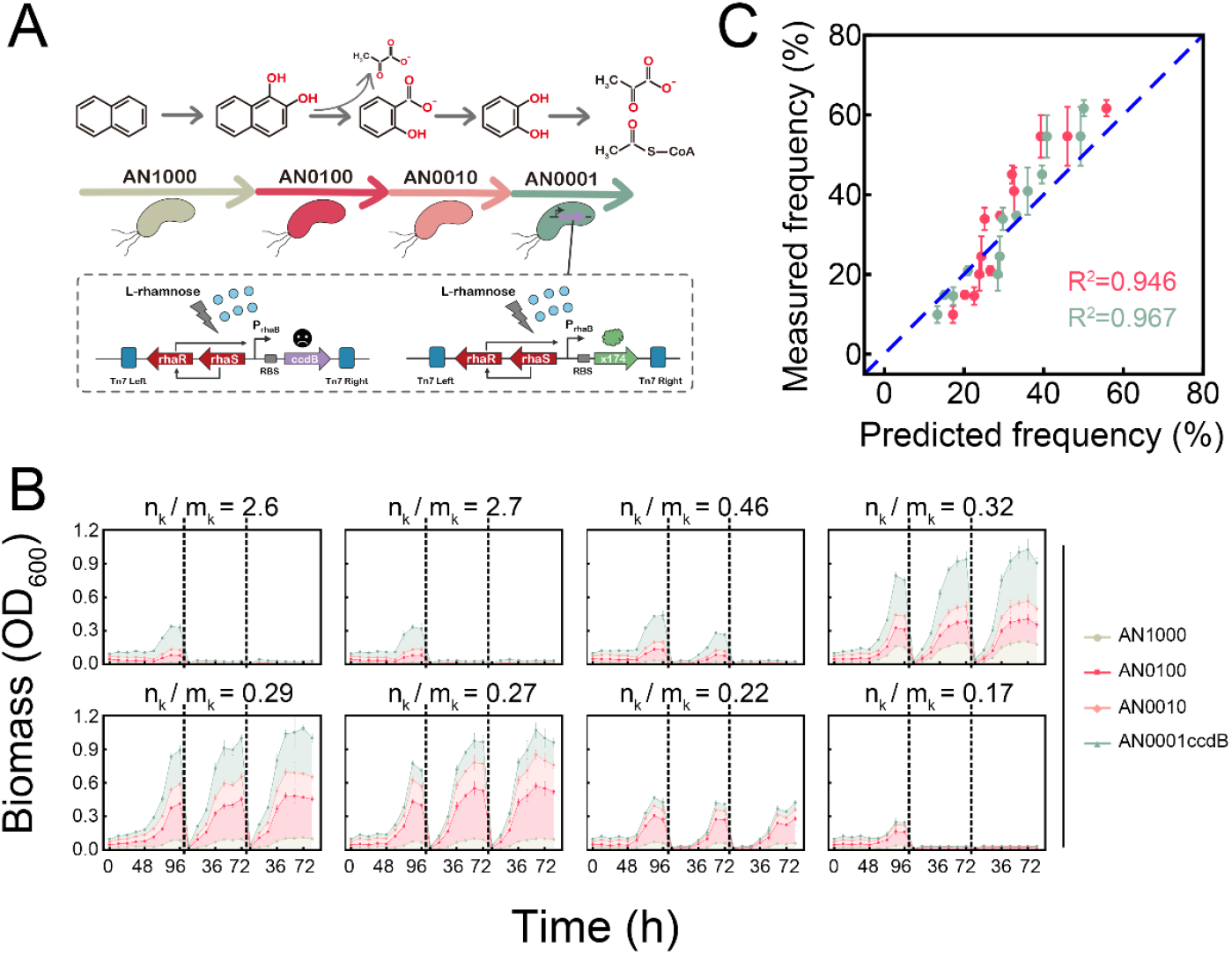
Dynamics of the four-step synthetic community. (A) Schematic diagram of the construction of the synthetic community. To experimentally modify *m*_*k*_, strain *P. stutzeri* AN0001ccdB or *P. stutzeri* AN0001×174 was used. (B) The dynamics of the synthetic community composed of *P. stutzeri* AN1000, *P. stutzeri* AN0100, *P. stutzeri* AN0010 and *P. stutzeri* AN0001ccdB from co-culture experiments under a gradient of *n*_*k*_*/m*_*k*_ value conditions (i.e., using medium with different rhamnose concentrations). The cultures were diluted by a factor of 20 after 96 h or 72h of culturing (indicated by the dashed line), and transferred into a new fresh medium. To measure their relative abundances, strain *P. stutzeri* AN1000 was labeled with ECFP, strain *P. stutzeri* AN0100 was labeled with DsRed, strain *P. stutzeri* AN0010 was labeled with mBeRFP, and strain *P. stutzeri* AN0001ccdB was labeled with EGFP. The dynamics of the community composed of *P. stutzeri* AN1000, *P. stutzeri* AN0100, *P. stutzeri* AN0010 and *P. stutzeri* AN0001×174 are shown in Figure S19A. (C) Predicting the relative abundance of the last populations (that is, *P. stutzeri* AN0001ccdB or strain *P. stutzeri* AN0001×174) by mathematical modelling. The experimental results are summarized from those stable communities shown in (B) and Figure S19A, in which the values of relative abundance at the end point of the third transfer were recorded. The red dots show the results calculated the Eqn. [3], while the green dots show the results predicted from the simulations considering the specific pathway mechanisms of naphthalene degradation (see Supplementary Information S4.2 for details).

We next cultured this consortium using naphthalene as the sole carbon source. Because the inherent growth rate of strain AN0001 is nearly identical to those of strains AN1000, AN0100, and AN0010, meaning *m*_*k*_ ≈ 0, the community showed signs of collapse (Figure S18), consistent with our basic predictions (Eqn. [5]). To generate a stable synthetic consortium, we applied a similar strategy to that used in stabilizing our two-step MDOL consortia: experimentally modifying the value of *m*_*k*_ by co-culturing a consortium containing an engineered *P. stutzeri* AN0001 (named as *P. stutzeri* AN0001*). In these experiments, we found that the community developed in a stable manner when the values of *n*_*k*_*/m*_*k*_ were present within a suitable range (Figure 5B; Figure S19A) given by Eqn. [5]. In addition, as shown in Figure 5C (Red dots), Eqn. [3] accurately predict the steady-state frequency of strain *P. stutzeri* AN0001* (R^2^= 0.946). Moreover, although the assembly of this synthetic consortium may be affected by various factors, including by-product benefit, spontaneous conversion of intermediates, as well as toxic effects of the metabolites, our numeric simulations indicate that these factors negligibly affect the prediction of the community assembly (Figure 5C; Green dots). Taken together, these results demonstrated that our theorical framework can be expanded to multi-step MDOL communities, and thus greatly contributing to a more detailed understanding of their assembly.

### Assembly influenced by initial population ratio

To test whether the structure of MDOL communities remains robust even when the initial ratio of different members is changed, we simultaneously performed mathematical simulations and experiments using our synthetic consortia, initiated using a wide range of ratios for starting strains.

For the two-step MDOL community, our simulations indicated that the rule given by Eqn. [2] still accurately predicts whether the two population stably co-exist (Figure S20 A). If the community maintains stable, we found that the steady-state ratio of the two genotypes converged to the same level that can be quantitatively predicted by Eqn. [1] (Figure S20 A and 20 B). Similar results were observed in our verification experiments (Figure S21-22). These results suggest that the assembly of two-step MDOL community is independent of the initial ratio of the two strains involved.

We then set out to test whether the assembly of multi-step MDOL community was influenced by the initial abundance of the different members involved. As shown in Figure S23-24, when one of the populations dominated the initial community, the parameter space for the stable co-existence decreased, suggesting that these extreme initial conditions considerably destabilize the community. If the community maintains stable, the frequency of one former population was significantly positively correlated with its initial frequency (Figure S25-27A), and was largely determined by the proportion of its initial abundance accounting for the total initial abundance of all the former populations (Figure S25-27B). We defined this proportion as *σ*_*k*_,

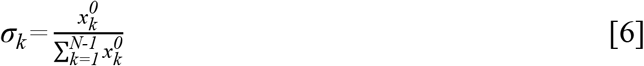

Here, 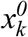 represents the initial biomass of the *k*th population (*k* = 1 ∼ *N*). Moreover, the frequency of the last population in the steady-state community remained unchanged despite changing initial conditions (Figure S23A; Figure S25-27A), but this frequency did not match well with the prediction of Eqn. [3]. We added *σ*_*k*_ as a weight to the original formula Eqn. [3], generating a novel formula,

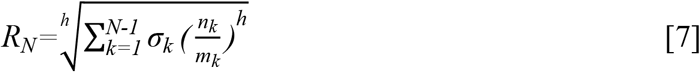

Eqn. [7] predicted the frequency of the last population more accurately than Eqn. [3] (Figure S23A; Figure S25-27C), indicating that the steady-state frequency of the last population was affected by the initial ratio of the former populations. These mathematical predictions were then verified in our synthetic consortium engaged in four-step MDOL (Figure 6B; Figure S19B; Figure S28A-B). Together, these results suggested that the initial population ratio, especially the initial ratio of the former populations, plays a critical role in governing the assembly of a multi-step MDOL community (see Supplementary Information S3.7 for further discussions).

**Figure 6.**
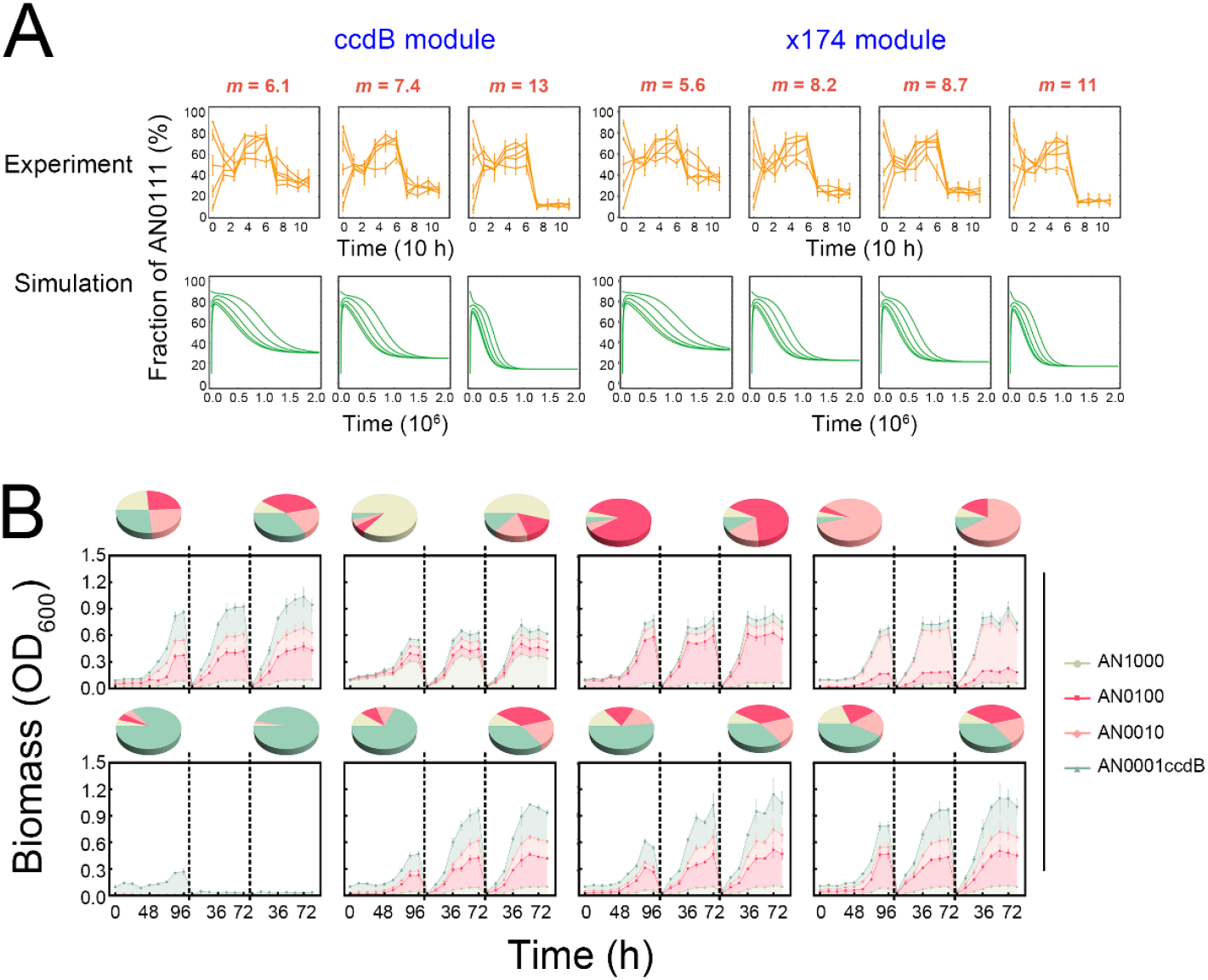
Effects of initial population ratio on the assembly rule of microbial community engaged in MDOL. (A) The assembly rule of two-step MDOL community is independent on the changing initial population ratio. Culture experiments and mathematical modelling of the community composed of *P. stutzeri* AN1000 and *P. stutzeri* AN0111ccdB (or *P. stutzeri* AN0111×174) were performed by setting seven different *m* values, as well as five different initial ratios. The relative abundances of *P. stutzeri* AN0111ccdB or *P. stutzeri* AN0111×174 from experiments (orange, first row) and mathematical modelling (green, second row) are shown. (B) Growth dynamics of the synthetic community composed of *P. stutzeri* AN1000, *P. stutzeri* AN0100, *P. stutzeri* AN0010, and *P. stutzeri* AN0001ccdB at eight different initial ratios. In these experiments, rhamnose concentration was set to 0.005% (*n*_*k*_*/m*_*k*_ value equals to 0.29). Each color region represents the relative abundance of each strain. The pie charts denote the community structure at the starting and end time points. Three independent replicates were performed for each condition. Results of same experiments using synthetic community composed of *P. stutzeri* AN1000, *P. stutzeri* AN0100, *P. stutzeri* AN0010, and *P. stutzeri* AN0001×174 are shown in Figure S19B.

## Discussion

Here, we proposed a simple rule to predict the assembly of microbial community engaged in metabolic division of labor (MDOL community) using a mathematical model. This rule was verified by experimentations using designed synthetic microbial consortia. Our rule demonstrates that the stability and assembly of MDOL community are mostly determined by how benefits are allocated among the community members. Importantly, our rule is built on the feature of most organic compound degradation pathways. One basic assumption of our rule is that the final product of a MDOL pathway is the sole carbon source for all the strains involved in the community. This feature offered a ‘private benefit’ to the last population, which represents a common challenge for developing MDOL communities. This selfish population is analogous to a human worker responsible for the final step of an assembly line, who pockets the final product without sharing the resulting profits with other workers. Therefore, we named this final population the ‘Embezzler’ population, and likened the instability of the community caused by ‘Embezzler’ as the ‘Embezzler dilemma’. This phenomenon has been observed in other MDOL communities. For instance, one study engineered a dual-species consortium for removal of the insecticide parathion, in which an *Escherichia coli* strain SD2 was responsible for hydrolyzing parathion, yielding two intermediates including *p*-nitrophenol, while another *Pseudomonas putida* strain KT2440 was responsible for metabolizing *p*-nitrophenol (Embezzler)^43^. That study found that the ‘Embezzler’ strain largely dominated the final community, which is in accordance with our observation of ‘Embezzler dilemma’. Another study investigated the interactions among five bacterial species in a cellulose-degrading community, which also found that the strains responsible for the last step of the cellulose degradation dominated the community^44^. Therefore, the ‘Embezzler dilemma’ may represent a common challenge when engineering microbial systems to remove pollutants.

Our model provides several avenues to address the issue of the ‘Embezzler dilemma’ in engineered microbial systems. Firstly, it is feasible to reduce the inherent fitness of the ‘Embezzler’ strain. This can be accomplished by either rationally engineering a slow-growing strain performing the last step, or assigning more tasks to this strain that incur higher energetic costs to the ‘Embezzler’ strain. Secondly, the ‘Embezzler dilemma’ can be also alleviated if the populations performing former steps are designed to be capable of obtaining metabolic by-products generated from the former reactions. From this perspective, when designing a synthetic consortium engaged in MDOL for pollutant degradation, it may be useful to assign a by-product to the former populations. Thirdly, our results also showed that if the substrate is toxic, the microbial system will be more stable, which is consistent with the results in our previous study^28^. In summary, our results provide a quantitative way to evaluate the feasibility of applying these strategies into a specific pathway engineering, and thus should greatly assist in designing and managing related artificial microbial systems.

Our study also has implications for our understanding on the evolution of MDOL among microorganisms. Our results suggest that the individuals of ‘Embezzler’ must evolve to possess lower relative fitness to eliminate ‘Embezzler dilemma’ and maintain system stability. From this perspective, the evolution of MDOL systems seems to contradict Darwin’s theory, which emphasizes that individuals must evolve to achieve greatest personal fitness and reproductive success^45^. This paradox suggests that evolutionary selection at the community-level^46-48^ may present a main driving force behind the evolution of MDOL communities. Our analysis presented here offers possible solutions to this paradox. As we discussed above, the presence of several specific pathway features, such as substrate toxicity and by-product production, relax the constraints ensuring the stability of a MDOL community. Therefore, MDOL among different members may be easier to evolve in the pathways that possess these specific features. This hypothesis can be examined using large-scale bioinformatic analysis, as well as the well-designed experimental evolution assays. Another solution to this paradox may be derived from the effects of spatial positioning^46^. Our previous study found that although ‘Embezzler’ strain generally grew more in a two-member microbial colony engaged in MDOL, the second strain is usually not completely excluded^28^, suggesting that specific spatial organizing of a community may facilitate their co-existence. In these spatially structured environments, interaction between the two populations would require their cells to be spatially proximal to each other^49-52^. Even ‘Embezzler’ cells can obtain asymmetric ‘private benefit’, but they must be located in close proximity to their partners. Therefore, spatial organization of a MDOL community may help to maintain the co-existence of its members, thus favoring the evolution of MDOL processes. This hypothesis can be examined by comparing the results of evolutionary experiments in biofilms and well-mixed systems.

In conclusion, our results provide a basis for a theory guiding the application of MDOL strategy to design and manipulate artificial microbial systems, and also provides new perspectives for understanding the evolution of natural MDOL systems.

## Methods

### Formulation of the ODE models

The mathematical models were built using ordinary differential equations (ODEs), which formulated the dynamics of intracellular and extracellular intermediates and end products, as well as the growth of all the populations involved in the community. In all cases, the models were built on a well-mixed system (or sufficiently fast metabolite diffusion). Here, the dimensionless forms of the models were presented. The detailed derivations of all models and justifications of our assumptions are described in Supplementary Information S1-S4. The definitions and dimensionless methods of all the variables and parameters are listed in Table S1-S5.

### The basic ODE models for two-step MDOL community

As described in the first part of the Results section, we assumed that a two-step pathway was implemented by MDOL between two populations (Figure S1A). Details about the construction of this basic ODE system are described in Supplementary Information S1.1. For simplicity, the basic model was built based on seven simple assumptions, namely transport via passive diffusion, intracellular metabolic reactions, negligible abiotic degradation of I and P, excess of initial substrate, as well as low levels of intracellular accumulation of I and P; importantly, P was assumed to be the sole and limited resource for the growth of the two populations and its consumption was calculated following Monod equations. Thus, the dynamics of intracellular and extracellular I and P are given by

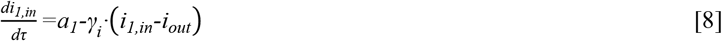

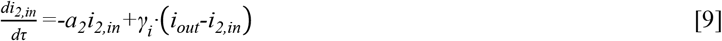

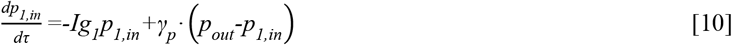

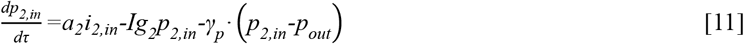

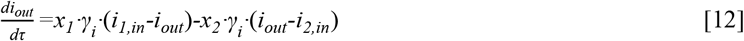

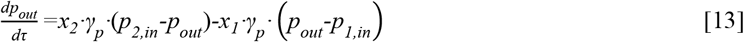

Here, *i*_*1,in*_ and *i*_*2,in*_ represent the intracellular I concentration of two populations; *i*_*out*_ is the extracellular concentration of I; *p*_*out*_ is the extracellular concentration of P; *p*_*1,in*_ and *p*_*2,in*_ are the intracellular P concentration of two populations; *x*_*1*_ and *x*_*2*_ are the biomass of two populations; *a*_*1*_ and *a*_*2*_ are the reaction rates of two reactions; *γ*_*i*_ and *γ*_*P*_ are the diffusion rates of I and P across cell membrane; *Ig*_*1*_ and *Ig*_*2*_ are the consumption rate of product for the growth of two populations. The growth of the two populations was modelled modeled using a general logistic function with first-order cell death:

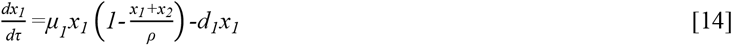

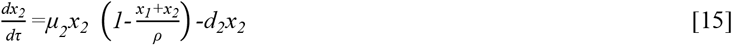

Here, *ρ* represents the carrying capacity of the whole communities; *d*_*1*_ and *d*_*2*_ represent the apparent maintenance rates of the two populations. The specific growth rates of the two populations, *μ*_*1*_ and *μ*_*2*_, are calculated by

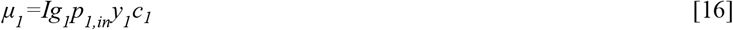

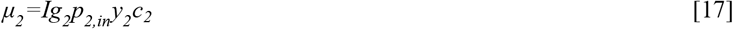

Eqn. [16] and [17] are linked with our basic assumption that P is the sole resource for growth. In addition, *y*_*1*_ and *y*_*2*_ represent the yield coefficients for biomass production of the two populations; coefficients *c*_*1*_ and *c*_*2*_ are used to describe the metabolic burdens of the two reactions.

### The ODE models that consider complex pathway mechanisms

The models that consider the complex pathway mechanisms were built by modifying or adding the related mathematical terms to the basic model. Details of these modifications are described in Supplementary Information S2.

### The ODE models for multiple-step MDOL community

Assuming that a metabolic pathway is segregated into *N* steps that are executed by *N* populations (Figure S1A), a more general models are built by expanding the basic framework of the two-step MDOL community, which was described in Supplementary Information S3 in detail. In this system, an organic substrate (S) is converted to a final product (P) through a long metabolic pathway containing *N* reactions and *N*-1 intermediate metabolites (I). Each reaction is carried out by one population by expressing a specific enzyme. The ODE model was built accordingly,

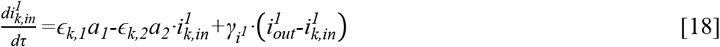

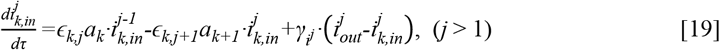

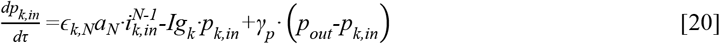

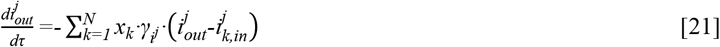

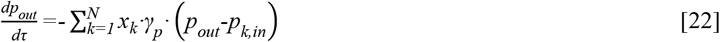

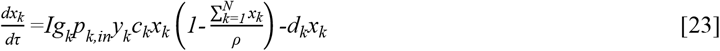

In the model, 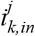 represents the intracellular concentration of the *j*th intermediate of the *k*th population; 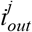 represents the extracellular concentration of the *j*th intermediate; *p*_*k,in*_ represents the intracellular P concentration of the *k*th population; *ε*_*k*_ represents a vector that characterizes the phenotype of the *k*th population, where *ε*_*k*_ equals [0,0,…,1,…0,0] denoting that the *k*th population is only capable of performing the *k*th reaction; *x*_*k*_ is biomass of the *k*th population; *a*_*k*_ is the reaction rates of the *k*th reactions; 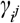 and *γ*_*P*_ are the diffusion rates of the *j*th intermediate and P across cell membrane; *Ig*_*k*_ are the consumption rate of P of the *k*th population; *ρ* is the carrying capacity of the whole communities; *d*_*k*_ is apparent maintenance rate of the *k*th population; *y*_*k*_ is the yield coefficients for biomass production of the *k*th population; *c*_*k*_ is the metabolic burdens of the *k*th population.

### The ODE model used to predict the assembly of our synthetic consortia

The models used to predict the assembly of the three 2-step MDOL synthetic consortia, as well as the four-step MDOL synthetic consortium were built by adding the specific mechanisms of each consortium to our basic ODE model. These effects include the toxicity of the naphthalene and the intermediates, the abiotic conversion of the intermediates, the generation of metabolic by-products in the step of converting 1, 2-hydroxynaphthalene to salicylate. In addition, the parameters, such as the reaction rates, the consuming rate of P, were experimentally measured or obtained from previous reports. Specifically, the value of *d*_*2*_ (2-step consortia) and *d*_*4*_ (four-step consortia) are estimated by the experimentally fitted function that links death rate to the rhamnose concentration (see Supplementary Information S4.1.4 for details of the measurements). Details of formulation of these predicting models are described in Supplementary Information S4.2, and the values and sources of all the parameters used in the predicting models are listed in Table S7.

### Model derivation and simulation protocols

Solving and simplifying of the ODE models were performed using the Solve, Dsolve and Simplify functions of *Wolfram Mathematica* (version 12.0), associated with manual arrangement. In order to perform sensitive analyses of the basic model, as well as solve those ODE systems that cannot be easily managed using simple analytic method, numeric simulations were performed using basic settings of NDsolve function of *Wolfram Mathematica*. In particular, derivation methods of the basic assembly rule of 2-step MDOL community (see Eqns. [1] and [2]) are described in Supplementary Information S1.3; Analyses of the models considering those complex mechanisms are described in Supplementary Information S2; Derivation methods of the assembly rule of multiple-step MDOL community (that is, Eqns. [3]-[5]) are described in Supplementary Information S3.2 and S3.4; Simulation protocols and parameterization of the models for predicting experimental results are described in Supplementary Information S4.2; Simulation protocols that test the effects of initial population ratio on the assembly rule are described in Supplementary Information S1.5 and S3.6. These analyses were performed using custom *Wolfram Mathematica* scripts. The generated data were then analyzed and visualized using basic functions in *Wolfram Mathematica*, of which the custom codes were integrated to the aforementioned scripts. Specifically, to fit simulation data with the proposed function, NonlinearModelFit function were used with the default settings.

The source codes used for all the models concerning two-step MDOL community are available on: https://github.com/RoyWang1991/MDOLcode/tree/master/MDOL-LVMM-twomember, while the codes for the models concerning multi-step MDOL community are available on https://github.com/RoyWang1991/MDOLcode/tree/master/MDOL-LVMM-mutiple.

### Construction and culturing of the synthetic microbial communities

#### Construction of the strains involved in the synthetic microbial communities

The strains and plasmids used in this study are summarized in Table S6. All *P. stutzeri* strains were engineered from a naphthalene-degrading bacterial strain *P. stutzeri* AN10^33,40,53^. Genes that encode the key enzymes responsible for the four metabolic steps in naphthalene degradation pathway were knocked out to generate different strains involved in the synthetic microbial consortia. The details of the strain design and construction are described in Supplementary information S4.1. To label the strains with fluorescence for the measurement of their relative abundance in synthetic consortia, *eCFP, dsRed, mBeRFP, mCherry, eGFP* genes were cloned into a constitutive vector, pMMPc-Gm^54^, using the Hieff Clone® Plus One Step Cloning Kit (Yeasen, Shanghai, China), and delivered to the host cells via triparental filter mating^55^.

#### Culturing of the synthetic microbial communities

Our synthetic microbial communities were cultured in 25-mL flask containing 5 mL new fresh minimum medium^56^ supplemented with naphthalene powder (1% w/v) as the sole carbon source. The biomass and relative fraction were measured using the method described previously^28,57^. The detailed protocols were described in Supplementary information S4.1.6.

## Acknowledgments

We wish to thank Professor Ping Xu (Shanghai Jiao Tong University, Shanghai, P.R. China) for supplying plasmid pMMPc-Gm, used for fluorescence labeling in this study; Dr. Min Lin (Chinese Academy of Agricultural Sciences, Beijing, P.R. China) for providing plasmid pK18mobsacB and pRK2013, used for genetic engineering in this work; Dr. T. Juelich (UCAS, Beijing) for linguistic assistance during the preparation of this manuscript.

This work was supported by National Key R&D Program of China (2018YFA0902100 and 2018YFA0902103), and National Natural Science Foundation of China (91951204, 31761133006, 31770120, and 31770118).

