## Supplementary Information for "Even allocation of benefits stabilizes microbial community engaged in metabolic division of labor"

### S1 Derivation of the basic model regarding the assembly of a two-step MDOL community

#### S1.1 Derivation of the basic model

As described in the main text, the basic model was constructed to describe the community dynamics composed of two populations, [1, 0] and [0, 1], in which [1, 0] degrades a substrate (S) into an intermediate (I), and then to the end product (P) by [0, 1]. We first made seven biologically relevant assumptions to simplify model development and analysis. These assumptions include:

(1) The systems for both configurations are well mixed in each compartment (inside a cell or in the extracellular space). (the effects of limiting mass diffusion are discussed in our previous study[1]).

(2) Transport of the metabolites, including substrate (S), intermediates (I) and end product (P), across the cell membrane occurs via passive diffusion at a rate proportional to the concentration gradient between two compartments. (Scenarios considering active transport of metabolites are discussed in Supplementary Information 2.1).

(3) Metabolic reactions were performed intracellularly. (Scenarios considering extracellularly catalyzed metabolic reactions are discussed in Supplementary Information 2.2).

(4) The spontaneous degradation of I and P is significantly slower than kinetic reactions and thus its rate may be set to approximately 0. (Scenarios considering considerable fast spontaneous degradation of I and P are discussed in Supplementary Information

2.3).

(5) The supply of S in the system is sufficient, and thus the rate of S consumption remains constant. (Scenarios considering limited supply of S are discussed in Supplementary Information 2.4).

(6) The intracellular accumulation of I and P in the system is negligible, and thus the rates of the two metabolic reactions follow the first-order kinetics. (Scenarios relaxing this assumption are discussed in Supplementary Information 2.5-2.6).

(7) The end product of pathway is the sole carbon source that all populations use for growth. (Scenarios considering another carbon source, the by-product generated from the first reaction, are discussed in Supplementary Information 2.7).

*S1.1.1 Equations regarding Intermediate and product dynamics*

We first built a system of ODEs describing intermediate and product concentrations for an intracellular pathway in a two-step MDOL community.

$$\frac{dI_{1,in}}{dt}\cdot V_c=\frac{k_1E_1}{K_1+S_{1,in}}S_{1,in}-r_I\cdot V_c\cdot\left(I_{1,in}-I_{out}\right) \quad [S1.1]$$

$$\frac{dI_{2,in}}{dt}\cdot V_c=-\frac{k_2E_2}{K_2+I_{2,in}}I_{2,in}+r_I\cdot V_c\cdot\left(I_{out}-I_{2,in}\right) \quad [S1.2]$$

$$\frac{dP_{1,in}}{dt}\cdot V_c=-\frac{kg_1}{Kg_1+P_{1,in}}P_{1,in}+r_P\cdot V_c\cdot\left(P_{out}-P_{1,in}\right) \quad [S1.3]$$

$$\frac{dP_{2,in}}{dt}\cdot V_c=\frac{k_2E_2}{K_2+I_{2,in}}I_{2,in}-\frac{kg_2}{Kg_2+P_{2,in}}P_{2,in}+r_P\cdot V_c\cdot\left(P_{out}-P_{2,in}\right) \quad [S1.4]$$

$$\frac{dI_{out}}{dt}=X_1\cdot r_I\cdot V_c\cdot\left(I_{1,in}-I_{out}\right)-X_2\cdot r_I\cdot V_c\cdot\left(I_{out}-I_{2,in}\right) \quad [S1.5]$$

$$\frac{dP_{out}}{dt}=X_2\cdot r_P\cdot V_c\cdot\left(P_{2,in}-P_{out}\right)-X_1\cdot r_P\cdot V_c\cdot\left(P_{out}-P_{1,in}\right) \quad [S1.6]$$

According to assumption (4) and (5), We further assumed that $S_{1,in}\gg K_1, K_2\gg I_{2,in}$,

$K_{g1} \gg P_{1,in}$, $K_{g2} \gg P_{2,in}$, $Ig_1=\frac{kg1}{K_{g1}}$, $Ig_2=\frac{kg2}{K_{g2}}$, which simplified the model to a linear system,

$$\frac{dI_{1,in}}{dt}\cdot V_c=k_1E_1-r_I\cdot V_c\cdot\left(I_{1,in}-I_{out}\right) \quad [S1.7]$$

$$\frac{dI_{2,in}}{dt}\cdot V_c=-\frac{k_2E_2}{K_2}I_{2,in}+r_I\cdot V_c\cdot\left(I_{out}-I_{2,in}\right) \quad [S1.8]$$

$$\frac{dP_{1,in}}{dt}\cdot V_c=-\frac{kg_1}{Kg_1}P_{1,in}+r_P\cdot V_c\cdot\left(P_{out}-P_{1,in}\right) \quad [S1.9]$$

$$\frac{dP_{2,in}}{dt}\cdot V_c=\frac{k_2E_2}{K_2}I_{2,in}-\frac{kg_2}{Kg_2}P_{2,in}+r_P\cdot V_c\cdot\left(P_{out}-P_{2,in}\right) \quad [S1.10]$$

$$\frac{dI_{out}}{dt}\cdot V_c=X_1\cdot r_I\cdot V_c\cdot\left(I_{1,in}-I_{out}\right)-X_2\cdot r_I\cdot V_c\cdot\left(I_{out}-I_{2,in}\right) \quad [S1.11]$$

$$\frac{dP_{out}}{dt}\cdot V_c=X_2\cdot r_P\cdot V_c\cdot\left(P_{2,in}-P_{out}\right)-X_1\cdot r_P\cdot V_c\cdot\left(P_{out}-P_{1,in}\right) \quad [S1.12]$$

To facilitate modeling analysis, we then non-dimensionalized these ODEs following the method described in Table S1 and Table S2, which formalized the model (Eqns. 8-13) in the Methods section of the main text.

*S1.1.2 Equations regarding population growth dynamics*

Our model then accounts for the growth dynamics of the two populations in a two-step MDOL community, which followed a general logistic function with first-order cell death

$$\frac{dX_1}{dt}=g_1X_1\left(1-\frac{X_1+X_2}{N_m}\right)-D_1X_1 \quad [S1.13]$$

$$\frac{dX_2}{dt}=g_2X_2\left(1-\frac{X_1+X_2}{N_m}\right)-D_2X_2 \quad [S1.14]$$

As described in assumption (6), we assume that all populations only use the end product as the sole carbon source, thus the growth rate of the two populations, $g_1$ and $g_2$, are calculated via a Monod equation to the intracellular concentration of the end product, so that,

$$g_1 = \frac{kg_1 P_{1,in}}{Kg_1 + P_{1,in}} Y_1 c_1 \quad \text{[S1.15]}$$

$$g_2 = \frac{kg_2 P_{2,in}}{Kg_2 + P_{2,in}} Y_2 c_2 \quad \text{[S1.16]}$$

We then assumed that $K_{g1} \gg P_{1,in}$ , $K_{g2} \gg P_{2,in}$ as mentioned above, and non-dimensionalized the two equations following the method described in Table S1 and Table S2, so obtain the dimensionless cell growth equations (Eqns. 14-17 in the Methods section of the main text.).

#### S1.2 Definitions and values of dimensionless variables and parameters

The definition of all the variables, their corresponding dimensionless counterparts, and their value range during the numerical simulations are listed in Table S1. In addition, the definition of all the parameters, their corresponding dimensionless methods (when applicable), and choice of their values (from previous studies) for numerical simulations are listed in Table S2. Parameter values are chosen/calculated according to estimated values from literature and are listed in Table S2. Dimensionless (ND) values are calculated based on methods in Table S2. Parameter ranges are chosen to incorporate at least ~10-fold higher or lower than the estimated value. When performing mathematical simulations, the initial values of the biomass of the two populations was set to $x_{i,0}$ (values listed in Table S2), while the initial values of other variables were set to zero.

#### S1.3 Derivation of the basic assembly rule of a two-step MDOL community

To derive the simple formula that defines the basic assembly rule of a two-step MDOL community, we solved the steady-state equations of basic model system (Eqns. [8]-[17] in Methods section of the main text), using Solve function of *Wolfram Mathematica*. The solving results only contains one set of non-zero algebraic solutions. We first focused on the solution of the steady-state biomass of the two populations, $x_1$ and $x_2$, which are given by

$$x_1=\rho\cdot\frac{\frac{c_1Ig_1y_1d_2}{c_2Ig_2y_2d_1}\left(-d_1\left(\frac{Ig_2}{\gamma_p}+1\right)+a_1c_1\gamma_py_1\right)+\left(\frac{Ig_1}{\gamma_p}+1\right)(d_1-a_1c_1y_1)y_2}{a_1c_1y_1\left(\frac{c_1Ig_1y_1d_2}{c_2Ig_2y_2d_1}-1\right)} \quad [S1.17]$$

$$x_2=\rho\cdot\frac{Ig_1}{\gamma_p}\cdot\frac{\frac{c_1Ig_1y_1d_2}{c_2Ig_2y_2d_1}\left(-d_1\left(\frac{Ig_2}{\gamma_p}+1\right)+a_1c_1\gamma_py_1\right)+\left(\frac{Ig_1}{\gamma_p}+1\right)(d_1-a_1c_1y_1)y_2}{a_1c_1y_1\left(\frac{c_1Ig_1y_1d_2}{c_2Ig_2y_2d_1}-1\right)\left(\frac{c_1Ig_1y_1d_2}{c_2Ig_2y_2d_1}-\left(\frac{Ig_1}{\gamma_p}+1\right)\right)} \quad [S1.18]$$

Then we formalized the steady-state relative abundance of [0, 1], as below:

$$R_{[0,1]}=\frac{x_1}{x_1+x_2}=\frac{Ig_1}{\gamma_p}\cdot\frac{\frac{c_2Ig_2y_2}{d_2}}{\left(\frac{c_1Ig_1y_1}{d_1}-\frac{c_2Ig_2y_2}{d_2}\right)} \quad [S1.19]$$

Eqn. [S1.19] can be simplified by defining $n=Ig_1/\gamma_p$ and $m=(\frac{c_1Ig_1y_1}{d_1}-\frac{c_2Ig_2y_2}{d_2})/\frac{c_2Ig_2y_2}{d_2}$, thus we obtain Eqn. [1] in the main text. Here, $Ig_1$ is the consuming rate of P for [1,0], while $\gamma_p$ is the diffusive coefficient of P. Since the first population [1, 0] absorbs P from the extracellular environment via passive diffusion, the value of $\gamma_p$ basically determines the supply rate of P for [1, 0]. Therefore, $n$ equals the ratio of $Ig_1$ to $\gamma_p$, reflecting whether the supply of P (produced and secreted by [0, 1]) is sufficient for the growth of [1, 0] (the 'Product demand gap' for [1, 0]), in which higher $n$ suggests lower 'Product demand gap'. Meanwhile, lower 'Product demand gap' of [1, 0] means that [0, 1] privatized more P and share less to [1, 0], thus $n$ also indicates the 'private benefit' of [0, 1] derived from the product privatization. Furthermore, $\frac{c_1Ig_1y_1}{d_1}$ and $\frac{c_2Ig_2y_2}{d_2}$ reflect the inherent net growth rate of [1, 0] and [0, 1], respectively, thus $m$ indicates the difference between the inherent growth rates of the two populations normalized by the inherent growth rate of [0, 1]. Therefore, Eqn. [1] indicates that the relative abundance of the [0, 1] in the final stable MDOL community equals the quotient of the 'private benefit' of [0, 1] and the growth difference between the two populations.

**S1.4 Sensitive analyses of the basic assembly rule of a two-step MDOL community**

For the sensitive analyses of the proposed rule, we performed numeric simulations to solve the ODE system of Eqns. 8-17 in Methods section of the main text. We tested the effects of the parameters that were not included in Eqn. [1], that is, the rates of the first ($a_1$) and the second reaction ($a_2$), maximum biomass capacity ($\rho$), as well as the diffusive coefficient of I ($\gamma_i$). For the test of each parameter, 10000 parameter sets containing an even 10×10 gradient of $n$ and $m$ were first generated. Then the values of the tested parameter were varied, and numerous simulations were performed with each setting, using NDsolve function of *Wolfram Mathematica*. Other parameters were set as the default value shown in Table S1.

As shown in Figure S1, While the change of $a_2$, $\rho$, and $\gamma_i$ rarely changed the predictions of our rule (Figure S1C-H), the rule was significantly affected by the change of the rate of first reaction ($a_1$), in which the lower $a_1$ would shrink the parameter space that defines the stability of a MDOL community (Figure S1A-B).

**S1.5 Testing the effects of initial strain ratio on the assembly rule of a two-step MDOL community**

In order to test whether the initial strain ratio affects the basic assembly rule of a two-step MDOL community, we still first generated the 10000 parameter sets with the 10×10 gradient of $n$ and $m$, as described above. Next, the initial total biomass ($x_1$+$x_2$) was set as 0.0002, and seven different initial strain ratios were designed for the two populations: 1:9, 1:7, 2:1, 3:2, 7:1, 9:1. In addition, three different reaction rates of the first step ($a_1$) were also included. In summary, a total of 212100 parameter sets were designed for subsequent numeric simulations based on our basic ODE system. The values of other parameters were set as the default shown in Table S2.

### S2 Generalizing the rule considering more complex pathway mechanisms

#### S2.1 Different mechanisms for the transport of the intermediate and product

Transport of metabolites, I and P, across the cell membrane can be carried out by passive diffusion or active transport. In the latter scenarios, I and P are possibly transported against the concentration gradient mediated by active transporter proteins. In a two-step MDOL community, transport of I and P includes four main processes (Figure S2A): (1) I being exported out of [1, 0] cells; (2) I being imported by imported into [0, 1] cells; (3) P being imported by imported into [1, 0] cells; (4) P being exported out of [0, 1] cells. While each process can be executed by passive diffusion or active transport, sixteen configurations are possibly present in the kinetics of a MDOL community (Figure S2C). We thus discussed the assembly rules of the community in all these scenarios. To formulize passive diffusion, the rate of metabolite transport was assumed to be proportional to the concentration gradient between two compartments; To formulize active transport, the rate of metabolite transport was assumed to be proportional to the concentration of the corresponding metabolite (Figure S2B). For example, in the scenario where all the four processes are fulfilled by active transport, the system of dimensionless ODEs corresponding to intermediate and product dynamics is given by:

$$\frac{di_{1,in}}{d\tau}=a_1-\gamma_{i1}^{a}\cdot i_{1,in} \qquad [S2.1]$$

$$\frac{di_{2,in}}{d\tau}=-a_2 i_{2,in}+\gamma_{i2}^{a}\cdot i_{out} \qquad [S2.2]$$

$$\frac{dp_{1,in}}{d\tau}=-Ig_1 p_{1,in}+\gamma_{p1}^{a}\cdot p_{out} \qquad [S2.3]$$

$$\frac{dp_{2,in}}{d\tau}=a_2 i_{2,in}-Ig_2 p_{2,in}+\gamma_{p2}^{a}\cdot p_{2,in} \quad [S2.4]$$

$$\frac{di_{out}}{d\tau}=x_1\cdot\gamma_{i1}^{a}\cdot i_{1,in}-x_2\cdot\gamma_{i2}^{a}\cdot i_{out} \quad [S2.5]$$

$$\frac{dp_{out}}{d\tau}=x_2\cdot\gamma_{p2}^{a}\cdot p_{2,in}-x_1\cdot\gamma_{p1}^{a}\cdot p_{out} \quad [S2.6]$$

As shown in (Figure S2C), the assembly rules in these scenarios can be clustered into three groups, as follows.

*S2.1.1 When the processes (3) and (4) are mediated by passive diffusion*

When P are transported by passive diffusion (the processes (3) and (4) are mediated by passive diffusion), no matter how I was transported (process (1) and (2)), the ODE systems can still be simply solved by the same analytic method as we used the basic model. The resulting formula is very similar as the basic form

$$R_{[0,1]}=\frac{x_1}{x_1+x_2}=\frac{Ig_1}{\gamma_{p1}}\cdot\frac{\frac{c_2 Ig_2 y_2}{d_2}}{(\frac{c_1 Ig_1 y_1}{d_1}-\frac{c_2 Ig_2 y_2}{d_2})} \quad [S2.7]$$

Eqn. [S2.7] can be then simplified by defining $n'=Ig_1/\gamma_{p1}$ and $m=(\frac{c_1 Ig_1 y_1}{d_1}-\frac{c_2 Ig_2 y_2}{d_2})/\frac{c_2 Ig_2 y_2}{d_2}$, thus

$$R_{[0,1]}=\frac{n'}{m} \quad [S2.8]$$

Eqn. [S2.7] is very same as Eqn. [1], except that here 'Private benefit' ($n'$) is only determined by the diffusive coefficient that governs the transport of P into [1, 0] cells ($\gamma_{p1}$). The conditions defining when the two populations can stable co-exist also rarely changed:

$$0 < n' < m \quad [S2.9]$$

Therefore, in the first group of scenarios, the assembly rule remains largely unchanged.

*S2.1.2 When the processes (3) is mediated by active transport, while (4) is by passive diffusion*

When P is exported out of [0, 1] cells by passive diffusion, but [1, 0] can actively transport P into the cells against the concentration gradient, the assembly rule of community changed. The formula derived from the same analytic method is given by

$$R_{[0,1]}=\frac{x_1}{x_1+x_2}=\frac{Ig_1}{\gamma_{p1}}\cdot\frac{\frac{c_2Ig_2y_2}{d_2}}{\frac{c_1Ig_1y_1}{d_1}} \quad [S2.10]$$

Assuming that $n'=Ig_1/\gamma_{p1}$ and $m=(\frac{c_1Ig_1y_1}{d_1}-\frac{c_2Ig_2y_2}{d_2})/\frac{c_2Ig_2y_2}{d_2}$, [S2.10] is simplified as

$$R_{[0,1]}=\frac{n'}{m+1} \quad [S2.11]$$

the conditions defining when the two populations can stable co-exist turned into

$$0<n'<m+1 \quad [S2.12]$$

Compared with the basic condition defined by Eqn. [2], Eqn. [S2.12] suggests that [1, 0] and [0, 1] becomes easier to stay co-exist. In addition, compared with the basic rule defined by Eqn. [1], Eqn. [S2.11] indicates that the relative abundance of [1, 0] in the steady-state community increases. These shifts are because in these cases, [1, 0] can actively absorbed P from environments, even against the concentration gradient, which help [1, 0] better acquire the limited P from environment. However, cells executing active transport commonly requires additional energy investment to synthesize the transporter protein, as well as the associated appendages. This investment increases the metabolic burden of [1, 0] (means $c_1$ should decrease), resulted in lower value of *m*. Therefore, whether active transport of P favors the growth of [1, 0], as well as the stability of a MDOL community, actually depends on the trade-off between the energy cost of the transport and its benefit from active absorption of P, which is related to specific metabolic pathway.

*S2.1.3 When the processes (4) is mediated by active transport*

When P is exported out of [0, 1] cells by active transport, no matter what other processes is mediated by, the assembly rule of community changed a lot. Derived from the same analytic method, the formula defining relative abundance of the [0, 1] ($R_{[0,1]}$) at steady-state becomes

$$R_{[0,1]}=\frac{x_1}{x_1+x_2}=\frac{Ig_1}{\gamma_{p1}}\cdot\left(\frac{\frac{c_2Ig_2y_2}{d_2}+\frac{Ig_1}{\gamma_{p1}}}{\frac{c_1Ig_1y_1}{d_1}}\right) \quad [S2.13]$$

Assuming that $n'=Ig_1/\gamma_{p1}$ and $m=(\frac{c_1Ig_1y_1}{d_1}-\frac{c_2Ig_2y_2}{d_2})/\frac{c_2Ig_2y_2}{d_2}$, Eqn. [S2.13] is simplified

$$R_{[0,1]}=\frac{n'}{n'+m+1} \quad [S2.14]$$

Since

$$0<\frac{n'}{n'+m+1}<1 \quad [S2.15]$$

we obtained

$$\gamma_{p2}^{a}<Ig_2 \quad [S2.16]$$

These expressions indicate that a MDOL community becomes much easier to be stable compared with the basic conditions when the processes (4) is mediated by active transport. However, it should be noted that actively exporting P out of the cells is an altruistic behavior, because [0, 1] unselfishly share its own resource to benefit the growth of [1, 0]. Without additional maintenance mechanisms (such as kin-selection), this kind of altruistic behavior is thought to be evolutionary unfavored[2,3].

To exclude the effect of such altruistic behavior, we added another assumption that the intracellular P concentration of [0, 1] must be greater than its extracellular concentration ($p_{2,in} > p_{out}$) in the steady-state, which ensure that [0, 1] cannot provide the 'public goods', P, for other populations in totally altruistic manners. When the population [1, 0] obtains P by passive diffusion, by solving the ODE system, the intracellular and extracellular P concentrations at the steady-state is given by

$$p_{2,in} = \frac{\frac{c_1 Ig_1 y_1}{d_1}}{\frac{c_2 Ig_2 y_2}{d_2}} \cdot \frac{1}{(\frac{Ig_1}{\gamma_{p1}}+1)} \cdot \frac{a_1 \gamma_{i2}}{\gamma_{i1}(Ig_2 - \gamma_{p2})} \quad [S2.17]$$

$$p_{out} = \frac{a_1 \gamma_{i2}}{\gamma_{i1}(Ig_2 - \gamma_{p2})} \quad [S2.18]$$

Following the assumption $p_{2,in} > p_{out}$, and Assuming that $n' = Ig_1/\gamma_{p1}$ and $m = (\frac{c_1 Ig_1 y_1}{d_1} - \frac{c_2 Ig_2 y_2}{d_2})/\frac{c_2 Ig_2 y_2}{d_2}$, we obtained an expression totally same as Eqn. [S2.9]

$$0 < n' < m \quad [S2.19]$$

Applying the same method, the condition when the population [1, 0] obtains P by active transport was also obtained, which is same as Eqn. [S2.12]

$$0 < n' < m+1 \quad [S2.20]$$

Therefore, when the altruistic behavior of [0, 1] is restrained to a certain degree (as defined by the additional assumption $p_{2,in} > p_{out}$), the condition that defines whether a MDOL community can be stable still follows our basic rule.

#### S2.2 Scenarios when metabolic reactions are performed extracellularly

Our basic model simply assumed that both metabolic reactions were performed in the intracellular space (Figure 1A). In natural systems, microorganisms are also capable of secreting enzymes to outside of cells, thus the metabolic reactions can be catalyzed extracellularly. We thus modified our basic model to formulate the assembly rule of a

MDOL community where alternative or both metabolic reactions were carried out extracellularly.

*S2.2.1 When the first metabolic reaction is performed extracellularly and the second is performed intracellularly*

In this scenario, strain [1, 0] secretes E1 to the extracellular environment and catalyzes the conversion of S to I. I was then transported into the cells of [0, 1], before it being transformed to P (Figure S3A; the first diagram). The dimensionless ODE systems considering the dynamics of intracellular and extracellular I and P are given as follow:

$$\frac{di_{1,in}}{d\tau}=-\gamma_i\cdot\left(i_{1,in}-i_{out}\right) \qquad [S2.21]$$

$$\frac{di_{2,in}}{d\tau}=-a_2 i_{2,in}+\gamma_i\cdot\left(i_{out}-i_{2,in}\right) \qquad [S2.22]$$

$$\frac{dp_{1,in}}{d\tau}=\gamma_p\cdot\left(p_{out}-p_{1,in}\right)-Ig_1 p_{1,in} \qquad [S2.23]$$

$$\frac{dp_{2,in}}{d\tau}=a_2 i_{2,in}-Ig_2 p_{2,in}-\gamma_p\cdot\left(p_{2,in}-p_{out}\right) \qquad [S2.24]$$

$$\frac{di_{out}}{d\tau}=a_1+x_1\cdot\gamma_i\cdot\left(i_{1,in}-i_{out}\right)-x_2\cdot\gamma_i\cdot\left(i_{out}-i_{2,in}\right) \qquad [S2.25]$$

$$\frac{dp_{out}}{d\tau}=x_2\cdot\gamma_p\cdot\left(p_{2,in}-p_{out}\right)-x_1\cdot\gamma_p\cdot\left(p_{out}-p_{1,in}\right) \qquad [S2.26]$$

The ODEs regarding the growth of the two populations remain unchanged as defined by Eqn. [14]-[17]. The entire modified ODE system can be simply solved by the same analytic method as the basic model, in which we obtained totally same assembly rule as defined by Eqn. [1] and Eqn. [2]. Therefore, if only the first metabolic reaction is performed extracellularly, the assembly rule remains unchanged.

*S2.2.2 When the second metabolic reaction is performed extracellularly and the first is performed intracellularly*

In this scenario, S was first transported into cells of [1, 0] and then converted to I intracellularly. After I was exported to the extracellular space, it was directly transformed to P, catalyzed by enzyme E2 secreted by strain [0, 1] (Figure S3A; the second diagram). The dimensionless ODE systems of this scenario are given by:

$$\frac{di_{1,in}}{d\tau}=a_1-\gamma_i\cdot\left(i_{1,in}-i_{out}\right) \quad [S2.27]$$

$$\frac{di_{2,in}}{d\tau}=\gamma_i\cdot\left(i_{out}-i_{2,in}\right) \quad [S2.28]$$

$$\frac{dp_{1,in}}{d\tau}=-Ig_1p_{1,in}+\gamma_p\cdot\left(p_{out}-p_{1,in}\right) \quad [S2.29]$$

$$\frac{dp_{2,in}}{d\tau}=-Ig_2p_{2,in}-\gamma_p\cdot\left(p_{2,in}-p_{out}\right) \quad [S2.30]$$

$$\frac{di_{out}}{d\tau}=-a_2i_{out}+x_1\cdot\gamma_i\cdot\left(i_{1,in}-i_{out}\right)-x_2\cdot\gamma_i\cdot\left(i_{out}-i_{2,in}\right) \quad [S2.31]$$

$$\frac{dp_{out}}{d\tau}=a_2i_{out}+x_2\cdot\gamma_p\cdot\left(p_{2,in}-p_{out}\right)-x_1\cdot\gamma_p\cdot\left(p_{out}-p_{1,in}\right) \quad [S2.32]$$

The ODEs regarding the growth of the two populations remain unchanged as defined by Eqn. [14]-[17]. However, the steady-state equations of this ODE system have an infinite number of solutions, which may be affected by the initial conditions (see below). We obtained two equations regarding the steady-state biomass of the two populations through mathematical analysis

$$\frac{a_1c_1Ig_1y_1\gamma_p(Ig_2+\gamma_p)x_1\left(1-\frac{x_1+x_2}{\rho}\right)}{\gamma_i(\gamma_p\left(Ig_1x_1+Ig_2x_2\right)+Ig_1Ig_2(x_1+x_2))}=d_1 \quad [S2.33]$$

$$\frac{a_1c_2Ig_2y_2\gamma_p(Ig_1+\gamma_p)x_1\left(1-\frac{x_1+x_2}{\rho}\right)}{\gamma_i(\gamma_p\left(Ig_1x_1+Ig_2x_2\right)+Ig_1Ig_2(x_1+x_2))}=d_2 \quad [S2.34]$$

Deriving from these two equations, we obtained a formula that defines a prerequisite for the co-existence of two populations

$$\frac{c_1 Ig_1 y_1}{(Ig_1+\gamma_p)d_1} = \frac{c_2 Ig_2 y_2}{(Ig_2+\gamma_p)d_2} \qquad [S2.35]$$

This equation indicates that the co-existence of two populations in this scenario requires strain [1, 0] and [0, 1] possess same inherent fitness.

To further test this rule, we searched 14641 parameter sets. In these parameter sets, only 41 parameter sets satisfy the condition defined by Eqn. [S2.35]. We performed simulations initialized with these parameter sets, as well as included nine different initial strain ratios. The results suggested that co-existence of three populations (that is, exhibiting stable community dynamics) is only present in the 41 parameter conditions (Figure S3B; left) that meets the requirement of Eqn. [S2.35]. This result confirms that the Eqn. [S2.35] is the necessary condition for the stability of a two-step MDOL community in this scenario. In addition, we also found that the community structures in steady-state of these simulations are nearly same as the initial community structures (Figure S3B; right), suggesting the assembly of a MDOL community in this scenario relies on initial conditions.

*S2.2.3 When both metabolic reactions are performed extracellularly*

In this scenario, conversion of S to I was catalyzed by E1 from strain [1, 0], and I was also transformed extracellularly to P by E2 from strain [0, 1] (Figure S3A; the third diagram). The dimensionless ODE systems considering the dynamics of intracellular and extracellular I and P are thus formulated by:

$$\frac{di_{1,in}}{d\tau} = -\gamma_i \cdot \left(i_{1,in} - i_{out}\right) \qquad [S2.36]$$

$$\frac{di_{2,in}}{d\tau}=\gamma_i\cdot\left(i_{out}-i_{2,in}\right) \quad [S2.37]$$

$$\frac{dp_{1,in}}{d\tau}=-Ig_1p_{1,in}+\gamma_p\cdot\left(p_{out}-p_{1,in}\right) \quad [S2.38]$$

$$\frac{dp_{2,in}}{d\tau}=-Ig_2p_{2,in}-\gamma_p\cdot\left(p_{2,in}-p_{out}\right) \quad [S2.39]$$

$$\frac{di_{out}}{d\tau}=a_1-a_2i_{out}+x_1\cdot\gamma_i\cdot\left(i_{1,in}-i_{out}\right)-x_2\cdot\gamma_i\cdot\left(i_{out}-i_{2,in}\right) \quad [S2.40]$$

$$\frac{dp_{out}}{d\tau}=a_2i_{out}+x_2\cdot\gamma_p\cdot\left(p_{2,in}-p_{out}\right)-x_1\cdot\gamma_p\cdot\left(p_{out}-p_{1,in}\right) \quad [S2.41]$$

The ODEs regarding the growth of the two populations remain unchanged as defined by Eqn. [14]-[17]. Again, we cannot solve sole solutions from the steady-state equations, but obtained two equations regarding the steady-state biomass of the two populations, which were similar as the ones derived from above scenarios

$$\frac{a_1c_1Ig_1y_1\gamma_p(Ig_2+\gamma_p)x_1\left(1-\frac{x_1+x_2}{\rho}\right)}{\gamma_p\left(Ig_1x_1+Ig_2x_2\right)+Ig_1Ig_2(x_1+x_2)}=d_1 \quad [S2.42]$$

$$\frac{a_1c_2Ig_2y_2\gamma_p(Ig_1+\gamma_p)x_1\left(1-\frac{x_1+x_2}{\rho}\right)}{\gamma_p\left(Ig_1x_1+Ig_2x_2\right)+Ig_1Ig_2(x_1+x_2)}=d_2 \quad [S2.43]$$

Eqn. [S2.35] can be also derived from these two equations. We also tested the validity of Eqn. [S2.35] in this scenario using numeric simulations following same protocol as used in S2.2.3, from which we obtained almost identical results as in S2.2.3 (Figure S3C). Taken together, we concluded that once the second metabolic reaction is performed extracellularly, the co-existence of two populations requires strain [1, 0] and [0, 1] possess same inherent fitness, and the steady-state structure of a MDOL community in this scenario relies on initial conditions.

Actually, when the second metabolic reaction is performed extracellularly, P is generated in the extracellular space, and becomes equivalently available for both populations. As a result, [0, 1] population no longer obtains 'private benefit' as in our basic model. The two populations simply compete for the limited resource (P), thus the assembly rule is similar as the one defined by the classical 'Competitive exclusive principle'[4], in which the co-existence of the populations requires very harsh condition: the competitive populations must possess totally similar ability for resource competition (i.e., same inherent fitness in the given competition). From this perspective, we concluded that once the second metabolic step is performed extracellularly, the criterion defining the stability of a MDOL community becomes tightening (Figure 2B).

**S2.3 Abiotic degradation of intermediate and product**

To consider the case where the intermediate and product can also be degraded without biological enzymatic catalysis (Figure S4A), we modified the basic form of our ODE system to include processes of abiotic degradation, which follows one-order reaction kinetics. The system of dimensionless ODEs corresponding to kinetics regarding Intermediate and product dynamics is thus given by:

$$\frac{di_{1,in}}{d\tau}=a_1-\gamma_i\cdot\left(i_{1,in}-i_{out}\right)-l_i\cdot i_{1,in} \quad [S2.44]$$

$$\frac{di_{2,in}}{d\tau}=-a_2 i_{2,in}+\gamma_i\cdot\left(i_{out}-i_{2,in}\right)-l_i\cdot i_{2,in} \quad [S2.45]$$

$$\frac{dp_{1,in}}{d\tau}=-Ig_1 p_{1,in}+\gamma_p\cdot\left(p_{out}-p_{1,in}\right)-l_p\cdot p_{1,in} \quad [S2.46]$$

$$\frac{dp_{2,in}}{d\tau}=a_2 i_{2,in}-Ig_2 p_{2,in}-\gamma_p\cdot\left(p_{2,in}-p_{out}\right)-l_p\cdot p_{2,in} \quad [S2.47]$$

$$\frac{di_{out}}{d\tau}=x_1\cdot\gamma_i\cdot\left(i_{1,in}-i_{out}\right)-x_2\cdot\gamma_i\cdot\left(i_{out}-i_{2,in}\right)-l_i\cdot i_{out} \quad [S2.48]$$

$$\frac{dp_{out}}{d\tau} = x_2 \cdot \gamma_p \cdot \left(p_{2,in} - p_{out}\right) - x_1 \cdot \gamma_p \cdot \left(p_{out} - p_{1,in}\right) - l_p \cdot p_{out} \quad \text{[S2.49]}$$

Here, $l_i$ is the abiotic degradation rate of I, while $l_p$ is the abiotic degradation rate of P. We simplified the steady-state expressions of the equations, and obtained two equations that determine the assembly of a MDOL community at steady-state:

$$\frac{\rho(l_p m+(-1+m-n+mn_2)\gamma_p)(l_i R_{[0,1]}(l_i+\gamma_i)(a_2+l_i+\gamma_i)+l_i(l_i+\gamma_i)x_2+a_2(l_i+R_{[0,1]}\gamma_i)x_2)}{krp} + \frac{a_1 a_2 c_1 n\gamma_i(\rho \cdot R_{[0,1]}-x_2)x_2 y_1 y_2}{d_1(-1+R_{[0,1]})} = 0$$

[S2.50]

$$l_p^{\,2} + l_p(\gamma_p + n\gamma_p + \frac{x_2}{R_{[0,1]}}) + \gamma_p(x_2 - mx_2 + \frac{nx_2}{R_{[0,1]}}) = 0$$

[S2.51]

Here, $n_2 = Ig_2/\gamma_p$. We further assumed that at steady-state, the summarized biomass of the two populations are very close to maximum capacity, that is,

$$x_1 + x_2 \approx \rho \;\text{ or }\; \rho \cdot R_{[0,1]} - x_2 \approx 0 \quad \text{[S2.52]}$$

Accordingly, we solved the equations and obtained

$$R_{[0,1]} \approx \frac{j_1(-j_2+(n+j_2)(j_1-j_2+j_3))(j_1+j_3+j_4)}{j_2(1+n+j_2)j_3 j_4 + m \cdot j_1(j_1+j_3)(j_1+j_3+j_4)} \quad \text{[S2.53]}$$

Here,

(1) $j_1 = l_i/\gamma_{p'}$ and $j_2 = l_p/\gamma_{p'}$, reflecting the effects of abiotic degradation of intermediate and product, respectively;

(2) $j_3 = \gamma_i/\gamma_p$, reflecting that the transport rate of the intermediate affects assembly of a MDOL community in this scenario.

(3) $j_4 = a_2/\gamma_p$, reflecting that the rate of second reaction affects assembly of a MDOL community in this scenario.

(4) Note that $n$ and $m$ still play important roles in shaping community structure at the steady-state.

We further applied numerical simulations to generate the parameter spaces that result in co-existence of the two MDOL populations in the community, as well as analyzed the assembly of those stable communities. We further compared the size of the parameter space at different sets of $l_i$ and $l_p$ to address how abiotic degradation of metabolites affect the assembly rule.

As shown in Figure S4B, compared with the basic rule, when intermediate and product can be degraded abiotically, the size of the parameter space reduces, indicating that a MDOL community becomes more difficult to maintain stability in this case. This is because the scenario of abiotic degradation decreases the production of the limiting resource, P. Therefore, P becomes more unavailable in the system, the asymmetric private benefit of [0, 1] is strengthened, so that [1, 0] need to hold larger growth edge to neutralize this asymmetry, resulted in more stringent conditions for the maintenance of the co-existence.

**S2.4 Substrate concentration is comparable to the Michaelis-Menten constant of the first reaction**

We then relaxed our assumption (5) to consider the scenario that the substrate concentration is comparable to the Michaelis-Menten constant of the first reaction. Therefore, the system of dimensionless ODEs corresponding to pathway kinetics becomes more complicated, given by:

$$\frac{ds_{1,in}}{d\tau}=-\frac{a_1 \cdot s_{1,in}}{(1+s_{1,in})}+\gamma_s \cdot (s_{out}-s_{1,in}) \qquad [S2.54]$$

$$\frac{ds_{2,in}}{d\tau}=\gamma_s \cdot (s_{out}-s_{2,in}) \qquad [S2.55]$$

$$\frac{ds_{out}}{d\tau}=-x_1 \cdot \gamma_s \cdot \left(s_{out}-s_{1,in}\right)-x_2 \cdot \gamma_s \cdot \left(s_{out}-s_{2,in}\right) \qquad [S2.56]$$

$$\frac{di_{1,in}}{d\tau}=\frac{a_1 \cdot s_{1,in}}{\left(1+s_{1,in}\right)}-\gamma_i \cdot \left(i_{1,in}-i_{out}\right) \qquad [S2.57]$$

$$\frac{di_{2,in}}{d\tau}=-a_2 i_{2,in}+\gamma_i \cdot \left(i_{out}-i_{2,in}\right) \qquad [S2.58]$$

$$\frac{dp_{1,in}}{d\tau}=-Ig_1 p_{1,in}+\gamma_p \cdot \left(p_{out}-p_{1,in}\right) \qquad [S2.59]$$

$$\frac{dp_{2,in}}{d\tau}=a_2 i_{2,in}-Ig_2 p_{2,in}-\gamma_p \cdot \left(p_{2,in}-p_{out}\right) \qquad [S2.60]$$

$$\frac{di_{out}}{d\tau}=x_1 \cdot \gamma_i \cdot \left(i_{1,in}-i_{out}\right) - x_2 \cdot \gamma_i \cdot \left(i_{out}-i_{2,in}\right) \qquad [S2.61]$$

$$\frac{dp_{out}}{d\tau}=x_2 \cdot \gamma_p \cdot \left(p_{2,in}-p_{out}\right)-x_1 \cdot \gamma_p \cdot \left(p_{out}-p_{1,in}\right) \qquad [S2.62]$$

It is difficult to derive a simple formula using the same derivation methods as the basic model, because steady-state solutions here rely on the initial conditions (in particular, the initial concentration of S). In addition, this system simulates batch culture of the microbial community, in which the community will finally collapses once the substrate was used up. However, our numeric simulations showed that when the substrate concentration was sufficient over a threshold, [1, 0] and [0, 1] can still maintain stability before the community collapse with some given condition (Figure S2.1A). Therefore, adequate supply of the substrate is a prerequisite for the stability of a MDOL community. In addition, our simulation also revealed that once the steady-state is present, the assembly of the final stable communities are independent on the initial substrate concentration (Figure S2.1). Therefore, in these well-mixed systems, initial substrate concentration may only affect whether the steady-state is present but not change the assembly of those stable MDOL communities. The analytic solutions for the steady-state can be further obtained by assuming that the supply of substrate in the system is constant, that is

$$\frac{ds_{out}}{d\tau} \approx 0 \qquad [S2.63]$$

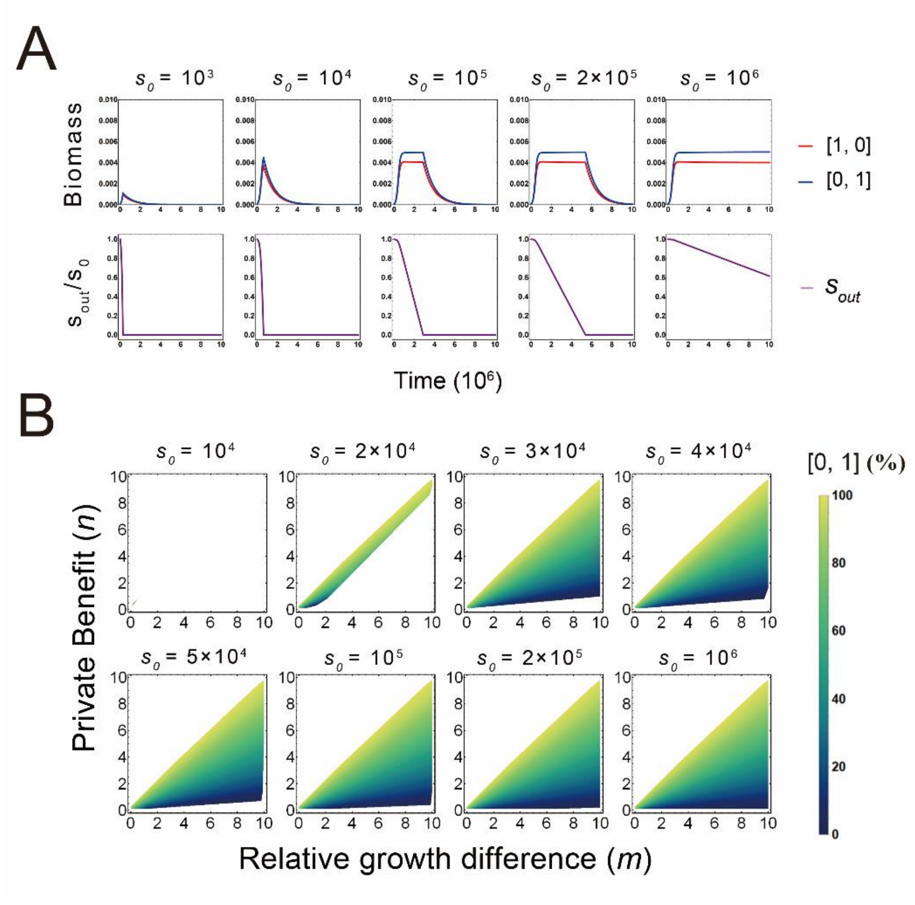

**Figure S2.1** adequate supply of the substrate is a prerequisite for the stability of a MDOL community. (A) Representative simuation dynamics of the MDOL communities. Here, we set $c_2$=1/7 and $\gamma_p$=3.0, while the default values given in Table S2 are used for other parameters.The first row shows the relative abundance dynamics of the two populations in the community, and the second row shows the depletion dynamics of substrate. (B) The density maps show the parameter spaces characterized by private benefit $n$ and relative growth difference $m$, as well as relative abundance of [0, 1] in the stable communities, under different initial substrate ($s_0$). These plots are generated by mathematical simulations. Values of other parameters are given in Table S2.

Given this constraint, we obtained the totally same assembly rule as defined by Eqn. [1] and Eqn. [2] from analytic analysis. Consistent with this analysis, when the supply of S is over a threshold, the density map defining the assembly rule becomes very similar as the one derived from the basic model (Figure S2.1B). Thus, if the supply of S in the system is sufficient, relaxing our assumption (4) will not change the basic assembly rule of a two-step MDOL community.

Notably, we applied this principle to the culturing of our synthetic microbial community. Since the solubility of naphthalene in water is lower, if we supply sufficient naphthalene, its extracellular concentration should constantly equal to its saturated concentration (meaning $\frac{ds_{out}}{d\tau} \approx 0$). Therefore, we can test the predictions from model by batch culturing of our synthetic microbial community supplying sufficient amount of naphthalene.

#### S2.5 Intermediate concentration is comparable to the Michaelis-Menten constant of the second reaction

We next considered the scenario that the intermediate concentration is comparable to the Michaelis-Menten constant of the second reaction (relaxing the first part of our assumption (5)). Therefore, the system of dimensionless ODEs corresponding to pathway kinetics is given by:

$$\frac{di_{1,in}}{d\tau} = a_1 - \gamma_i \cdot \left(i_{1,in} - i_{out}\right) \quad [S2.64]$$

$$\frac{di_{2,in}}{d\tau} = -\frac{a_2' \cdot i_{2,in}}{\left(\beta_2 + i_{2,in}\right)} + \gamma_i \cdot \left(i_{out} - i_{2,in}\right) \quad [S2.65]$$

$$\frac{dp_{1,in}}{d\tau}=-Ig_1 p_{1,in}+\gamma_p\cdot\left(p_{out}-p_{1,in}\right) \quad [S2.66]$$

$$\frac{dp_{2,in}}{d\tau}=\frac{a_2^{'}\cdot i_{2,in}}{\left(\beta_2+i_{2,in}\right)}-Ig_2 p_{2,in}-\gamma_p\cdot\left(p_{2,in}-p_{out}\right) \quad [S2.67]$$

$$\frac{di_{out}}{d\tau}=x_1\cdot\gamma_i\cdot\left(i_{1,in}-i_{out}\right)-x_2\cdot\gamma_i\cdot\left(i_{out}-i_{2,in}\right) \quad [S2.68]$$

$$\frac{dp_{out}}{d\tau}=x_2\cdot\gamma_p\cdot\left(p_{2,in}-p_{out}\right)-x_1\cdot\gamma_p\cdot\left(p_{out}-p_{1,in}\right) \quad [S2.69]$$

Conducting with the same analytic method, we obtained the totally same assembly rule as defined by Eqn. [1] and Eqn. [2]. Therefore, even the accumulation of I makes its concentration comparable to the Michaelis-Menten constant of the second reaction, the assembly of a MDOL community still follows our basic rule.

**S2.6 Product concentration is comparable to the Half-saturation constant of the Monod growth kinetics**

To further test whether relaxing our assumption (5) will change our assembly rule of a MDOL community, we next considered the scenario that product concentration is comparable to the Half-saturation constant of the Monod growth kinetics. Here, the system of dimensionless ODEs becomes more complicated, given by:

$$\frac{di_{1,in}}{d\tau}=a_1-\gamma_i\cdot\left(i_{1,in}-i_{out}\right) \quad [S2.70]$$

$$\frac{di_{2,in}}{d\tau}=-a_2\cdot i_{2,in}+\gamma_i\cdot\left(i_{out}-i_{2,in}\right) \quad [S2.71]$$

$$\frac{dp_{1,in}}{d\tau}=-\frac{Ig_1^{'}p_{1,in}}{\beta_{g1}+p_{1,in}}+\gamma_p\cdot\left(p_{out}-p_{1,in}\right) \quad [S2.72]$$

$$\frac{dp_{2,in}}{d\tau}=a_2\cdot i_{2,in}-\frac{Ig_2^{'}p_{2,in}}{\beta_{g2}+p_{2,in}}-\gamma_p\cdot\left(p_{2,in}-p_{out}\right) \quad [S2.73]$$

$$\frac{di_{out}}{d\tau}=x_1\cdot\gamma_i\cdot\left(i_{1,in}-i_{out}\right)-x_2\cdot\gamma_i\cdot\left(i_{out}-i_{2,in}\right) \quad [S2.74]$$

$$\frac{dp_{out}}{d\tau}=x_2\cdot\gamma_p\cdot\left(p_{2,in}-p_{out}\right)-x_1\cdot\gamma_p\cdot\left(p_{out}-p_{1,in}\right) \quad [S2.75]$$

$$\frac{dx_1}{d\tau}=\mu_1 x_1\left(1-\frac{x_1+x_2}{\rho}\right)-d_1 x_1 \quad [S2.76]$$

$$\frac{dx_2}{d\tau}=\mu_2 x_2\left(1-\frac{x_1+x_2}{\rho}\right)-d_2 x_2 \quad [S2.77]$$

$$\mu_1=\frac{Ig_1' p_{1,in}}{\beta_{31}+p_{1,in}} y_1 c_1 \quad [S2.78]$$

$$\mu_2=\frac{Ig_2' p_{2,in}}{\beta_{32}+p_{2,in}} y_2 c_2 \quad [S2.79]$$

Via mathematical derivation, we obtained a nonlinear equation that determines the assembly of the steady-state community

$$\left(\frac{1-R_{[0,1]}}{R_{[0,1]}(1-R_{[0,1]}+u\cdot R_{[0,1]})}-\left(\frac{v_1}{R_{[0,1]}}+b_1\right)\right)\left(\frac{u\cdot R_{[0,1]}}{(1-R_{[0,1]}+u\cdot R_{[0,1]})}-v_2\right)+b_2 u\left(\frac{1-R_{[0,1]}}{1-R_{[0,1]}+u\cdot R_{[0,1]}}-v_1\right)=0$$

[S2.80]

Here,

(1) $u=\frac{c_1 y_1}{d_1}/\frac{c_2 y_2}{d_2}$, reflecting the relative fitness cost between the two populations.

(2) $v_1=\frac{Ig_1'}{a_1}$, $v_2=\frac{Ig_2'}{a_1}$, $b_1=\frac{\beta_{g1}}{a_1\gamma_p}$, $b_2=\frac{\beta_{g2}}{a_1\gamma_p}$.

Therefore, in this scenario, the assembly of a two-step MDOL community is determined by these five parameters. We next performed numeric simulations to investigate how these parameters affect our basic assembly rule of a MDOL community. As shown in Figure S2.2, the size of parameter space for the co-existence of the two populations largely decreases compared with the rule defined by Eqn. [2], suggesting that a MDOL community becomes more difficult to reach stable when product concentration is comparable to the Half-saturation constant of the Monod growth kinetics. Moreover, with the increase of $b_1$and $b_2$, the size parameter space gradually increase to apporach the basic condition. In summary, relaxing assumption (5) may results in more strict condition for the co-existence of the members in a two-step MDOL community.

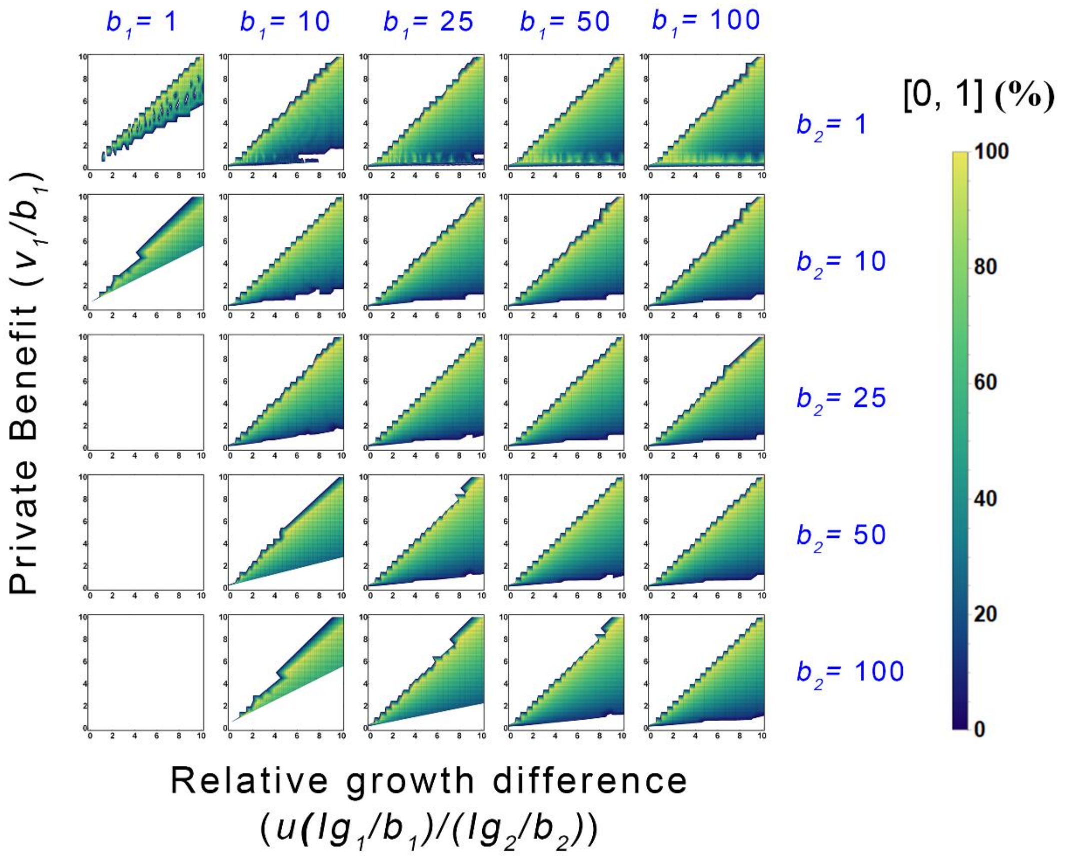

**Figure S2.2** when product concentration is comparable to the Half-saturation constant of the Monod growth kinetics, co-existence of the two populations becomes more difficult than the scenario in our basic model. The density map array shows the range of stable parameter set characterized by privatization benefit $n$ and relative growth advantage $m$, as well as relative abundance of embezzler population under different values of $u$, $v_1$, $v_2$, $b_1$, and $b_2$. In these simulations, the definition of $n$ and $m$ is slightly different, as shown in the axis titles. The default values given in Table S2 are used for all other parameters.

### S2.7 Presence of byproduct generated from the first reaction

We next considered the case in which metabolic by-products can be generated from the first-step reaction. We hypothesized that a by-product, P', is generated during the conversion of S to I by [1, 0], which can also serve as the limited carbon source to support the growth of both populations (Figure S5A). The consuming rates of P' of the two populations were assumed to be $bg_1$ and $bg_2$, respectively. In addition, the relative contribution of P' to P to the biomass accumulation was assumed as $\delta$. Following these assumptions, the system of dimensionless ODEs turned into:

$$\frac{di_{1,in}}{d\tau}=a_1-\gamma_i\cdot\left(i_{1,in}-i_{out}\right) \quad [S2.81]$$

$$\frac{di_{2,in}}{d\tau}=a_2i_{2,in}+\gamma_i\cdot\left(i_{out}-i_{2,in}\right) \quad [S2.82]$$

$$\frac{dp'_{1,in}}{d\tau}=a_1-bg_1p'_{1,in}-\gamma_{p'}\cdot\left(p'_{1,in}-p'_{out}\right) \quad [S2.83]$$

$$\frac{dp'_{2,in}}{d\tau}=\gamma_{p'}\cdot\left(p'_{out}-p'_{2,in}\right)-bg_2p'_{2,in} \quad [S2.84]$$

$$\frac{dp_{1,in}}{d\tau}=-Ig_1p_{1,in}+\gamma_p\cdot\left(p_{out}-p_{1,in}\right) \quad [S2.85]$$

$$\frac{dp_{2,in}}{d\tau}=a_2i_{2,in}-Ig_2p_{2,in}-\gamma_p\cdot\left(p_{2,in}-p_{out}\right) \quad [S2.86]$$

$$\frac{di_{out}}{d\tau}=x_1\cdot\gamma_i\cdot\left(i_{1,in}-i_{out}\right)-x_2\cdot\gamma_i\cdot\left(i_{out}-i_{2,in}\right) \quad [S2.87]$$

$$\frac{dp'_{out}}{d\tau}=x_1\cdot\gamma_{p'}\cdot\left(p'_{1,in}-p'_{out}\right)-x_2\cdot\gamma_{p'}\cdot\left(p'_{out}-p'_{2,in}\right) \quad [S2.88]$$

$$\frac{dp_{out}}{d\tau}=x_2\cdot\gamma_p\cdot\left(p_{2,in}-p_{out}\right)-x_1\cdot\gamma_p\cdot\left(p_{out}-p_{1,in}\right) \quad [S2.89]$$

$$\frac{dx_1}{d\tau}=\mu_1x_1\left(1-\frac{x_1+x_2}{\rho}\right)-d_1x_1 \quad [S2.90]$$

$$\frac{dx_2}{d\tau}=\mu_2x_2\left(1-\frac{x_1+x_2}{\rho}\right)-d_2x_2 \quad [S2.91]$$

$$\mu_1=(Ig_1 p_{1,in}+\delta\cdot bg_1 p'_{1,in})y_1 c_1 \qquad [S2.92]$$

$$\mu_2=(Ig_2 p_{2,in}+\delta\cdot bg_2 p'_{2,in})y_2 c_2 \qquad [S2.93]$$

Via mathematical derivation, we obtained a nonlinear equation that determines the assembly of the steady-state community

$$\frac{R_{[0,1]}p_1(1-R_{[0,1]}+p_2)}{p_1}=\frac{a_1m_2(R_{[0,1]}+n_1)n_2+R_{[0,1]}\rho(n_2R_{[0,1]}+n_1(1-R_{[0,1]}+n_2)+a_1m_1((-1+R_{[0,1]})n_1-(n_2R_{[0,1]}+n_1(1-R_{[0,1]}+n_2))\delta))}{R_{[0,1]}(a_1(-1+R_{[0,1]})m_1n_1+n_2R_{[0,1]}+n_1(1-R_{[0,1]}+n_2))\rho+a_1m_2(-R_{[0,1]}n_2(-1+\delta)+n_1(n_2+(-1+R_{[0,1]})\delta-n_2\delta))}$$

[S2.94]

Here,

(1) $p_1 = bg_1/\gamma_{p'}$ and $p_2 = bg_2/\gamma_{p''}$ which quantify the effects of metabolic byproduct to the two populations, respectively;

(2) $n_1 = Ig_1/\gamma_p$ and $n_2 = Ig_2/\gamma_p$, reflecting the 'Product demand gap' of the two populations, respectively.

(3) $m_1 = y_1c_1/d_1$ and $m_2 = y_2c_2/d_2$, reflecting the inherent fitness of the two populations, respectively.

Note that $m=(\frac{c_1Ig_1y_1}{d_1}-\frac{c_2Ig_2y_2}{d_2})/\frac{c_2Ig_2y_2}{d_2} = (m_1n_1-m_2n_2)/m_2n_2$, and $n = Ig_1/\gamma_p = n_1$. Thus, Eqn. [S2.94] reflects that the main factors in presence of metabolic byproduct including $m$ and $n$ (also present in the basic equation Eqn. [1]), $p_1$ and $p_2$ (the effects of metabolic byproduct), $a_1$ (rate of the first reaction), as well as $\delta$ (relative contribution of byproduct to population growth). We next performed numeric simulation to address how the presence of byproduct affect the assembly rule of a two-step MDOL community. As shown in Figure S5B, our simulations indicated that the presence of metabolic by-products would increase the size of the parameter space for the co-existence of two populations, suggesting that by-product generation facilitates the stability of a MDOL community. Notably, with the increase of the $\delta$ (that is, if the by-product contributes more to the biomass accumulation), or with the increase the rate of the first reaction ($a_1$), the size of the parameter space significantly increases. This result indicated that when the rate of first reaction and availability of by-products reaches a high level, co-existence of the two populations may be maintained even if the private benefit of [0, 1] (*n*) are greater than relative growth advantages of [1, 0] (*m*). This shift is because the by-product is generated from the first reaction that occurs inside the cells of [1, 0], so that [1, 0] cells possess the preferential access to the by-product, resulted in a 'private benefit' from the acquirement of this limiting resource. This the byproduct 'private benefit' of [1, 0] may neutralize the 'private benefit' from the end product of [0, 1], thus maintain the balance of the resource allocation and promote the co-existence of the two populations in a MDOL community.

**S2.8 Biotoxicity of substrate, intermediate and product**

We next adjusted our basic model to account for the presence biotoxicity of substrate, intermediate and product. If the two populations possess different tolerance to the biotoxicity of the involved substance, the relative fitness of the two populations should be affected by their relative ability of toxic tolerance. However, it is difficult to quantify this relative ability, because it is highly dependent on physiological properties of the strains involved in specific microbial systems. In addition, the effects of this difference can be also treated as a part of 'inherent fitness' of each population, thus can be partially quantified by the existed parameters $c_1$ and $c_2$. However, the biotoxicity of a substrate or metabolite is inseparable from its local concentration. The goal of this part is to investigate whether and how the differences in heterogeneity of local substance concentration influence the relative toxic effects of the substance to the two populations, and then govern the assembly of a MDOL community.

Therefore, here we add a model assumption that both populations possessed same ability of toxic tolerance to S, I, or P. In addition, our study assumes that S, I, and P are rapidly well-mixed in the extracellular space (assumption (1)). Thus, if bio-toxic effects occur extracellularly (for example, directly harms the cell wall or membrane), it will have same impact on the both populations since the extracellular concentration of S, I, and P for the two populations is totally same. Therefore, in this section, we only discuss the scenario in which the bio-toxic effects occur intracellularly. These effects were represented by parameters $T_{S,i}$, $T_{I,i}$ and $T_{P,i}$ in the models, which represents the strength of the intracellular bio-toxicity of S, I or P to the *i*th population, respectively. We considered three different expression forms to characterize each toxic coefficient[5], that is, reciprocal form, linear form and exponential form. For example, the three expressions of $T_{S,i}$ are given by:

$$T_{S,i,R}=\frac{1}{1+\frac{S_{i,in}}{K_{s,tox}}} \quad [S2.95]$$

$$T_{S,i,L}=1-\frac{S_{i,in}}{K_{s,tox}} \quad [S2.96]$$

$$T_{S,i,E}=e^{-\frac{S_{i,in}}{K_{s,tox}}} \quad [S2.97]$$

To fit into our dimensionless model, we non-dimensionalized Eqns. [S2.95]-[S2.97], given by

$$t_{s,i,r}=\frac{1}{1+\theta_s s_{i,in}} \quad [S2.98]$$

$$t_{s,i,l}=1-\theta_s s_{i,in} \quad [S2.99]$$

$$t_{s,i,e}=e^{-\theta_s s_{i,in}} \quad [S2.100]$$

In which a novel non-dimensionalized variable $\theta_s$ is defined as

$$\theta_s=\frac{K_1}{K_{s,tox}} \quad [S2.101]$$

Therefore, the dimensionless ODEs regarding the growth of the two populations are given by

$$\frac{dx_1}{d\tau}=\mu_1 t_{s,1} x_1\left(1-\frac{x_1+x_2}{\rho}\right)-d_1 x_1 \quad [S2.102]$$

$$\frac{dx_2}{d\tau}=\mu_2 t_{s,2} x_2\left(1-\frac{x_1+x_2}{\rho}\right)-d_2 x_2 \quad [S2.103]$$

Similar as described above, the dimensionless parameters of $T_{S,i}$, $T_{I,i}$ and $T_{P,i}$ are summarized in detail in Table S3.

*S2.7.1 Biotoxicity of substrate*

We modified our basic ODE system to investigate how intracellular biotoxicity of S affects the assembly of a two-step MDOL community (Figure S6A). Based on the modified ODEs, we derived a novel formula, as well as the associated condition, as follow

$$R_{[0,1]}=\frac{n}{(m+1)to_s-1} \quad [S2.104]$$

$$0< n <(m+1)to_s-1 \quad [S2.105]$$

Here, $to_s$ was a derived parameter that defines the effect of intracellular biotoxicity of S to microbial community assembly. When the toxic coefficient ($t_{s,i,r}$) follows the reciprocal form (Eqn. [S2.93]), the linear form (Eqn. [S2.94]) and the exponential form

(Eqn. [S2.95]), the expressions of $to_s$ are respectively given by

$$to_{s,r}=\frac{1+\theta_s s_0}{(1+\theta_s s_0)-\frac{a_1\theta_s}{\gamma_s}} \quad [S2.106]$$

$$to_{s,l}=\frac{a_1\theta_s}{\gamma_s}\cdot\frac{1}{(1-\theta_s s_0)}+1 \quad [S2.107]$$

$$to_{s,e}=e^{\frac{a_1\theta_s}{\gamma_s}} \quad [S2.108]$$

Eqn. [S2.101]- [S2.103] indicates that, no matter what form $to_s$ is, its value is greater than 1, which further suggests that in Eqn. [S2.100], the formula $(m+1)to_s\text{-}1 > m$ is always true. Therefore, compared with the basic condition shown in Eqn. [2], Eqn. [S2.100] suggests a relaxed condition for the stability of a MDOL community. Similarly, Eqn. [S2.99] suggests that the frequency of [0, 1] in the stable community increases in this case compared with the scenario in the basic model. These shifts are visualized in Figure S6B, which further indicates that size of parameter space increases with $to_s$, and the final relative abundance of [0, 1] significantly decreases with the increase of $to_s$.

The effect of toxic substrate is due to the different intracellularly accumulation level of S of the two populations. Since [1, 0] can intracellularly transfer S to I while [0, 1] cannot, S generally accumulates more inside the cells of [0, 1], resulted in greater toxic effect on [0, 1] and neutralizing its private benefit from P privatization. Our mathematical analysis indicates that this effect can be quantified by a simple parameter $to_s$, which is determined by four specific parameters, that is, the toxic strength ($\theta_s$), initial substrate concentration ($s_0$), rate of the first reaction ($a_1$), and the diffusivity coefficient of S ($\gamma_s$). In summary, intracellular biotoxicity of substrate facilitate the co-existence of the two MDOL members, and favors the population executing the first metabolic step.

*S2.7.2 Biotoxicity of intermediate*

Next, we turned to investigate how intracellular biotoxicity of intermediate affects the assembly of a two-step MDOL community (Figure S7A). However, the modified ODE system cannot be directly solved by the same derivation methods as the basic model. Alternatively, we derived nonlinear equations that determines the assembly of the steady-state community.

(1) when the toxic index ($t_i$) follows the reciprocal form:

$$\frac{(q_1 - q_2)R_{[0,1]} - q_1}{(q_1 - q_2)R_{[0,1]} - q_1(q_2+1)} = \frac{n+R_{[0,1]}}{(m+1)\cdot R_{[0,1]}} \qquad [S2.109]$$

(2) when $t_i$ follows the linear form

$$\frac{q_2R_{[0,1]} + q_1(R_{[0,1]} - q_2 - 1)}{q_2R_{[0,1]} + q_1(R_{[0,1]} - 1)} = \frac{n+R_{[0,1]}}{(m+1)\cdot R_{[0,1]}} \qquad [S2.110]$$

(3) when $t_i$ follows the exponential form

$$e^{\frac{q_1}{R_{[0,1]}}} = \frac{n+R_{[0,1]}}{(m+1)\cdot R_{[0,1]}} \qquad [S2.111]$$

Here, $q_1$ reflects the accumulation speed of the toxic effect of I, defined as

$$q_1 = \frac{a_1\theta_i}{\gamma_i} \qquad [S2.112]$$

$q_2$ reflects the consuming speed of I, defined as

$$q_2 = \frac{a_2}{\gamma_i} \qquad [S2.113]$$

Eqn. [S2.109]- [S2.111] indicates that $q_1$ and $q_2$ largely affect the assembly of a two-step MDOL community in presence of intermediate toxicity. Thus, we performed numeric simulations to test the effects of $q_1$, $q_2$. As shown in Figure S7B-D, we found that the presence of intermediate toxicity generally decreases the size of the parameter space for the stability of a MDOL community. The effect of $q_2$ is more sensitive than $q_1$. With the increase of $q_2$, the size of the parameter space significant decreases, suggesting that a MDOL community is more difficult to maintain stability.

In two-step MDOL system, I is transferred from S and accumulated intracellularly in [1, 0]. In contrast, it can be converted into P in [0, 1]. Therefore, the intracellular I concentration of [1, 0] is higher than that of [0, 1], so that toxic effects of I on [1, 0] is greater than that of [0, 1]. [0, 1] thus obtains more growth advantage which harms the co-existence of the two population. Consistent with this prediction, our model offers a quantitative profile that this effect can be characterized by parameter $q_1$ and $q_2$, as described above. In summary, intracellular biotoxicity of intermediate harms the co-existence of the two MDOL members, and favors the population executing the second metabolic step.

*S2.7.3 Biotoxicity of product*

To investigate how intracellular biotoxicity of end product affects the assembly of a two-step MDOL community (Figure S8A), we applied same analyzing method as used in the above section. The nonlinear equations that determines the assembly of the steady-state community are given by

(1) when the toxic index ($t_p$) follows the reciprocal form:

$$\frac{w_1\left(R_{[0,1]} - 1\right)\left(n+R_{[0,1]}\right) + n\left(R_{[0,1]} - 1 - w_1\right)R_{[0,1]} - w_2 R_{[0,1]}^2}{\left(w_1\left(R_{[0,1]} - 1\right) - w_2 R_{[0,1]} + n\left(R_{[0,1]} - 1 - w_2\right)\right)R_{[0,1]}} = \frac{n+R_{[0,1]}}{(m+1)R_{[0,1]}} \quad \text{[S2.114]}$$

(2) when $t_p$ follows the linear form

$$\frac{w_1(R_{[0,1]}-1)(n+R_{[0,1]})+n(R_{[0,1]}-1-w_1)R_{[0,1]}-w_2R_{[0,1]}^2}{(w_1(R_{[0,1]}-1)-w_2R_{[0,1]}+n(R_{[0,1]}-1-w_2))R_{[0,1]}}=\frac{n+R_{[0,1]}}{(m+1)R_{[0,1]}} \quad [S2.115]$$

(3) when $t_p$ follows the exponential form

$$e^{-\frac{R_{[0,1]}(n(R_{[0,1]}-1))}{R_{[0,1]}(n(R_{[0,1]}-1-w_1)R_{[0,1]}-w_2R_{[0,1]})}}=\frac{n+R_{[0,1]}}{(m+1)R_{[0,1]}}$$

[S2.116]

Here, $w_1$ reflects the accumulation speed of the toxic effect of P, defined as

$$w_1=\frac{a_1\theta_p}{\gamma_p} \quad [S2.117]$$

$w_2$ reflects the consuming speed of P, defined as

$$w_2=\frac{Ig_2}{\gamma_p} \quad [S2.118]$$

Inspired by Eqn. [S2.114]- [S2.116], we performed numeric simulations to test how $w_1$ and $w_2$ affect the assembly of a two-step MDOL community. As shown in Figure S8B-D, we found that the presence of product toxicity strongly inhibits the growth of [0, 1], and harms the stability of a MDOL community. When $t_p$ follows the reciprocal form, with the increase of $w_1$, or decrease of $w_2$, although the size of the parameter space rarely changed, the relative frequency of [0, 1] in those stable MDOL communities significantly decreases (Figure S8B), in which [1, 0] largely dominates the community (Figure S8E). When the relative abundance of [0, 1] is lower like that, occasionally random loss may result in the its extinction[6,7] when facing environmental fluctuations. Therefore, this seemingly stable community may be also evolutionary unstable. When $t_p$ follows the linear form or exponential form, the presence of intermediate toxicity decreases the size of the parameter space for the stability of a MDOL community (Figure S8C-D). In the cases even when [1, 0] possess a growth advantage (*m*) over the private benefit (*n*), the community exhibited collapse dynamics due to the rapid growth of [1, 0] (Figure S8F).

Theoretically, P is produced and accumulated in the cells of [0, 1], thus the intracellular toxicity of P influences the growth of [0, 1] more than [1, 0]. Consistently, our model predicts that the presence of P biotoxicity strongly inhibits [0, 1]. In addition, our model also indicates this inhibitory effect does not play a role to neutralize the private benefit of [0, 1] but directly destabilizes the community. As described above, this effect can be quantified by parameter $w_1$ and $w_2$. In summary, intracellular biotoxicity of product inhibits the population executing the second metabolic step, and also harms the co-existence of the two MDOL members.

### S3 Derivation of the model regarding the assembly of a multi-step MDOL community

#### S3.1 Derivation of the model regarding the assembly of a three-step MDOL community

In this section, we expanded our basic model to characterize the dynamics of a three-step MDOL community. In this community, a population [1, 0, 0] degrades a substrate (S) into the first intermediate (I1), which is then converted to the second intermediate (I2) by the second population [0, 1, 0]. I2 is finally transferred to the end product (P) by the last population named [0, 0, 1] (Figure S16A). Construction of this model still follows the seven biological assumptions listed in Section S1.

*S3.1.1 Equations regarding Intermediates and product dynamics*

We first built a system of ODEs describing dynamics of Intermediates and product concentrations for an intracellular pathway in a three-step MDOL community. The model is simplified by the same assumptions as described in Section S1, given by

$$\frac{dI1_{1,in}}{dt}\cdot V_c=k_1E_1-r_{I1}\cdot V_c\cdot\left(I1_{1,in}-I1_{out}\right) \quad [S3.1]$$

$$\frac{dI1_{2,in}}{dt}\cdot V_c=-\frac{k_2E_2}{K_2}I1_{2,in}+r_{I1}\cdot V_c\cdot\left(I1_{out}-I1_{2,in}\right) \quad [S3.2]$$

$$\frac{dI1_{3,in}}{dt}\cdot V_c=r_{I1}\cdot V_c\cdot\left(I1_{out}-I1_{3,in}\right) \quad [S3.3]$$

$$\frac{dI2_{1,in}}{dt}\cdot V_c=r_{I2}\cdot V_c\cdot\left(I2_{out}-I2_{1,in}\right) \quad [S3.4]$$

$$\frac{dI2_{2,in}}{dt}\cdot V_c=\frac{k_2E_2}{K_2}I2_{2,in}+r_{I2}\cdot V_c\cdot\left(I_{out}-I_{2,in}\right) \quad [S3.5]$$

$$\frac{dI2_{3,in}}{dt}\cdot V_c=-\frac{k_3E_3}{K_3}I2_{3,in}-r_{I2}\cdot V_c\cdot\left(I2_{3,in}-I3_{out}\right) \quad [S3.6]$$

$$\frac{dP_{1,in}}{dt}\cdot V_c=-\frac{kg_1}{Kg_1}\ P_{1,in}+r_P\cdot V_c\cdot\left(P_{out}-P_{1,in}\right) \quad [S3.7]$$

$$\frac{dP_{2,in}}{dt}\cdot V_c=-\frac{kg_2}{Kg_2}\ P_{2,in}+r_P\cdot V_c\cdot\left(P_{out}-P_{2,in}\right) \quad [S3.8]$$

$$\frac{dP_{3,in}}{dt}\cdot V_c=\frac{k_3E_3}{K_3}I2_{3,in}-\frac{kg_3}{Kg_3}\ P_{3,in}+r_P\cdot V_c\cdot\left(P_{out}-P_{3,in}\right) \quad [S3.9]$$

$$\frac{dI1_{out}}{dt}=\sum_{j=1}^{3}X_j\cdot r_{I1}\cdot(I1_{j,in}-I1_{out}) \quad [S3.10]$$

$$\frac{dI2_{out}}{dt}=\sum_{j=1}^{3}X_j\cdot r_{I2}\cdot(I2_{j,in}-I2_{out}) \quad [S3.11]$$

$$\frac{dP_{out}}{dt}=\sum_{j=1}^{3}X_j\cdot r_P\cdot(P_{j,in}-P_{out}) \quad [S3.12]$$

The equations were then non-dimensionalized following the method described in Table S4 and Table S5, which formalized a simple, non-dimensionalized model, given by

$$\frac{di1_{1,in}}{d\tau}=a_1-\gamma_{i1}\cdot\left(i1_{1,in}-i1_{out}\right) \quad [S3.13]$$

$$\frac{di1_{2,in}}{d\tau}=-a_2i1_{2,in}+\gamma_{i1}\cdot\left(i1_{out}-i1_{2,in}\right) \quad [S3.14]$$

$$\frac{di1_{3,in}}{d\tau}=\gamma_{i1}\cdot\left(i1_{out}-i1_{3,in}\right) \quad [S3.15]$$

$$\frac{di2_{1,in}}{d\tau}=\gamma_{i2}\cdot\left(i2_{out}-i2_{1,in}\right) \quad [S3.16]$$

$$\frac{di2_{2,in}}{d\tau}=a_2i2_{2,in}-\gamma_{i2}\cdot\left(i2_{2,in}-i2_{out}\right) \quad [S3.17]$$

$$\frac{di2_{3,in}}{d\tau}=-a_3i2_{3,in}+\gamma_{i2}\cdot\left(i2_{out}-i2_{3,in}\right) \quad [S3.18]$$

$$\frac{dp_{1,in}}{d\tau}=-Ig_1p_{1,in}+\gamma_P\cdot\left(p_{out}-p_{1,in}\right) \quad [S3.19]$$

$$\frac{dp_{2,in}}{d\tau}=-Ig_2p_{2,in}+\gamma_P\cdot\left(p_{out}-p_{2,in}\right) \quad [S3.20]$$

$$\frac{dp_{3,in}}{d\tau}=a_3i_{2,in}-Ig_3p_{3,in}-r_P\cdot\left(p_{3,in}-p_{out}\right) \quad [S3.21]$$

$$\frac{di1_{out}}{d\tau}=\sum_{j=1}^{3}x_j\cdot\gamma_{i1}\cdot(i1_{j,in}-i1_{out}) \quad [S3.22]$$

$$\frac{di2_{out}}{d\tau}=\sum_{j=1}^{3} x_j\cdot\gamma_{i2}\cdot(i2_{j,in}\text{-}i2_{out}) \qquad [S3.23]$$

$$\frac{dp_{out}}{d\tau}=\sum_{j=1}^{3} x_j\cdot\gamma_{i2}\cdot(p_{j,in}\text{-}p_{out}) \qquad [S3.24]$$

*S3.1.2 Equations regarding population growth dynamics*

Expanding the basic equations described in Section S1.1.2, our model then accounts for the growth dynamics of the two populations in a three-step MDOL community.

$$\frac{dX_1}{dt}=g_1X_1\left(1\text{-}\frac{X_1+X_2+X_3}{N_m}\right)\text{-}D_1X_1 \qquad [S3.25]$$

$$\frac{dX_2}{dt}=g_2X_2\left(1\text{-}\frac{X_1+X_2+X_3}{N_m}\right)\text{-}D_2X_2 \qquad [S3.26]$$

$$\frac{dX_3}{dt}=g_3X_3\left(1\text{-}\frac{X_1+X_2+X_3}{N_m}\right)\text{-}D_3X_3 \qquad [S3.27]$$

Since all populations can only use the end product P as the sole carbon source, the growth rate of the three populations, $g_1$, $g_2$ and $g_3$, are calculated by,

$$g_1=\frac{kg_1P_{1,in}}{Kg_1+P_{1,in}}Y_1c_1 \qquad [S3.28]$$

$$g_2=\frac{kg_2P_{2,in}}{Kg_2+P_{2,in}}Y_2c_2 \qquad [S3.29]$$

$$g_3=\frac{kg_3P_{3,in}}{Kg_3+P_{3,in}}Y_3c_3 \qquad [S3.30]$$

The model was simplified following the same methods as described in Section S1.1.2, and then non-dimensionalized following the method described in Table S4 and Table S5, so obtain the dimensionless cell growth equations for the three MDOL members.

$$\frac{dx_1}{d\tau}=\mu_1x_1\left(1\text{-}\frac{x_1+x_2+x_3}{\rho}\right)\text{-}d_1x_1 \qquad [S3.31]$$

$$\frac{dx_2}{d\tau}=\mu_2x_2\left(1\text{-}\frac{x_1+x_2+x_3}{\rho}\right)\text{-}d_2x_2 \qquad [S3.32]$$

$$\frac{dx_3}{d\tau}=\mu_3x_3\left(1\text{-}\frac{x_1+x_2+x_3}{\rho}\right)\text{-}d_3x_3 \qquad [S3.33]$$

$$\mu_1=Ig_1p_{1,in}y_1c_1 \quad [S3.34]$$

$$\mu_2=Ig_2p_{2,in}y_2c_2 \quad [S3.35]$$

$$\mu_3=Ig_3p_{3,in}y_3c_3 \quad [S3.36]$$

### S3.2 Derivation of the assembly rule of a three-step MDOL community

#### *S3.2.1 Derivation of the conditions required for the stability of a three-step MDOL community*

To derive the simple formula that defines the basic assembly rule of a three-step MDOL community, we solve the steady-state equations of ODE systems containing Eqns. [S3.13-3.24] and Eqns. [S3.31-3.36]. However, the steady-state equations system has an infinite number of solutions, which may be due to the initial conditions (such as the initially inoculating ratio of different populations, see Section S5 for further discussion). Alternatively, three equations regarding the steady-state biomass of the three populations were obtained from the analyses of the steady-state equations

$$\frac{a_1c_1y_1n1(n2+1)x_1\left(1-\frac{x_1+x_2+x_3}{\rho}\right)}{(n3+1)(n1(n2+1)x_1+n2(n1+1)x_2)+n3(n1+1)(n2+1)x_3}=d_1 \quad [S3.37]$$

$$\frac{a_1c_2y_2n2(n1+1)x_1\left(1-\frac{x_1+x_2+x_3}{\rho}\right)}{(n3+1)(n1(n2+1)x_1+n2(n1+1)x_2)+n3(n1+1)(n2+1)x_3}=d_2 \quad [S3.38]$$

$$\left(1+\frac{(n1+1)(n2+1)x_3}{(n3+1)(n1(n2+1)x_1+n2(n1+1)x_2)+n3(n1+1)(n2+1)x_3}\right)=\frac{d_3(n3+1)}{a_1c_3y3n3x_1\left(1-\frac{x_1+x_2+x_3}{\rho}\right)} \quad [S3.39]$$

Here,

$$n_1=\frac{Ig_1}{\gamma_p} \quad [S3.40]$$

$$n_2=\frac{Ig_2}{\gamma_p} \quad [S3.41]$$

$$n_3=\frac{Ig_3}{\gamma_p} \qquad [S3.42]$$

Deriving from Eqn. [S3.37] and Eqn. [S3.38], we obtained a formula that defines a prerequisite for the co-existence of three populations

$$\frac{n_1+1}{m_1+1}=\frac{n_2+1}{m_2+1} \qquad [S3.43]$$

$$m_1=\frac{\frac{c_1Ig_1y_1}{d_1}-\frac{c_3Ig_3y_3}{d_3}}{\frac{c_3Ig_3y_3}{d_3}} \qquad [S3.44]$$

$$m_2=\frac{\frac{c_2Ig_2y_2}{d_2}-\frac{c_3Ig_3y_3}{d_3}}{\frac{c_3Ig_3y_3}{d_3}} \qquad [S3.45]$$

Similar as the definitions in Section S1.3, $n_1$, $n_2$ is the 'Product demand gap' of the [1, 0, 0] and [0, 1, 0], respectively, reflecting the relative 'private benefit' of the population performing the last step against the corresponding population. In addition, $\frac{c_1Ig_1y_1}{d_1}$, $\frac{c_2Ig_2y_2}{d_2}$, and $\frac{c_3Ig_3y_3}{d_3}$ reflect the inherent net growth rate of [1, 0, 0], [0, 1, 0], and [0, 0, 1], respectively. Thus, $m_1$ indicates the difference between the inherent growth rates of [1, 0, 0] and [0, 0, 1], normalized by the inherent growth rate of [0, 0, 1], while $m_2$ is the similar index derived from the difference between [1, 0, 0] and [0, 0, 1]. In particular,

$$m_1+1=\frac{\frac{c_1Ig_1y_1}{d_1}}{\frac{c_3Ig_3y_3}{d_3}} \qquad [S3.46]$$

$$m_2+1=\frac{\frac{c_2Ig_2y_2}{d_2}}{\frac{c_3Ig_3y_3}{d_3}} \qquad [S3.47]$$

Therefore, $m_1+1$ quantifies the ratio of the inherent net growth rate of [1, 0, 0] to that of [0, 0, 1], reflects the inherent growth advantage of [1, 0, 0] to [0, 0, 1]. Adding that

$n_1$ reflects the relative 'private benefit' of [0, 0, 1] against [1, 0, 0], $\frac{n_1+1}{m_1+1}$ indicates inherent growth advantage of [1, 0, 0] divides the advantage of [0, 0, 1] from product privatization, reflecting the summarized relative fitness between [1, 0, 0] and [0, 0, 1]. Similarly, $\frac{n_2+1}{m_2+1}$ reflects the relative fitness between [0, 1, 0] and [0, 0, 1]. Therefore, Eqn. [S3.43] indicates a prerequisite for the co-existence of three populations that the relative fitness between [1, 0, 0] and [0, 0, 1] should be equal to that of [0, 1, 0] and [0, 0, 1]. In particular, when the 'Product demand gap' of the [1, 0, 0] and [0, 1, 0] is equal, that is, $n_1=n_2$, we obtain $m_1=m_2$, meaning that the two population must hold same inherent relative fitness.

To test this rule, we searched 308913 parameter sets in which the values of $n_1, n_2, m_1$, and $m_2$ were set in a range covering 0.1~10. In these parameter sets, 2783 sets satisfy the condition defined by Eqn. [S3.43]. Numeric simulations initialized with these parameter sets indicate that 890 parameter sets leads to the co-existence of three populations (that is, exhibiting stable community dynamics), and these sets are all belong to the 2783 sets that satisfy Eqn. [S3.43] (Figure S16B). This result confirms that the Eqn. [S.43] is the necessary condition for the stability of a three-step MDOL community, but not sufficient.

We next investigated whether there are other conditions required for the stability of a three-step MDOL community. Suppose that

$$\frac{n_1+1}{m_1+1}=\frac{n_2+1}{m_2+1}=ra \qquad [S3.48]$$

Since that

$$n_1 > 0 \quad [S3.49]$$

$$m_1 = \frac{\frac{c_1 Ig_1 y_1}{d_1} - \frac{c_3 Ig_3 y_3}{d_3}}{\frac{c_3 Ig_3 y_3}{d_3}} > -1 \quad [S3.50]$$

$$n_2 > 0 \quad [S3.51]$$

$$m_2 = \frac{\frac{c_2 Ig_2 y_2}{d_2} - \frac{c_3 Ig_3 y_3}{d_3}}{\frac{c_3 Ig_3 y_3}{d_3}} > -1 \quad [S3.52]$$

We then obtained

$$-\frac{(-1+ra)\frac{n_1}{m_1}}{ra-\frac{n_1}{m_1}} > 0 \quad [S3.53]$$

$$-\frac{-1+ra}{ra-\frac{n_1}{m_1}} > -1 \quad [S3.54]$$

$$-\frac{(-1+ra)\frac{n_2}{m_2}}{ra-\frac{n_2}{m_2}} > 0 \quad [S3.55]$$

$$-\frac{-1+ra}{ra-\frac{n_2}{m_2}} > -1 \quad [S3.56]$$

Solving these inequalities (using Reduce function of *Wolfram Mathematica*) offered us three possible regions for the values of $\frac{n_1}{m_1}$ and $\frac{n_2}{m_2}$

(1) $0 < \frac{n_1}{m_1} < ra < 1,\ 0 < \frac{n_2}{m_2} < ra < 1$

(2) $ra > 1,\ \frac{n_1}{m_1} < 0,\ \frac{n_2}{m_2} < 0$

(3) $1 < ra < \frac{n_1}{m_1},\ 1 < ra < \frac{n_2}{m_2}$

To test whether all these three conditions can lead to stable community dynamics, we designed 62500 parameter sets (design of each set also follows Eqn. [S3.43]) that meet each condition and searched for the parameter sets that leads to the co-existence of the three populations. Notably, for the simulations using the 125000 parameter sets that follows the second and third conditions, no parameter sets leads to stable community dynamics. In contrast, when the values of $\frac{n_1}{m_1}$ and $\frac{n_2}{m_2}$ follows the first condition, 35938 of 62500 parameter sets leads to the co-existence of the three populations. Therefore, the condition

$$0 < \frac{n_1}{m_1} < ra < 1 \qquad [S3.57]$$

$$0 < \frac{n_2}{m_2} < ra < 1 \qquad [S3.58]$$

is another prerequisite required for the stability of a three-step MDOL community, which indicates that the populations [1, 0, 0] and [0, 1, 0] should hold a growth advantage that outweighs the 'private benefit' of the [0, 0, 1]. In summary, we proposed that two prerequisites, as defined by Eqn. [S3.43] and Eqns. [S3.57-3.58], are required for the stability of a three-step MDOL community.

*S3.2.2 Derivation of the formula predict the assembly of a three-step MDOL community*

We next focused on the conditions that can leads to the stability of a three-step MDOL community and deduced the mathematical rule that can predict the relative abundance of the populations in the stable three-step MDOL communities. We first analyzed the results of the above-mentioned simulations using the 890 parameter sets. We proposed a formula to predict the relative abundance of [0, 0, 1] ($R_{[0,\,0,\,1]}$) with the parameters $n_1$, $n_2$, $m_1$, and $m_2$, given by

$$R_{[0,\,0,\,1]} = \sqrt[h]{\frac{(\frac{n_1}{m_1})^h + (\frac{n_2}{m_2})^h}{2}} \qquad [S3.59]$$

We fitted the results from the 890 parameter sets to Eqn. [S3.59] (NonlinearModelFit function of *Wolfram Mathematica*), and found this formula can well predict the community assembly in these simulations (Figure 4B). Here, the best fitting of the mean square index ($h$) is 2.40, and the correlation coefficient R2 equals 0.997. Therefore, Eqn. [S3.59] suggests a simple formula to predict the assembly of a three-step MDOL community.

To further test the predicting power of this formula, we designed 72030 parameter sets, which follow the condition defined by Eqn. [S3.43] and considers gradients of $ra$ and $n_3$ values. As expected, the simulation results indicate that stability of the three-step communities requires the parameter sets follow the perquisite defined by Eqns. [S3.57-3.58], and importantly, Eqn. [S3.59] can well predict the abundance of [0, 0, 1] (Figure S3.1A). In addition, we also found that with the increase of $n_3$, the size of the parameter space decreases (Figure S3.1B), suggesting that the value of $n_3$ also affect the stability of a MDOL community. Moreover, the best fitting value of index $h$ is largely influenced by the value of $ra$ and $n_3$. With the increase of $ra$ or decrease of $n_3$, the value of $h$ increases, ranging from 1 to 4 (Figure S3.1C), suggesting that the value of $ra$ and $n_3$ also affect the assembly of a stable three-step MDOL community. Finally, while these simulations were initialized with equal inoculating ratio of the three populations, the frequencies of [1, 0, 0] and [0, 1, 0] in the final stable community are very close (nearly equals 1:1 ratio; Figure S3.1D). Actually, the frequencies of [1, 0, 0] and [0, 1, 0] are largely affected by the initial ratios of the populations, which we will further discussed in Section S5.

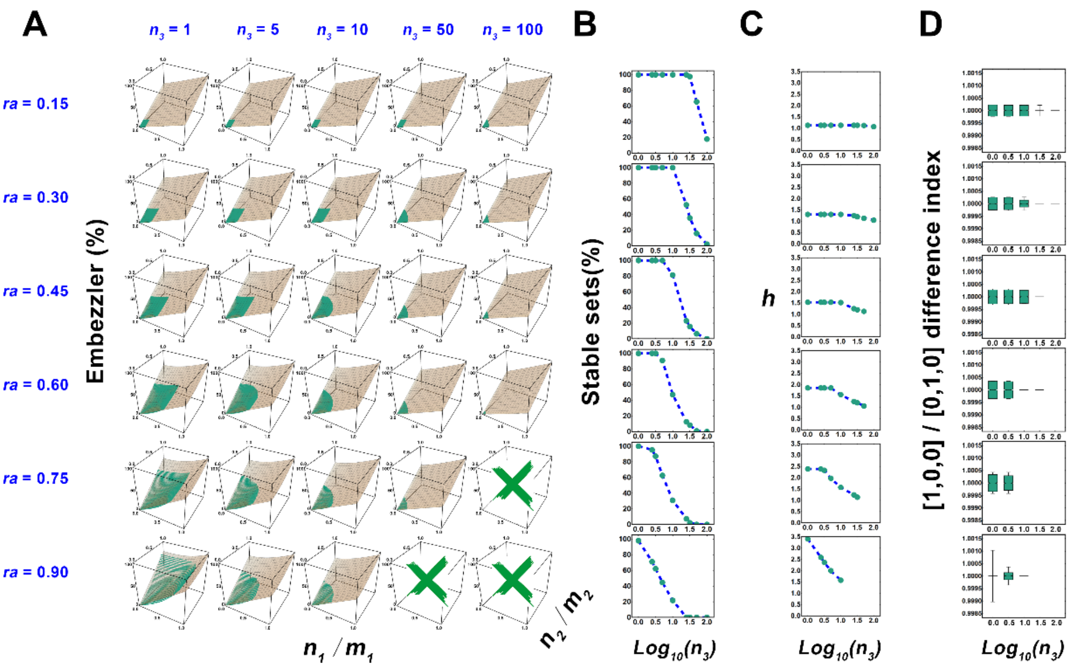

**Figure S3.1** Effects of $ra$ and $n_3$ on the stable criteria and steady-state community structure of a three-step MDOL community. (A) The range of stable parameter sets characterized by ratios $\frac{n_1}{m_1}$ and $\frac{n_2}{m_2}$, as well as relative abundance of embezzler population under different $ra$ and $n_3$ combinations. The scatters show the relative abundance of the embezzler population obtained by simulations different parameter sets. The surface diagram shows the relative abundance distribution of embezzler population predicted by Eqn. [S3.59]. The green cross indicates that there are no $\frac{n_1}{m_1}$ and $\frac{n_2}{m_2}$ combinations that can stabilize the community in the parameter set under the corresponding $ra$ and $n_3$. (B) The shift of the ratio of parameter sets that can stabilize the community with $n_3$ under the same $ra$ value. (C) The shift of the optimal mean square index $h$ fitted according to Eqn. [S3.59] with $n_3$ under the same $ra$ value. (D) The shift of the distribution of relative proportional index with $n_3$ under the same $ra$ value. Because the values are so concentrated, the upper and lower quantiles are set to

0.999 and 0.001 respectively when plotting the boxplot. Except for parameters related to the parameter set generation, the default values of other parameters are given in Table S5.

*S3.2.3 Sensitive analyses of the assembly rule of a three-step MDOL community*

For the sensitive analyses of the proposed rule, we performed numeric simulations to test how the reaction rates (that is $a_1$, $a_2$ and $a_3$), and maximum biomass capacity ($\rho$) affect the proposed rule for the assembly of a three-step MDOL community. For the test of each parameter, 2401 parameter sets that follows the condition defined by Eqn. [S3.43] and Eqns. [S3.57-3.58] were first generated. Then the values of the tested parameter were varied, and numerous simulations were performed with each set. Other parameters were set as the default value shown in Table S5.

Our test of the effects of reaction rates indicates that the shifts of the reaction rates of the second and third step ($a_2$ and $a_3$) rarely changed our rule regarding the assembly of a three-step MDOL community (Figure S3.2A-C). In contrast, the reaction rate of the first step ($a_1$) largely affect the assembly rule, in which the lower $a_1$ would shrink the parameter space that defines when a three-step MDOL community are stable (Figure S3.2D). Therefore, the stability of a three-step MDOL community also required the reaction rate of the first step over a threshold. Moreover, the reaction rate of the first step also affect the relative abundance of [0, 0, 1] in the final stable community. As shown in (Figure S3.2E), with the increase of $a_1$, the best fitting value of $h$ increases. When the value of $a_1$ large enough, the value of $h$ reached a maximum value of approximately 3.40 and no longer changed with the increase of $a_1$ and $n_3$. Therefore, the reaction rate of the first step affect the relative abundance of [0, 0, 1] by varying the value of the index $h$.

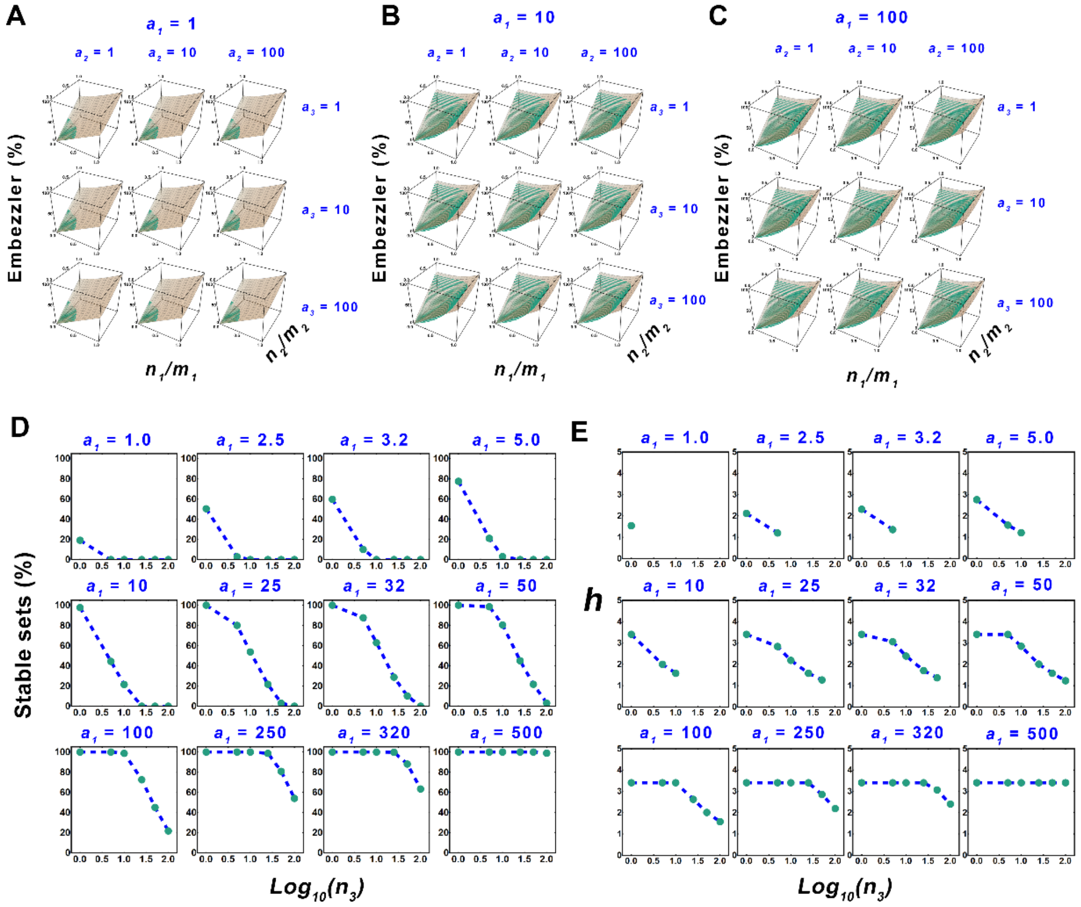

**Figure S3.2** Effects of reaction rates of each step on stable criteria and steady-state community structure of a three-step MDOL community. (A-C) The range of stable parameter sets characterized by ratios $\frac{n_1}{m_1}$ and $\frac{n_2}{m_2}$, as well as relative abundance of embezzler population under different metabolic reaction rate combinations ($a_1$, $a_2$ and $a_3$) . The scatters show the relative abundance of the embezzler population obtained by simulations different parameter sets. The surface diagram shows the relative abundance distribution of embezzler population predicted by Eqn. [S3.59]. (D) The shift of the ratio of parameter sets that can stabilize the community with $n_3$ under different first-step reaction rate ($a_1$). (E) The shift of the optimal mean square index $h$ fitted according to Eqn. [S3-48] with $n_3$ under different first-step reaction rate ($a_1$). Except for parameters related to the parameter set generation, default values of other parameters are given in Table S5.

Our further analyses indicated that the shift of the maximum biomass capacity ($\rho$) rarely changed the size of the parameter space required for the stability of a three-step MDOL community (Figure S3.3A), as well as the best fitting value of $h$ (Figure S3.3B). Therefore, our assembly rule of a three-step MDOL community is robust to the shift of the maximum biomass capacity. Notably, the results of these analyses are similar as those for a two-step MDOL community (Section S1.4).

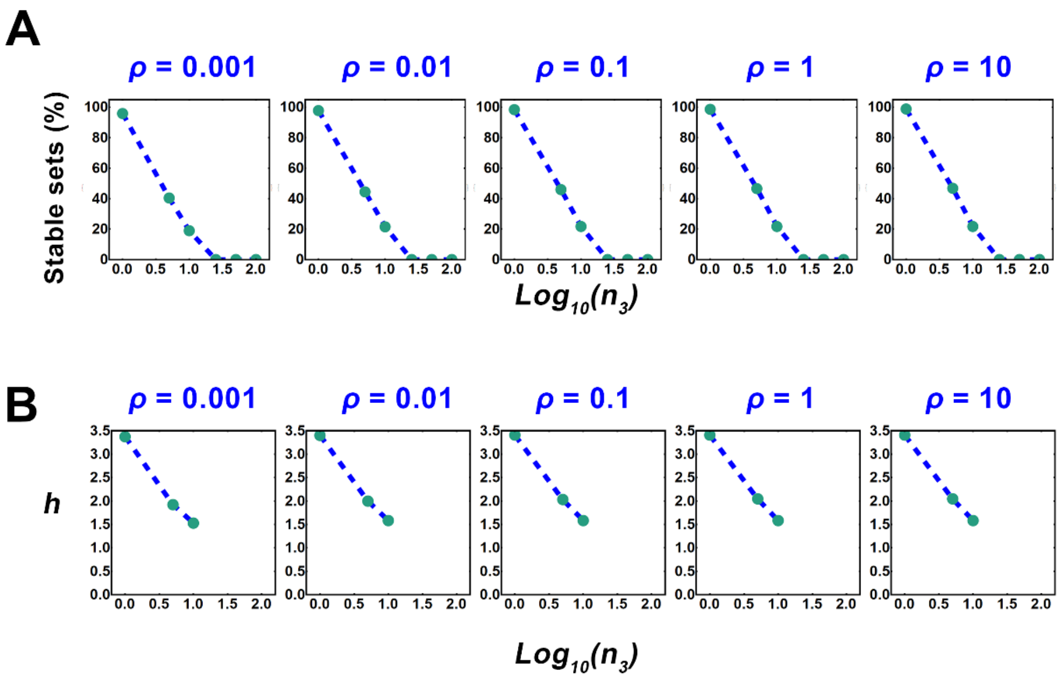

**Figure S3.3** Effects of carrying capacity of the whole populations on stable criteria and steady-state community structure of a three-step MDOL community. (A) The shift of the ratio of parameter sets that can stabilize the community with $n_3$ under different carrying capacity ($\rho$). (B) The shift of the optimal mean square index $h$ fitted according to Eqn. [S3.59] with $n_3$ under under different carrying capacity ($\rho$). Except for parameters related to the parameter set generation, the default values of other parameters are given in Table S5.

**S3.3 Derivation of the model regarding the assembly of a *N*-step MDOL community**

After deriving the rule governing the assembly of a three-step MDOL community, we further generalized the rule to predict the assembly of a MDOL community that executes a degradation pathway by dividing it into *N*-steps (Figure 4A). Totally *N* populations are included in this community. The community converts a substrate (S) to the end product (P) through *N*-1 intermediates, and the *j*th intermediate was labelled with $I^j$. Each reaction can be only carried out by one population, which is represented by a vector

$$\varepsilon_k=\left[\epsilon_{k,1},\ \epsilon_{k,2},\ \ldots,\ \epsilon_{k,k},\ \ldots,\ \epsilon_{k,N-1},\ \epsilon_N\right] \qquad [S3.60]$$

to characterize the phenotype of each population, in which

$$\epsilon_{k,j}=1\ (j=k),\ \ \epsilon_{k,j}=0\ (j\neq k) \qquad [S3.61]$$

Here, 1 denotes that the corresponding population is capable of executing the *k*th step, and 0 denotes it cannot. Following these assumptions, we expanded the ODE model of the two-step and a three-step MDOL community to construct the general model regarding the dynamics of a *N*-step MDOL community. Construction of this model still follows the seven biological assumptions listed in Section S1.

*S3.3.1 Equations regarding Intermediates and product dynamics*

We first built a system of ODEs describing dynamics of Intermediates and product concentrations for an intracellular pathway in a *N*-step MDOL community. The model is simplified by the same assumptions as described in Section S1, given by

$$\frac{dI_{k,in}^{1}}{dt}\cdot V_c=\epsilon_{k,1}k_1E_1-r_{I1}\cdot V_c\cdot\left(I_{k,in}^{1}-I_{out}^{1}\right) \quad [S3.62]$$

$$\frac{dI_{k,in}^{j}}{dt}\cdot V_c=\epsilon_{k,j}\frac{k_kE_k}{K_k}I_{k,in}^{j-1}-\epsilon_{k,j+1}\frac{k_{k+1}E_{k+1}}{K_{k+1}}I_{k,in}^{j}+r_{I1}\cdot V_c\cdot\left(I_{out}^{j}-I_{k,in}^{j}\right) \quad (j>1) \quad [S3.63]$$

$$\frac{dP_{k,in}}{dt}\cdot V_c=\epsilon_{k,N}\frac{k_{N-1}E_{N-1}}{K_{N-1}}I_{k,in}^{N-1}-\frac{kg_k}{Kg_k}P_{k,in}+r_P\cdot V_c\cdot\left(P_{out}-P_{k,in}\right) \quad [S3.64]$$

$$\frac{dI_{out}^{j}}{dt}=-\sum_{k=1}^{N}X_k\cdot r_{I^j}\cdot\left(I_{out}^{j}-I_{k,in}^{j}\right) \quad [S3.65]$$

$$\frac{dP_{out}}{dt}=-\sum_{k=1}^{N}X_k\cdot r_p\cdot\left(P_{out}-P_{k,in}\right) \quad [S3.66]$$

The equations were then non-dimensionalized following the method described in Table S4 and Table S5, which formalized a non-dimensionalized model given by Eqns. [18]-[22] in the main text.

*S3.3.2 Equations regarding population growth dynamics*

Expanding the basic equations described in Section S1.1.2, our model then accounts for the growth dynamics of the populations in a *N*-step MDOL community.

$$\frac{dX_k}{dt}=g_kX_k\left(1-\frac{\sum_{k=1}^{N}X_k}{N_m}\right)-D_kX_k \quad [S3.67]$$

Since all populations can only use the end product P as the sole carbon source, the growth rate of the *k*th population ($\boldsymbol{g_k}$), are calculated by,

$$g_k=\frac{kg_kP_{k,in}}{Kg_K+P_{k,in}}Yc_k \quad [S3.68]$$

The model was simplified following the same methods described in Section S1.1.2, and then non-dimensionalized following the method described in Table S4 and Table S5, so obtain the dimensionless cell growth equation given by Eqn. [23] in the main text.

### S3.4 Derivation of the assembly rule of a *N*-step MDOL community

#### *S3.4.1 Derivation of the conditions required for the stability of a N-step MDOL community*

To derive the conditions required for the stability of a *N*-step MDOL community, we analyzed the ODE systems containing Eqns. [18-23], using the similar method as the analyses for a three-step MDOL community (Section S3.2.1). By analyzing the steady-state equations of the ODE system using the same way as Section S3.2.1, we obtained the first condition similar as the one defined by Eqn. [S3.43], as shown in Eqn. [3] in the main text. Note that

$$n_k = \frac{Ig_k}{\gamma_p} \qquad [S3.69]$$

$$m_k = \frac{\frac{c_k Ig_k y_k}{d_k} - \frac{c_N Ig_N y_N}{d_N}}{\frac{c_N Ig_N y_N}{d_N}} \qquad [S3.70]$$

Therefore, $n_k$ is the 'Product demand gap' of the *k*th population, reflecting the relative 'private benefit' of the population performing the last step against the *k*th population. In addition, $\frac{c_k Ig_k y_k}{d_k}$ reflects the inherent net growth rate of the *k*th population, while $\frac{c_N Ig_N y_N}{d_N}$ reflects the inherent net growth rate of the population performing the last step. Thus, $m_k$ indicates the difference between the inherent growth rates of the *k*th population and the last population, normalized by the last population. In particular,

$$m_k+1=\frac{\frac{c_k Ig_k y_k}{d_k}}{\frac{c_N Ig_N y_N}{d_N}} \qquad [S3.71]$$

which suggests that $m_k+1$ quantify the ratio of the inherent net growth rate of the *k*th population to that of the last population. Adding that $n_k$ reflects the relative 'private benefit' of the last population against the *k*th population, $\frac{n_k+1}{m_k+1}$ indicates inherent growth advantage of the *k*th population divides the advantage of the last population from product privatization, reflecting the summarized relative fitness between *k*th population and the last population. Therefore, Eqn. [S3.71] indicates a prerequisite for the stability of the community engaged in *N*-step MDOL that the relative fitness between the former population and the last population should be equal.

In particular, when the 'Product demand gap' of the former populations is totally equal, that is, $n_1 = n_2 = ... = n_N$, we obtain $m_1 = m_2 = ... = m_N$, meaning that all the former populations must hold same inherent relative fitness. This scenario matched with our synthetic community engaged in four-step MDOL, since all the strains are derived from a same ancestral strain, possess similar maximum product consuming rate ($Ig_k$) and product diffusive coefficient, and thus hold same 'Product demand gap' ( $n_k$).

We next generalized the condition defined by Eqn. [S3.56-57] to explain the condition required for the stability of *N*-step community. similar as Section S3.2.1, we supposed that

$$\frac{n_1+1}{m_1+1}=\frac{n_2+1}{m_2+1}=...=\frac{n_k+1}{m_k+1}=...=\frac{n_{N-1}+1}{m_{N-1}+1}=ra \qquad [S3.72]$$

Since that

$$n_k > 0 \qquad [S3.73]$$

$$m_k = \frac{\frac{c_k I g_k y_k}{d_k} - \frac{c_N I g_N y_N}{d_N}}{\frac{c_N I g_N y_N}{d_N}} > -1 \qquad [S3.73]$$

We then obtained

$$-\frac{(-1+ra)\frac{n_k}{m_k}}{ra-\frac{n_k}{m_k}} > 0 \qquad [S3.74]$$

$$-\frac{-1+ra}{ra-\frac{n_k}{m_k}} > -1 \qquad [S3.75]$$

Solving the two inequalities (using Reduce function of *Wolfram Mathematica*) again suggested us three possible regions for the values of $\frac{n_k}{m_k}$

(1) $0 < \frac{n_k}{m_k} < ra < 1$

(2) $ra > 1$, $\frac{n_k}{m_k} < 0$

(3) $1 < ra < \frac{n_k}{m_k}$

We then tested whether the values of $\frac{n_k}{m_k}$ located in these regions would lead to stable community dynamics by performing simulations regarding four-, five-, six-, seven-, and eight-step MDOL systems. For each community set, we designed at least 300000 parameter sets (design of each set follows Eqn. [3]) for possible region and searched for the parameter sets that leads to the stable community dynamics. Again, we found that only when the values of $\frac{n_k}{m_k}$ are located in the first region, did the parameter sets (190082 of 345600 parameter sets for a four-step MDOL community; 197556 of 366025 parameter sets for a five-step MDOL community; 218488 of 420175 parameter sets for a six-step MDOL community; 239048 of 468720 parameter sets for a seven-step MDOL community; 342098 of 589824 parameter sets for an eight-step

MDOL community) leaded to the co-existence is present. Therefore, the condition defined by the first region, as shown in Eqn. [5] of the main text, is the second general prerequisite required for the stability of a *N*-step MDOL community. The condition indicates that in a multi-step MDOL community, the populations performing the former step should hold a growth advantage that outweighs the 'private benefit' of the one performing the last step. Taken together, we generalized the two prerequisites defined for the stability of a three-step MDOL community to predict the scenarios of a *N*-step MDOL communities, as defined by Eqn. [4] and Eqn. [5].

*S3.4.2 Derivation of the formula predict the assembly of a N-step MDOL community*

In section S3.2.2, we proposed a simple formula (Eqn. [S3.59]) that can predict the relative abundance of the population performing the last step (population [0, 0, 1]) in a stable three-step MDOL communities. we hypothesized that this rule can be generalized to predict the steady-state relative abundance of the population performing the last step in a *N*-step MDOL community, as given by Eqn. [6] in the main text.

We tested this hypothesis in the four-, five-, six-, seven-, and eight-step MDOL communities. For each community set, we designed at least 13000 parameter sets that follows the condition defined by Eqn. [4] and Eqn. [5]. As expected, simulations with these parameter sets indicates that relative abundances of the last population obtained from simulations matched well with the ones calculated from Eqn. [6] (Figure 4C), suggesting that our rule can be generalized to explain the assembly of a *N*-step MDOL community (at least, it was applicable to a MDOL community in which the pathway is distributed among ≤ 8 steps).

Moreover, we also set the four-, five-, and six-step MDOL communities as examples to investigate the effects of $ra$ and $n_N$ (where $n_N=Ig_N/\gamma_p$) on the assembly rule. We found that with the increase of $n_N$, the size of the parameter space decreases (Figure S3.3-S3-5B), suggesting that the value of $n_N$ also affect the stability of a $N$-step MDOL community. Moreover, the best fitting value of index $h$ is also largely influenced by the value of $ra$ and $n_N$. With the increase of $ra$ or decrease of $n_N$, the value of $h$ increases (ranging from 1.0 to 4.0 in a four-step MDOL community; from 1.0 to 3.0 in the five- and six-step MDOL communities; Figure S3.3-S3-5C), suggesting that the values of $ra$ and $n_N$ also affect the assembly of a stable $N$-step MDOL community.

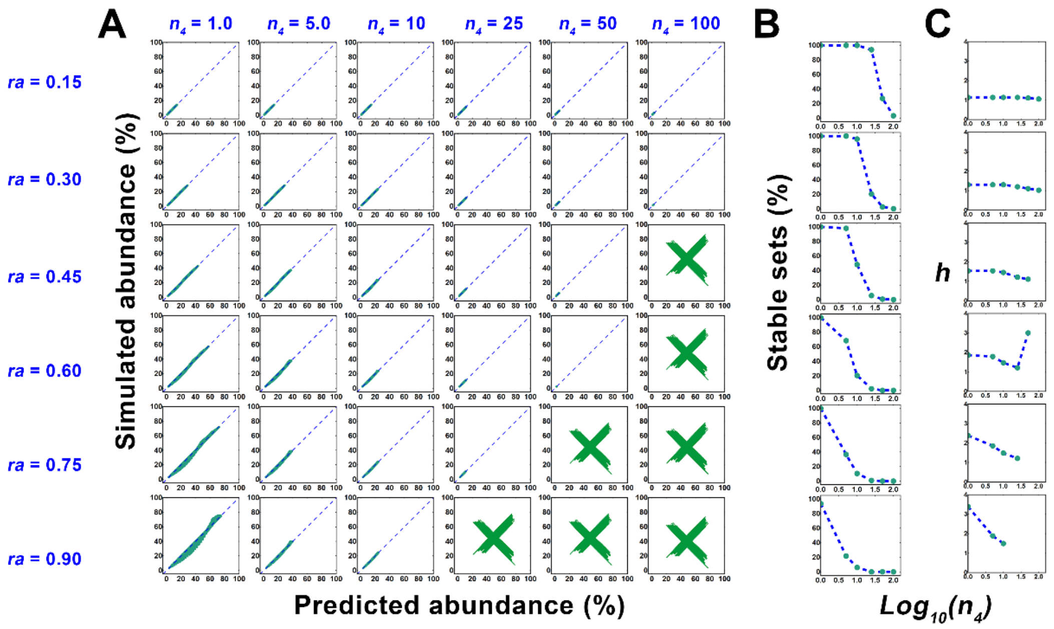

**Figure S3.3** The effects of $ra$ and $n_4$ on the stable criteria and steady-state community structure of the four-step metabolic division community. (A) The linear correlation between the abundance of embezzler [0, 0, 0, 1] predicted by Eqn. [6] and the abundance of embezzler population obtained by simulations of four-step MDOL system under different $ra$ and $n_4$ combinations. The scatters show the distribution of relative abundance and corresponding predicted values of the embezzler population obtained by simulations with different parameter sets. The dashed line shows the fitting correlation curve. Green cross indicates that under the corresponding $ra$ and $n_4$, there is no parameter combination can stabilize the community. (B) The shift of the ratio of parameter sets that can stabilize the community with different $ra$ and $n_4$. (C) The shift of the optimal mean square index $h$ fitted according to Eqn. [6] with different $ra$ and $n_4$. Except for the parameters related to parameter set generation, the default values of all other parameters are given in Table S5.

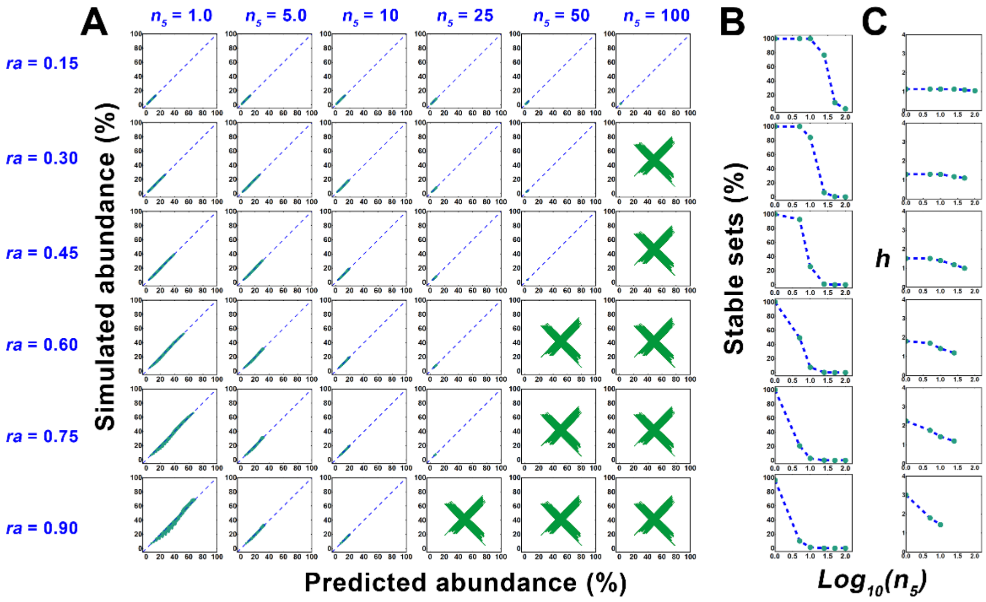

**Figure S3.4** The effects of $ra$ and $n_5$ on the stable criteria and steady-state community structure of the five-step metabolic division community. (A) The linear correlation between the abundance of embezzler [0, 0, 0, 0, 1] predicted by Eqn. [6] and the abundance of embezzler population obtained by simulations of four-step MDOL system under different $ra$ and $n_5$ combinations. The scatters show the distribution of relative abundance and corresponding predicted values of the embezzler population obtained by simulations with different parameter sets. The dashed line shows the fitting correlation curve. Green cross indicates that under the corresponding $ra$ and $n_5$, there is no parameter combination can stabilize the community. (B) The shift of the ratio of parameter sets that can stabilize the community with different $ra$ and $n_5$. (C) The shift of the optimal mean square index $h$ fitted according to Eqn. [6] with different $ra$ and $n_5$. Except for the parameters related to parameter set generation, the default values of all other parameters are given in Table S5.

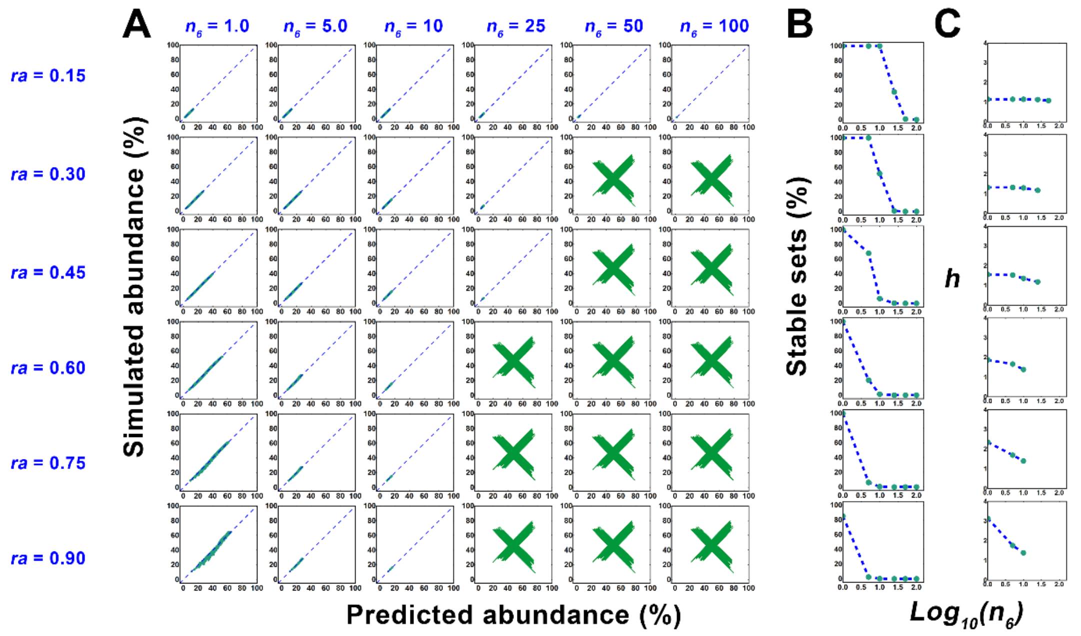

**Fig S3-5** The effects of $ra$ and $n_6$ on the stable criteria and steady-state community structure of the six-step metabolic division community. (A) The linear correlation between the abundance of embezzler [0, 0, 0, 0, 0, 1] predicted by Eqn. [6] and the abundance of embezzler population obtained by simulations of four-step MDOL system under different $ra$ and $n_6$ combinations. The scatters show the distribution of relative abundance and corresponding predicted values of the embezzler population obtained by simulations with different parameter sets. The dashed line shows the fitting correlation curve. Green cross indicates that under the corresponding $ra$ and $n_6$, there is no parameter combination can stabilize the community. (B) The shift of the ratio of parameter sets that can stabilize the community with different $ra$ and $n_6$. (C) The shift of the optimal mean square index $h$ fitted according to Eqn. [6] with different $ra$ and $n_6$. Except for the parameters related to parameter set generation, the default values of all other parameters are given in Table S5.

*S3.4.3 Sensitive analyses of the assembly rule of a N-step MDOL community*

Inspired by the previous results (Section S1.4 and Section 3.2.3), we performed numeric simulations to test how the rate of first reaction, and maximum biomass capacity ($\boldsymbol{\rho}$) affect the proposed the rule govern the assembly rule of a *N*-step MDOL community. For the test of each parameter, we set the four-, five-, and six-step MDOL communities as examples, and numerous parameter sets (13824 for a four-step MDOL community; 14641 for a five-step MDOL community; 16807 for a six-step MDOL community) that follows the condition defined by Eqn. [4] and Eqn. [5] were first generated. Then the values of the tested parameter were varied, and simulations were performed with each set. Other parameters were set as the default value shown in Table S5.

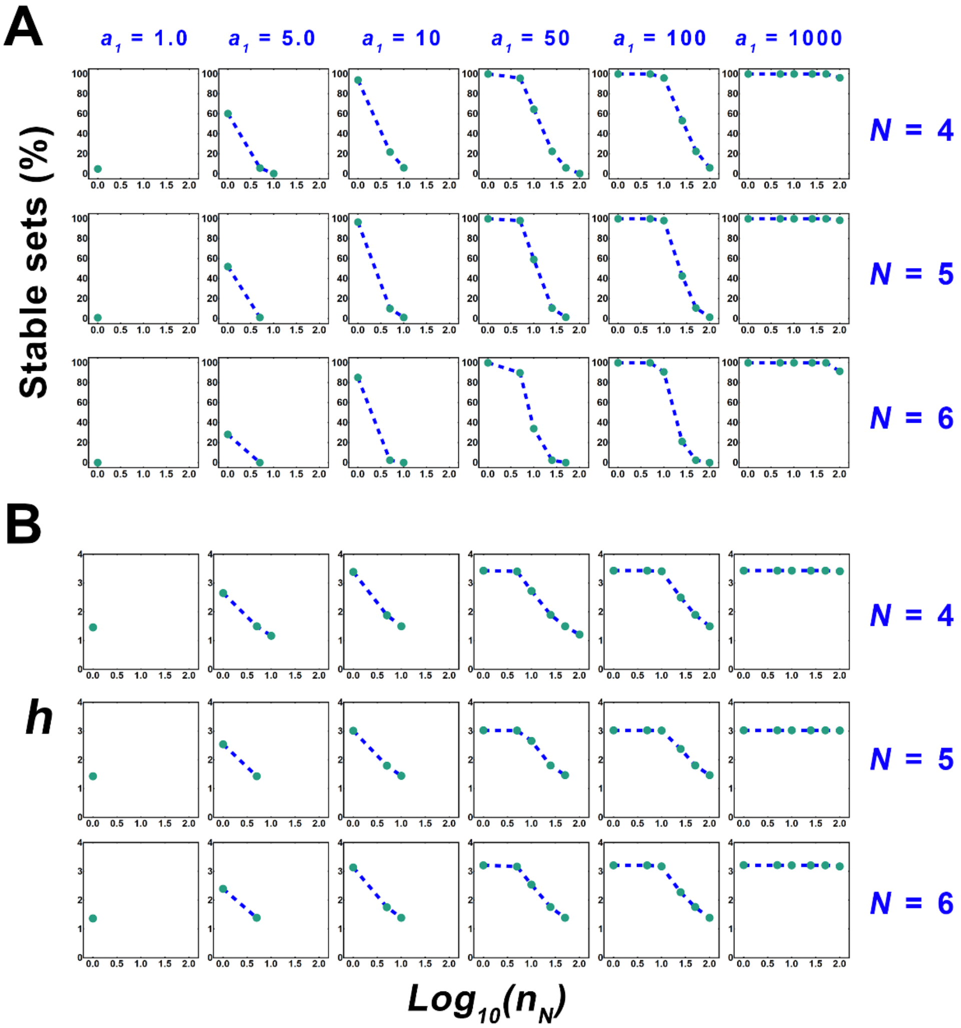

**Fig S3.6** The effects of first-step reaction rate ($a_1$) on the stable criteria and steady-state community structure of the four-, five- and six-step MDOL communities. (A) The shift of the ratio of parameter sets that can stabilize the community with different first-step reaction rate ($a_1$). (B) The shift of the optimal mean square index $h$ fitted according to Eqn. [6] with different first-step reaction rate ($a_1$). Except for parameters related to the parameter set generation, the default values of other parameters are given in Table S5.

Furthermore, our analyses also indicated that the shift of the maximum biomass capacity ($\rho$) rarely changed the size of the parameter space required for the stability of a $N$-step MDOL community (Figure S3.7A), as well as the best fitting value of $h$ (Figure S3.7B). Therefore, our generalized assembly rule is robust to the shift of the maximum biomass capacity.

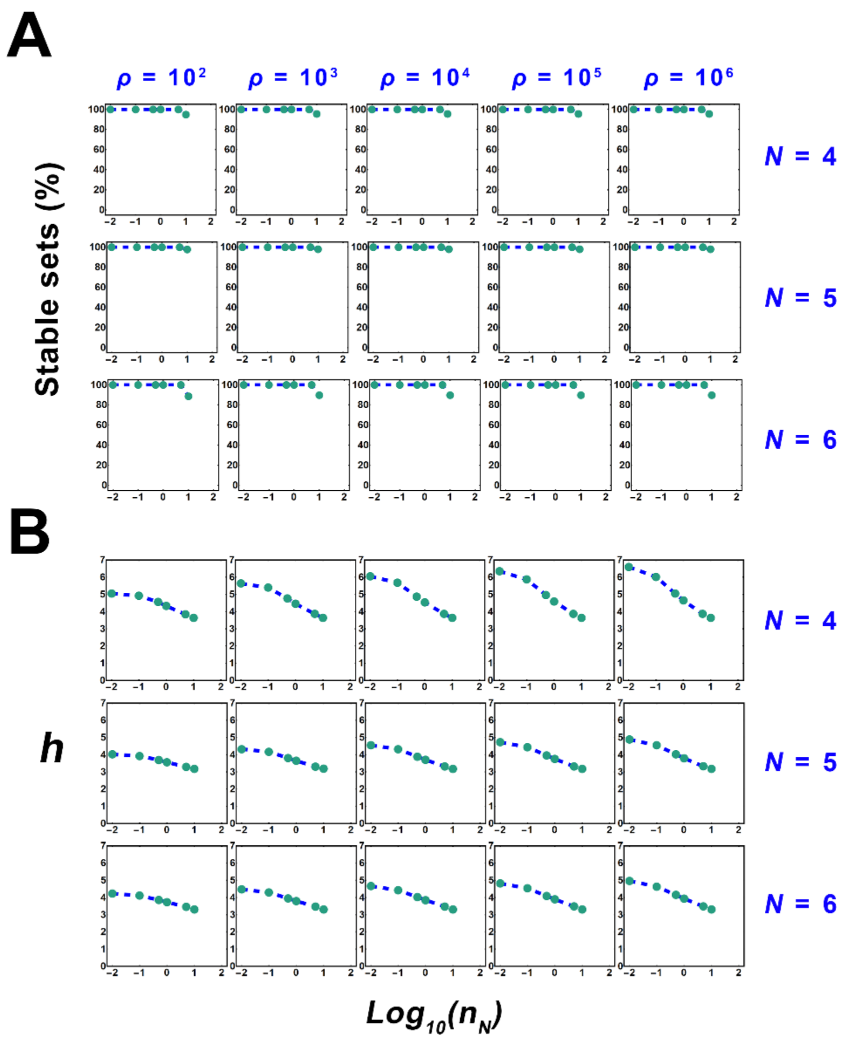

**Fig S3.7** The effects of Carrying capacity of the whole populations ($\rho$) on the stable criteria and steady-state community structure of four-, five- and six-step MDOL communities. (A) The shift of the ratio of parameter sets that can stabilize the community with different $\rho$. (B) The shift of the optimal mean square index $h$ fitted according to Eqn. [6] with different $\rho$. Except for parameters related to the parameter set generation, the default values of other parameters are given in Table S5.

Similar as the results about the assembly of a three-step MDOL community, we found that the reaction rate of the first step ($a_1$) largely affect the assembly rule, in which the lower $a_1$ would shrink the parameter space that defines when a $N$-step MDOL community is stable (Figure S3.6A). Therefore, the stability of a three-step MDOL community required the reaction rate of the first step over a threshold. $a_1$ also affect the relative abundance of last population in the final stable community. As shown in Figure S3.6B), with the increase of $a_1$, the best fitting value of $h$ increases. When the value of $a_1$ large enough, the value of $h$ reached a maximum value of approximately 3.43 for a four-step MDOL community, 3.02 for a five-step MDOL community, and 3.21 for a six-step MDOL community and no longer changed with the increase of $a_1$ and $n_N$. Therefore, the reaction rate of the first step affect the relative abundance of last population by varying the value of the index $h$.

**S3.5 Definitions and values of dimensionless variables and parameters**

The definition of all the variables included in the model regarding a three or $N$-step MDOL community, their corresponding dimensionless counterparts, and their value range during the numerical simulations are listed in Table S4. Meanwhile, the definition of all the parameters, their corresponding dimensionless methods, and their values for numerical simulations are listed in Table S5. Parameter values are chosen/calculated following the same protocol as described in section S1.2. To construct parameter sets, range of each focal parameter ranges are chosen to incorporate at least ~10-fold higher or lower than the default value and at least ~5 values with gradient were set for each parameter. When performing mathematical simulations, the initial values of the biomass of the all the populations was set to be equal as $x_{i,0}$ (values listed in Table S2), while the initial values of other variables were set to zero. Situations where the initial biomass were varied for different populations are discussed in Section S5.

**S3.6 Testing the effects of initial strain ratio on the assembly rule of a multi-step MDOL community**

In order to investigate how the initial ratio of different populations affect our proposed assembly rule of a multi-step MDOL community, we set the four-, five-, and six-step MDOL communities as examples. Assuming that $ra = 0.9$ and $n_N = 1.0$, numerous parameter sets (13824 for a four-step MDOL community; 14641 for a five-step MDOL community; 16807 for a six-step MDOL community) that follows the condition defined by Eqn. [4] and Eqn. [5] were first generated. Then different initial ratios were designed, in which the initial abundance of one population was changed (10%, 20%, 30%, 40%, 50%, 60%, 70%, 80%, 90%), while the initial abundance of other populations was controlled to be equal. In summary, 497664, 658845, and 907578 parameter sets were generated for the simulations regarding the four-, five-, and six-step MDOL communities, respectively. Other parameters were set as the default value shown in Table S5. After simulations, the stable abundance of the last population was collected, and compared with the values calculated by Eqn. [7] by linear fitting analyses using LinearModelFit function in *Wolfram Mathematica*.

### S3.7 Further discussions on the effects of initial strain ratio on the assembly rule of a MDOL community

Recently, several studies suggested that metabolic interdependency among different members increases the community robustness facing changing the initial member ratio[8,9]. In contrast, species competition promotes the emerging frequency of multi-stability of the communities under the condition of different initial ratios[10]. In our two-step MDOL, [1, 0] and [0, 1] developed into a strong interdependent pattern: [1, 0] provides intermediate metabolite (I) to [0, 1] to producing final product (P) and, in turn, [0, 1] supplies P to support [0, 1]. This interdependence contributes to increasing the robustness of a two-step MDOL community, which has been observed in previous studies[9]. In comparison, in a multi-step MDOL community, metabolic interdependency among the former populations is much weaker: direct and unidirectional metabolic exchanges only occur among the ($k$-$1$)th, $k$th and ($k$+$1$)th population. Moreover, these former populations must compete for the limiting resource (P) for their survival. The weaker metabolic interdependency and stronger resource competition in a multi-step MDOL community may result in lower robustness. Our experiments also suggested that a MDOL community initiated with more evenness strain ratio resulted in a more efficient community of organic compound degradation (Figure S28 C-D), resembling the observations in other model communities[11]. Therefore, to achieve both stability and functionality, it is critical to initiate a multi-step MDOL community at conditions characterized by high evenness.

### S4 Experiment verification of the proposed rule

We experimentally verified our rule regarding the assembly of a MDOL community by engineering several synthetic microbial communities, in which different engineered bacterial strains executing MDOL to degrade naphthalene. Here we described the detailed protocols of the construction and verifications of the strains involved in the synthetic communities, as well as the modification and parameterization of the mathematical model to predict the the community dynamics. The strains and plasmids used in this study were summarized in Table S6.

#### S4.1 Construction and culturing of the synthetic microbial communities engaged in MDOL

*S4.1.1 Construction of the 'super strain' that degrades naphthalene autonomously*

The strain used for experimental verification were all engineered from a *Pseudomonas stutzeri* strains AN10[12-14], which possesses a great ability to degrade naphthalene autonomously through a long pathway. Our preliminary design is to knock out its genes encoding the enzymes responsible for each metabolic step, aiming to construct the mutants that is only capable of performing a single metabolic step, and then co-culture these strains to execute MDOL. In strain AN10, the genes encoding the enzymes for naphthalene degradation were reported to mainly locate in two operons[13,14]. However, when we searched its genome, we found another two groups of genetic elements may also be included in the naphthalene degradation pathway, that is, a *nahW* gene encoding a salicylate hydroxylase[13,15,16] and a Cat gene cluster encoding enzymes for the degradation of catechol (one of the intermediates in naphthalene degradation). Therefore, we first knocked out the *nahW* gene and the *catA* gene encoding a catechol 1,2-dioxygenase[17]. Then we replaced the original promoters of the two operons (induced by one of the intermediates, salicylate[13,14]) with the IPTG induced promoter, $P_{tac}$[18,19]. The derived strain can degrade naphthalene autonomously, and its genes encoding the enzymes for naphthalene degradation all locate in the two engineered operons (Figure S4.1). Therefore, its ability of naphthalene degradation can be quantitatively controlled by modulating the addition of IPTG (Figure S4.2). We named this controllable 'super strain' as *P. stutzeri* AN1111, denoting that it can executing all the metabolic steps of naphthalene degradation.

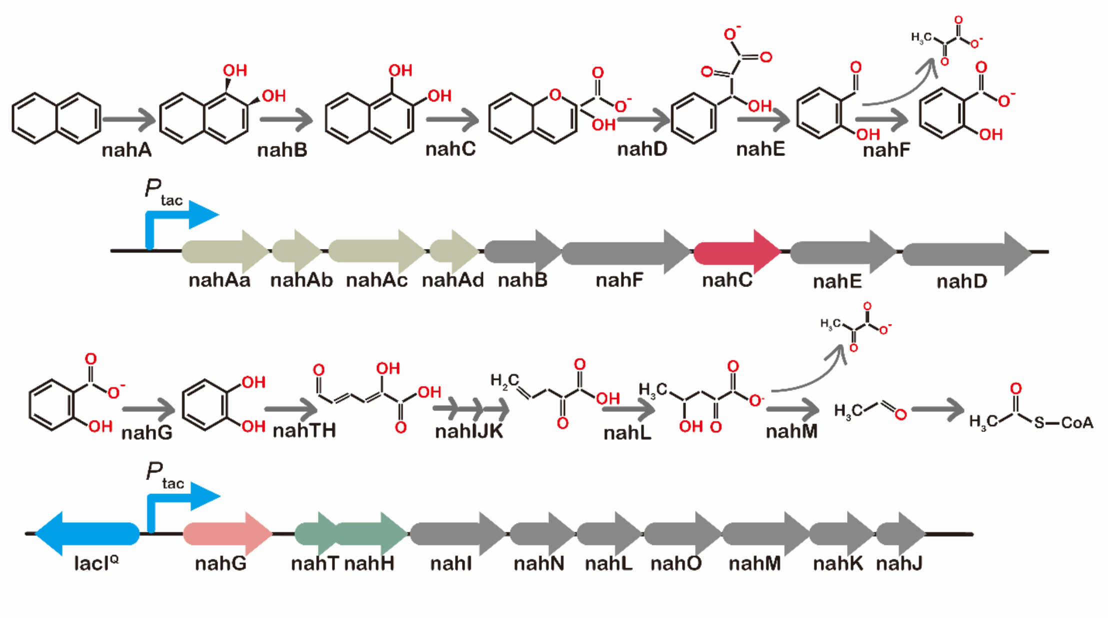

**Fig S4.1** The upstream and downstream gene clusters of naphthalene degradation in *P. stutzeri* AN1111 and their corresponding degradation pathways.

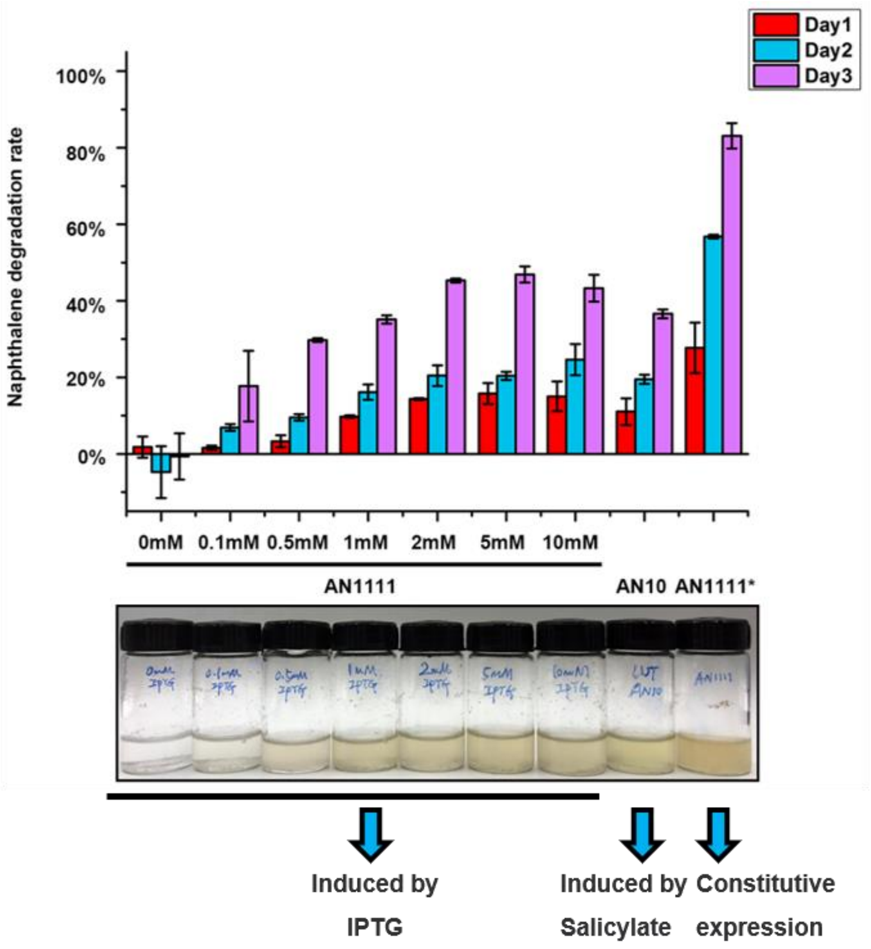

**Fig S4.2** Measurement of the naphthalene degradation rate and growth of the 'super strain' *P. stutzeri* AN1111 under different IPTG concentrations. Note that *P. stutzeri* AN10 is the wild type strain, and *P. stutzeri* AN1111$^{*}$ is a derived strain that lacks *lacI*$^{Q}$ gene, thus the expression of naphthalene degradation genes in this strain is constitutive.

*S4.1.2 Construction of the mutants that is only able to perform a subset of metabolic steps of naphthalene degradation*

The naphthalene degradation pathway in *P. stutzeri* AN1111 was further divided into four steps, which was associated with three intermediates, 1, 2- hydroxynaphthalene, salicylate, and catechol (Figure 5A). The four key enzymes responsible for the four metabolic steps were encoded by *nahA* gene (encodes a naphthalene dioxygenase), *nahC* gene (encodes a 1, 2-dihydroxynaphthalene dioxygenase), *nahG* gene (encodes a salicylate hydroxylase), and *nahH* gene (encodes a catechol 2, 3-dioxygenase).

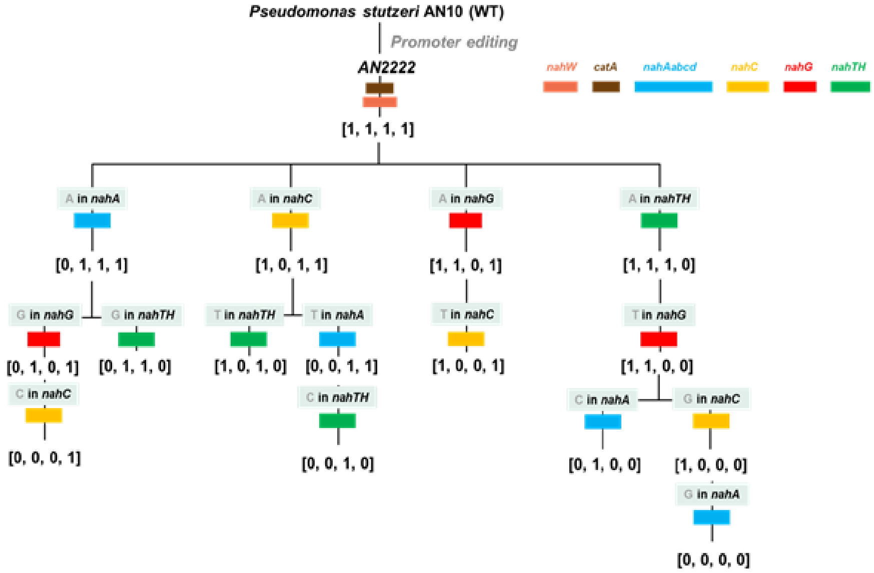

Fig S4.3 Schematic diagram showing the standard workflow to construct the strains in this study. Colorized rectangle indicates that different genes were knocked out in each step.

To construct the mutants that only possess ability to performed one or a subset of metabolic steps in naphthalene degradation, we knocked out the four key genes one by one following a standard workflow, as shown in Figure S4.3. As a result, we totally obtained 16 mutants that possess varied ability of naphthalene degradation. In the strain name, '1' denotes that the related key gene is retained in the strain, thus it can carry out the corresponding step. In contrast, '0' denotes that the related key gene is defective, thus it is not capable of carrying out the corresponding step. For example, strain *P. stutzeri* AN1000 can convert naphthalene to 1,2-dihydroxynaphthalene, but cannot carrying out the rest of metabolic steps. In this study, we only use eight strains to construct the synthetic communities, that is, *P. stutzeri* AN1000, *P. stutzeri* AN0100, *P. stutzeri* AN0010, *P. stutzeri* AN0001, *P. stutzeri* AN0011, *P. stutzeri* AN1100, *P. stutzeri* AN0111, and *P. stutzeri* AN1110.

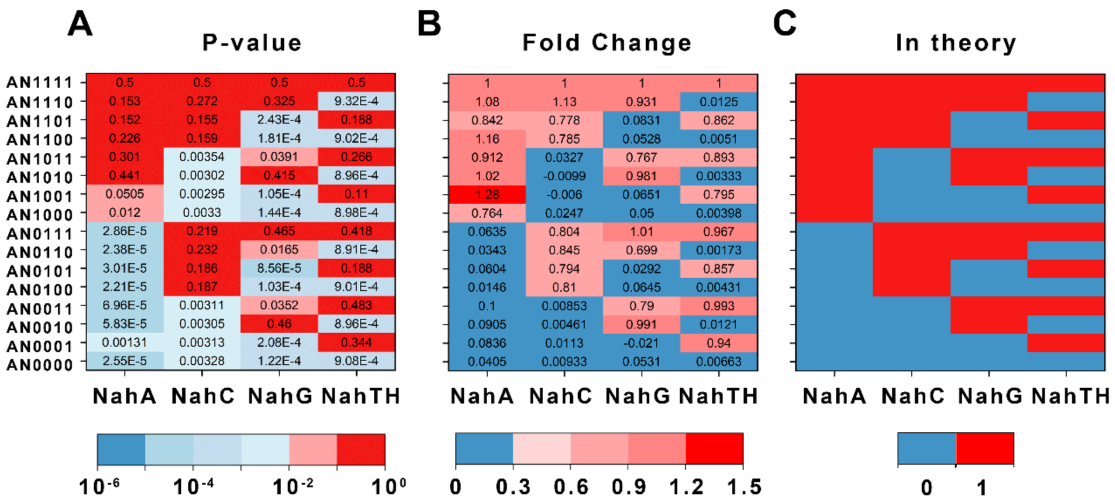

**Figure S4.4** Enzyme activity assays of all the strains used for constructing synthetic communities. (A) Two-tailed T-test showed the significant difference ($p$-value) of the enzyme activity of the four key enzymes between all other strains with *P. stutzeri* AN1111. (B) Average ratio of enzyme activity of four key enzymes in all strains to that of *P. stutzeri* AN1111. (C) Theoretical phenotypes of all strains. '0' means the strain should not possess the enzyme activity, while '1' means the strain should possess the enzyme activity.

#### *S4.1.3 Genetic manipulation and strain validation*

The genetic manipulations were implemented by allele exchange using the suicide plasmid pK18mobsacB [20,21]. The constructed strains were validated by PCR and DNA sequencing. In addition, enzymic activity assays were performed to verify the phenotypes of different strains, following the methods reported before (Reference[22] for naphthalene dioxygenase, Reference[23] for 1, 2-dihydroxynaphthalene dioxygenase, Reference[24] for salicylate 1-hydroxylase, and Reference[22] for catechol 2, 3-dioxygenase). Moreover, we also mono-cultured these strains in minimal medium supplemented with naphthalene, 1, 2- hydroxynaphthalene, salicylate, and catechol as sole carbon source. As shown in Figure S4.4 and Figure S4.5, phenotypes of all the strains is consistent with our design.

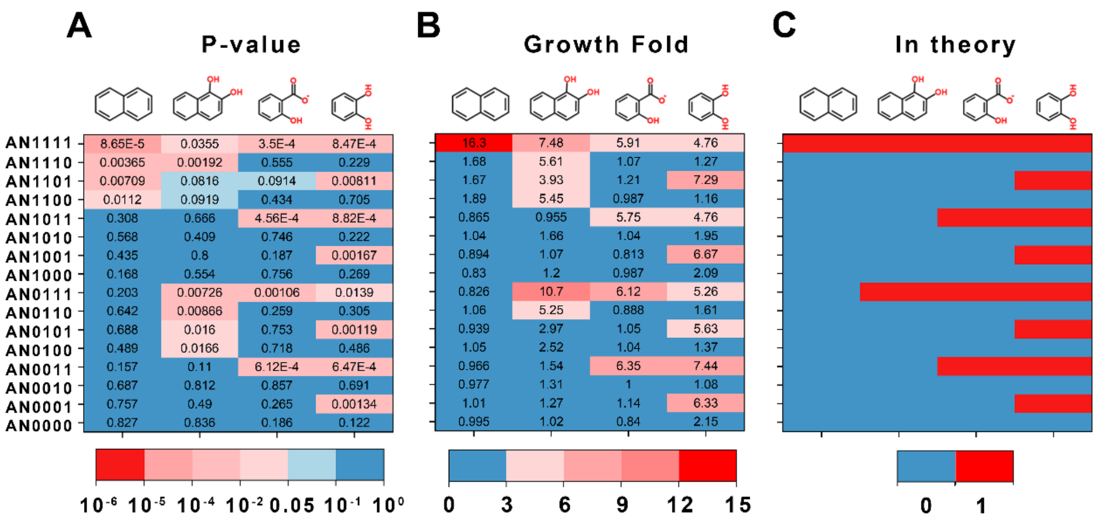

**Fig S4.5** The growth of all the strains using naphthalene and three intermediates as the sole carbon source. (A) Double-tailed T-test showed significant difference (*P*-values) between growth fold of all strains with that of *P. stutzeri* AN1111 in each carbon source. (B) relative growth fold between all strains with *P. stutzeri* AN1111 in each carbon source. (C) Theoretical phenotypes of all strains. '0' means the strain should not grow in the related carbon source, while '1' means the strain should grow in the related carbon source.

*S4.1.4 Modifying m value of the synthetic microbial communities by constructing two additional genetic modules*

For experimentally verifying our proposed rule, we need to experimentally modify the relative growth difference (*m*) between the different members of synthetic microbial communities. To achieve this goal, we designed two genetic modules, as shown in Figure 3A. In these modules, where the expression of toxic protein, CcdB[25], or X174[26], are controlled by a rhamnose-inducing promoter system, *rhaSR-PrhaBAD*. To construct these modules, fragment of *ccdB* or *x174* gene synthesized and cloned into a previous-designed miniTn7 plasmid, pJM220[27] and delivered into the strain by four-parental mating conjugation following the standard protocol[28]. The constructed strains were validated by PCR and DNA sequencing, in which the genetic modules were located 25-nucleotides downstream of the *glmS* gene in the chromosomes.

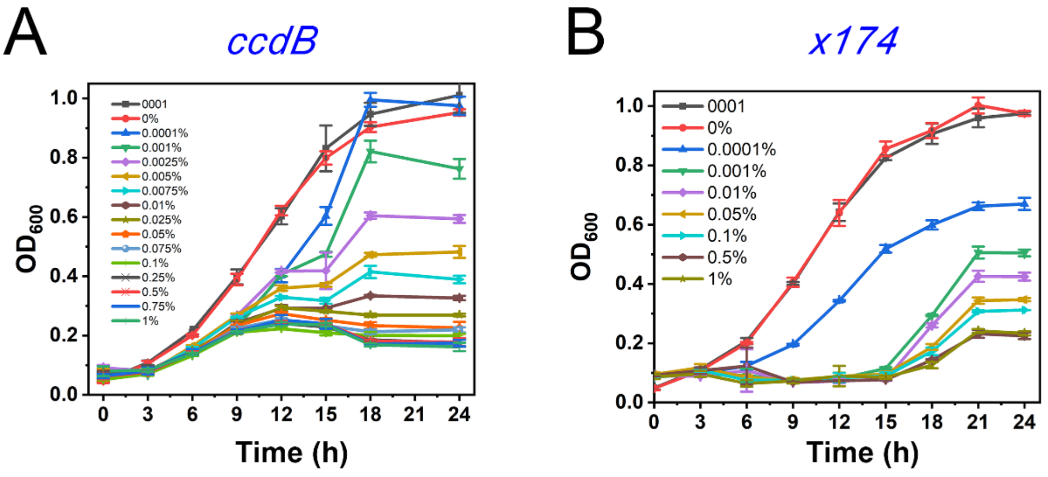

Fig S4.6 Effects of inducer concentration on the growth of the derivative strains of *P. stutzeri* AN0001 containing the death-rate modulating modules. Growth curves of *P. stutzeri* AN0001ccdB (A), and *P. stutzeri* AN0001x174 (B) at different IPTG concentrations are shown.

The expression of toxic proteins poisons the strain carrying the modules, so its death rate should increase with the concentration of the inducer, rhamnose. To quantify the relationship between rhamnose concentration and the strain death rate, we cultured the two strains, *P. stutzeri* AN0001ccdB and AN0001x174 who carries CcdB- and X174-modules, respectively, in minimum medium using one of the products, pyruvate, as the carbon source, and supplemented with different concentration of rhamnose. We measured the growth curve of the strains in these conditions (Figure S4.6), and fitted these data using the growth equations same as in our model:

$$\frac{dX}{dt}=gX\left(1-\frac{X}{N_m}\right)-DX \qquad [S4.1]$$

in which the specific growth rate ($g$) and the maximum biomass capacity ($N_m$) were set to be constant that is measured and fitted from the growth of the strain do not carrying the modules. Next, we fitted the death rates with the concentration of rhamnose, in which we found that the death rates possess a significant linear relationship with the logarithmic form of the rhamnose concentration (Table S7). Therefore, we can quantitively predict the death rates of the modified strains from the specific rhamnose concentration using the fitted formula in Table S7, so that calculate the $m$ value of the synthetic microbial communities in a given culture condition.

*S4.1.5 Construction of the synthetic microbial communities engaged in MDOL*

We constructed three synthetic microbial communities engaged in two-step MDOL, including

(1) Community composed of strain AN1000 and AN0111 (or its derived strain containing the death-rate modulating modules), in which AN1000 intracellularly converts naphthalene to 1, 2- hydroxynaphthalene, and the other strains further transform the intermediate to final end products, pyruvate and acetyl-CoA (Figure 3A).

(2) Community composed of strain AN1100 and AN0011 (or its derived strain containing the death-rate modulating modules), in which AN1100 intracellularly converts naphthalene to salicylate, and the other strain further transform the intermediate to final end products (Figure S9A).

(3) Community composed of strain AN1110 and AN0001 (or its derived strain containing the death-rate modulating modules), in which AN1110 intracellularly converts naphthalene to catechol, and the other strain further transform the intermediate to final end products (Figure S10A).

We also constructed a synthetic microbial community engaged in four-step MDOL, composed of strain AN1000, strain AN0100, strain AN0010, strain AN0001 (or its derived strain containing the death-rate modulating modules). In this community, strain AN1000 converts naphthalene to 1, 2- hydroxynaphthalene, which is then converted to salicylate by strain AN0100, then to catechol by strain AN0010, finally to the final end products by strain AN0001 (Figure 5A). Culture condition of the synthetic communities are described in Methods section.

*S4.1.6 Culturing of the synthetic microbial communities engaged in MDOL*

Our synthetic microbial communities were cultured in 25-mL flask containing 5 mL new fresh minimum medium[29]. To prepare the inoculum, *P. stutzeri* strains were first grown at 30˚C RB liquid medium (Yeast extract 10 g/L, beef extract 6 g/L, peptone 10 g/L, ammonium sulfate 5 g/L) by shaking at 220 rpm, supplemented with 50 μg/mL gentamicin. The cells were then washed by the minimum medium for twice to make an inoculum. For co-culture experiments, inocula of the two or four strains involved in a synthetic consortium were concentrated to an Optical Density (OD, measured at 600 nm) of 5.0, and mixed at equal initial abundance, or at a pre-designed initial ratio. The cultures were then inoculated to a 25-mL flask containing 5 mL new fresh minimum medium (starting OD: 0.05), supplemented with 2 mM IPTG, 50 μg/mL gentamicin, a pre-designed amount of rhamnose (ranges from 0 % to 0.1 % w/v), as well as naphthalene powder (1% w/v) as the sole carbon source. During the culture period, culture liquid (100 μL) was taken from system to measure OD to estimate the total biomass, as well as measure fluorescence intensity to estimate the growth of each population. The relative fraction was calculated by a method previously described[30]. Briefly, cultures of each populations were grown to mid-log phase at 30˚C (OD: ~0.3), diluted two-fold for eleven times, and the dilutions measured for their OD and fluorescence. Correlations between OD and fluorescence were then determined using the basic method defined in the LinerModelFit function of Mathematica software. Eventually, fluorescence values were transformed to the OD-estimated biomass to assess the growth of each population, and relative fraction was then calculated. These measurements, as well as the related measurements described below, were performed using a microplate reader (Molecular Devices, Sunnyvale, America). When applicable, the cultures were diluted by a factor of 20 after 72, 96 or 108 h, and transferred into a new fresh medium.

### S4.2 Mathematical modelling to predict the dynamics of the synthetic communities.

#### *S4.2.1 Modifying our basic model to match the property of our synthetic communities*

To mathematically predict the dynamics and assembly of our synthetic communities, we modify our mathematical model to match the property of our experiment settings. Firstly, the naphthalene degradation pathway possesses several specific mechanisms (The effect of each mechanism has been thoroughly discussed in Section S2), mainly including

(1) the substrate (naphthalene[31,32]) and the three intermediates (1, 2-hydroxynaphthalene[32], salicylate[32,33], or catechol[31,32,34]) are all bio-toxic;

(2) The intermediates 1,2-hydroxynaphthalene and catechol have been reported to be chemically unstable and may be converted to unavailable chemicals (1,2-naphthoquinone or benzoquinone) via a spontaneous way[35];

(3) Although the end products of the naphthalene degradation are the main carbon source that supports the growth of the community, a by-product, pyruvate, will be generated from the reaction that converts 1,2-hydroxynaphthalene to salicylate, which can be used as a direct carbon source[14] (Figure 3A).

Therefore, following the consideration of these additional mechanisms described in Section S2, we modified our basic model to match these settings. The dimensionless model regarding the synthetic communities engaged in two-step MDOL is given by

$$\frac{ds_{1,in}}{d\tau}=-a_1+\gamma_s\cdot\left(s_{out}-s_{1,in}\right) \qquad [S4.2]$$

$$\frac{ds_{2,in}}{d\tau}=\gamma_s\cdot\left(s_{out}-s_{1,in}\right) \quad [S4.3]$$

$$\frac{ds_{out}}{d\tau}=-x_1\cdot\gamma_s\cdot\left(s_{out}-s_{1,in}\right)-x_2\cdot\gamma_s\cdot\left(s_{out}-s_{1,in}\right) \quad [S4.4]$$

$$\frac{di_{1,in}}{d\tau}=a_1-\gamma_i\cdot\left(i_{1,in}-i_{out}\right)-l_i\cdot i_{1,in} \quad [S4.5]$$

$$\frac{di_{2,in}}{d\tau}=a_2 i_{2,in}+\gamma_i\cdot\left(i_{out}-i_{2,in}\right)-l_i\cdot i_{2,in} \quad [S4.6]$$

$$\frac{dp'_{1,in}}{d\tau}=a_1-bg_1 p'_{1,in}-\gamma_{p'}\cdot\left(p'_{1,in}-p'_{out}\right) \quad [S4.7]$$

$$\frac{dp'_{2,in}}{d\tau}=\gamma_{p'}\cdot\left(p'_{out}-p'_{2,in}\right)-bg_2 p'_{2,in} \quad [S4.8]$$

$$\frac{dp_{1,in}}{d\tau}=-Ig_1 p_{1,in}+\gamma_p\cdot\left(p_{out}-p_{1,in}\right) \quad [S4.9]$$

$$\frac{dp_{2,in}}{d\tau}=a_2 i_{2,in}-Ig_2 p_{2,in}-\gamma_p\cdot\left(p_{2,in}-p_{out}\right) \quad [S4.10]$$

$$\frac{di_{out}}{d\tau}=x_1\cdot\gamma_i\cdot(i_{1,in}-i_{out})-x_2\cdot\gamma_i\cdot(i_{out}-i_{2,in})-l_i\cdot i_{out} \quad [S4.11]$$

$$\frac{dp'_{out}}{d\tau}=x_1\cdot\gamma_{p'}\cdot\left(p'_{1,in}-p'_{out}\right)-x_2\cdot\gamma_{p'}\cdot\left(p_{out}-p_{1,in}\right) \quad [S4.12]$$

$$\frac{dp_{out}}{d\tau}=x_2\cdot\gamma_p\cdot\left(p_{2,in}-p_{out}\right)-x_1\cdot\gamma_p\cdot\left(p_{out}-p_{1,in}\right) \quad [S4.13]$$

$$\frac{dx_1}{d\tau}=\mu_1 t_1 x_1\left(1-\frac{x_1+x_2}{\rho}\right)-d_1 x_1 \quad [S4.14]$$

$$\frac{dx_2}{d\tau}=\mu_2 t_2 x_2\left(1-\frac{x_1+x_2}{\rho}\right)-d_2 x_2 \quad [S4.15]$$

$$\mu_1=(Ig_1 p_{1,in}+\delta\cdot bg_1 p'_{1,in})y_1 c_1 \quad [S4.16]$$

$$\mu_2=(Ig_2 p_{2,in}+\delta\cdot bg_2 p'_{2,in})y_2 c_2 \quad [S4.17]$$

$$t_1=(1-\theta_s s_{1,in})\cdot\frac{1}{1+\theta_i i_{1,in}} \quad [S4.18]$$

$$t_2=(1-\theta_s s_{2,in})\cdot\frac{1}{1+\theta_i i_{2,in}} \quad [S4.19]$$

The dimensionless model regarding the synthetic communities engaged in four-step MDOL is given by

$$\frac{ds_{1,in}}{d\tau}=-a_1+\gamma_s\cdot\left(s_{out}-s_{1,in}\right) \quad [S4.20]$$

$$\frac{ds_{k,in}}{d\tau}=\gamma_s\cdot\left(s_{out}-s_{k,in}\right) \qquad (k=2\sim 4) \quad [S4.21]$$

$$\frac{di^1_{1,in}}{d\tau}=a_1+\gamma_{i^1}\cdot\left(i^1_{out}-i^1_{1,in}\right)-l_{i^1}\cdot i_{1,in} \qquad [S4.22]$$

$$\frac{di^1_{2,in}}{d\tau}=-a_2\cdot i^1_{2,in}+\gamma_{i^1}\cdot\left(i^1_{out}-i^1_{2,in}\right)-l_{i^1}\cdot i_{2,in} \qquad [S4.23]$$

$$\frac{di^1_{3,in}}{d\tau}=\gamma_{i^1}\cdot\left(i^1_{out}-i^1_{3,in}\right)-l_{i^1}\cdot i^1_{3,in} \qquad [S4.24]$$

$$\frac{di^1_{4,in}}{d\tau}=\gamma_{i^1}\cdot\left(i^1_{out}-i^1_{4,in}\right)-l_{i^1}\cdot i^1_{4,in} \qquad [S4.25]$$

$$\frac{di^2_{1,in}}{d\tau}=\gamma_{i^2}\cdot\left(i^2_{out}-i^2_{1,in}\right) \qquad [S4.26]$$

$$\frac{di^2_{2,in}}{d\tau}=a_2\cdot i^2_{2,in}+\gamma_{i^2}\cdot\left(i^2_{out}-i^2_{2,in}\right) \qquad [S4.27]$$

$$\frac{di^2_{3,in}}{d\tau}=-a_3\cdot i^2_{3,in}+\gamma_{i^2}\cdot\left(i^2_{out}-i^2_{3,in}\right) \qquad [S4.28]$$

$$\frac{di^2_{4,in}}{d\tau}=\gamma_{i^2}\cdot\left(i^2_{out}-i^2_{4,in}\right) \qquad [S4.29]$$

$$\frac{di^3_{1,in}}{d\tau}=\gamma_{i^3}\cdot\left(i^3_{out}-i^3_{1,in}\right)-l_{i^3}\cdot i^3_{1,in} \qquad [S4.30]$$

$$\frac{di^3_{2,in}}{d\tau}=\gamma_{i^3}\cdot\left(i^3_{out}-i^3_{2,in}\right)-l_{i^3}\cdot i^3_{2,in} \qquad [S4.31]$$

$$\frac{di^3_{3,in}}{d\tau}=a_3\cdot i^3_{3,in}+\gamma_{i^3}\cdot\left(i^2_{out}-i^3_{3,in}\right) \qquad [S4.32]$$

$$\frac{di^3_{4,in}}{d\tau}=-a_4\cdot i^3_{4,in}+\gamma_{i^3}\cdot\left(i^2_{out}-i^3_{3,in}\right) \qquad [S4.33]$$

$$\frac{dp_{4,in}}{d\tau}=a_4\cdot i^3_{4,in}-Ig_4\cdot p_{4,in}+\gamma_p\cdot\left(p_{out}-p_{4,in}\right) \qquad [S4.34]$$

$$\frac{dp_{k,in}}{d\tau}=Ig_k\cdot p_{k,in}+\gamma_p\cdot\left(p_{out}-p_{k,in}\right) \qquad (k=1\sim3) \qquad [S4.35]$$

$$\frac{dp_{4,in}}{d\tau}=a_4\cdot i^3_{4,in}-Ig_4\cdot p_{4,in}+\gamma_p\cdot\left(p_{out}-p_{4,in}\right) \qquad [S4.36]$$

$$\frac{dp'_{1,in}}{d\tau}=a_1-bg_1\cdot p'_{1,in}-\gamma_{p'}\cdot\left(p'_{1,in}-p'_{out}\right) \qquad [S4.37]$$

$$\frac{dp'_{k,in}}{d\tau}=-bg_k\cdot p'_{k,in}+\gamma_p\cdot\left(p'_{out}-p'_{k,in}\right) \qquad (k=1\sim3) \qquad [S4.38]$$

$$\frac{ds_{out}}{d\tau}=-\sum_{k=1}^{4}x_k\cdot\gamma_{i^j}\cdot\left(s_{out}-s_{k,in}\right) \qquad [S4.39]$$

$$\frac{di^j_{out}}{d\tau}=-\sum_{k=1}^{4}x_k\cdot\gamma_i\cdot\left(i^j_{out}-i^j_{k,in}\right) \qquad (j=1\sim3) \qquad [S4.40]$$

$$\frac{dp_{out}}{d\tau}=-\sum_{k=1}^{4}x_k\cdot\gamma_p\cdot\left(p_{out}-p_{k,in}\right) \qquad [S4.41]$$

$$\frac{dp'_{out}}{d\tau}=-\sum_{k=1}^{N}x_k\cdot\gamma_p\cdot\left(p'_{out}-p'_{k,in}\right) \qquad [S4.42]$$

$$\frac{dx_k}{d\tau}=\mu_k x_k\left(1-\frac{\sum_{k=1}^{N}x_k}{\rho}\right)-d_k x_k \qquad [S4.43]$$

$$\mu_k=\left(Ig_k p_{k,in}+\delta\cdot bg_k p'_{k,in}\right)y_k c_k t_k \qquad [S4.44]$$

$$t_k=\left(1-\theta_s s_{1,in}\right)\cdot\prod_{j=1}^{3}\frac{1}{1+\theta_j i^j_{k,in}} \qquad [S4.45]$$

*S4.2.2 Parameterization of the model*

To parameterize the model for accurate prediction, we combined experimental measurements and data collection from previous studies. The values and sources of all the parameters used for experimental verification are listed in Table S8. Detailed protocols for parameterization are

(1) The reaction kinetics of each metabolic steps ($a_j$) are parameterized by considering the kinetics of the four key enzymes. These kinetic parameters are taken from previous reports. The values were converted and dimensionalized according to the method in Table S8. In a two-step MDOL community, several steps may be combined into one metabolic step in one strain. In these cases, we used the reaction rate of the slowest sub-step (that is, the rate-limiting step) for model calculation.

(2) The passive diffusion coefficients of naphthalene, 1, 2- hydroxynaphthalene, salicylate, catechol, and the end product pyruvate ($\gamma_s$, $\gamma_i$ and $\gamma_p$) were estimated using the predicted values of permeability of different aromatic hydrocarbons reported in Reference[36]. We assumed that the cell surface area is 5 μm$^2$ [37], and the extracellular

space of the culture system was 5 mL. Then the diffusion coefficients were calculated using the formula reported in Reference[36] and dimensionalized according to the method in Table S8.

(3) The metabolic burden of different strains ($c_k$) are determined by the competitive fitness assays between the measured strains using minimum medium supplemented with 2 mM IPTG, 50 μg/mL gentamicin, as well as pyruvate (0.5 % w/v) as the sole carbon source. In this condition, the strains can grow using the directly supplemented end product and still synthesize the enzymes for naphthalene degradation (since the expression is induced by IPTG). Therefore, the difference in relative fitness in this condition reflects the difference in metabolic burdens from expressing different sets of enzymes. The measurements of relative fitness followed the standard protocol reported in Reference[38] and reference[39].

(4) The maximum consumption rates of the final product ($Ig_k$) and the maximum biomass capacity ($\rho$) were estimated by measuring the growth curve of the strain *P. stutzeri* AN0001 using pyruvate as the sole carbon source, and fitted the data using the growth equations [S4.1]. In these measurements, we assumed that the dry biomass of each bacterial cell was $10^{-12}$ g, and the cell number per OD·mL strain is $1.6\times10^9$, thus the value estimated by OD can be transformed dry biomass, as we assumed in the model. Finally, the values were dimensionalized according to the method in Table S8 for model calculation.

(5) The initial biomass of each population ($x_{k,0}$) was set same as the experimental settings, transformed to dry biomass same as the above method, and then

dimensionalized according to the method in Table S8 for model calculation.

(6) As described in Section 2.3, In our culture system, naphthalene was suppled as powder and its extracellular concentration should constantly equal to its saturated concentration. Therefore, the initial naphthalene concentration ($S_0$) was determined by the previous reported saturation concentration of naphthalene under the circumstance of sufficient biological surfactant produced by *Pseudomonas*[40], set as $1.25\times10^{-3}$ M. The value is then dimensionalized according to the method in Table S8 for model calculation.

(7) As described above, a by-product, pyruvate, will be generated from the reaction converting 1,2-hydroxynaphthalene to salicylate. The maximum consumption rates of the by-product ($bg_k$) was assumed to be equal to $Ig_k$ (since the end-product and by-product are both pyruvate).

To measure the relative contribution of the by-product to end-product ($\delta$), we compared the growth fold of strain *P. stutzeri* AN1110 and strain *P. stutzeri* AN1111 supplemented with naphthalene, because in this condition, the former strain can only use the by-product as the direct sole carbon source, while the later can use both end-product and by-product. The resulting value of is $\delta$ approximately 0.1, which is then used for model simulation.

(8) The toxicity coefficient ($\theta_s$, $\theta_j$) of naphthalene, 1,2-hydroxynaphthalene, salicylate, and catechol, were determined using the similar protocol as before[1]. Briefly, strain *P. stutzeri* AN0000 (named as 'Cheater', who cannot perform both steps) were cultured in minimum medium supplemented with sufficient amount of pyruvate (0.34 mol/L, which means adding more pyruvate will no longer increase the bacterial growth rate)

and varied amount of the toxic substrate or intermediates. Growth curves were then estimated for each condition, and the data fitted to calculate the specific growth rate $g_s$ by a Boltzmann function using the Nonlinearmodelfit function of the Wolfram Mathematica. Subsequently, the similar equations described in Eqns. [S65]-[S73] (Section S2) was applied to characterize the toxic effects and fit the growth rate with the mass concentration, given by

$$T_R=\frac{1}{1+\frac{S_{i,in}}{K_{s,tox}}} \quad [S4.46]$$

$$T_L=1-\frac{S_{i,in}}{K_{s,tox}} \quad [S4.47]$$

$$T_E=e^{-\frac{S_{i,in}}{K_{s,tox}}} \quad [S4.48]$$

$$\frac{dX}{dt}=g_s TX\left(1-\frac{X}{\rho}\right)-DX \quad [S4.49]$$

The fit results suggest that the optimal fit of the toxicity coefficient of naphthalene holds a linear relationship with its concentration [Eqn. S4.47], and the best fit of its $K_{s,tox}$ is 0.0043 (Figure S11A). Furthermore, the optimal fits of the toxicity coefficient of the three intermediates holds a reciprocal relationship with their concentrations [Eqn. S4.46], and the best fit of its $K_{i,tox}$ are 0.00057, 0.00031 and 0.0028, respectively (Figure S11B-C). The values are then dimensionalized according to the method in Table S8, thus obtained values of $\theta_s$, $\theta_j$ for model calculation.

*S4.2.3 Predicting the dynamics of the synthetic communities*

To predict the dynamics of the synthetic community engaged in two-step MDOL, we performed numeric simulations using the ODE system shown in Eqns. [S4.2]-[S4.19] (Figure 3B-D; Figure S9B-D; Figure S10B-D). To predict the dynamics of the synthetic community engaged in four-step MDOL, in addition to the simulations using the ODE system shown in Eqns. [S4.20]-[S4.45] (Figure 5C; Green dots), we also directly used Eqn. [3] to predict the relative abundance of the strain AN0001, or its derived strain containing the death-rate modulating modules (Figure 5C; Red dots).

**S5 Supplementary Tables**

**Table S1** Definitions of variables and dimensionless methods for the model regarding a two-step MDOL community

| Variable | Description | Units | ND variable | ND Value Range |
|---|---|---|---|---|
| $S_{i,in}$ | Concentration of Intracellular *substrate* in the *ith* population ($i$=1, 2). | M | $s_{i,in}=S_{i,in}/K_1$ | $0 \sim 10^5$ |
| $S_{out}$ | Concentration of extracellular *substrate*. | M | $s_{out}=S_{out}/K_1$ | $0 \sim 10^5$ |
| $I_{i,in}$ | Concentration of Intracellular *intermediate* in the *ith* population ($i$=1, 2). | M | $i_{i,in}=I_{i,in}/K_1$ | $0 \sim 10^2$ |
| $I_{out}$ | Concentration of extracellular *intermediate*. | M | $i_{out}=I_{out}/K_1$ | $0 \sim 10^2$ |
| $P_{i,in}$ | Concentration of Intracellular *product* in the *ith* population ($i$=1, 2). | M | $p_{i,in}=P_{i,in}/K_1$ | $0 \sim 10^2$ |
| $P_{out}$ | Concentration of extracellular *product*. | M | $p_{out}=P_{out}/K_1$ | $0 \sim 10^2$ |
| $X_i$ | Total biomass of the *ith* population ($i$=1, 2). | $g\cdot L^{-1}$ | $x_i=X_i\cdot V_c$ | $0 \sim \rho$ |
| $t$ | Time | | $\tau=t\cdot k_1$ | |
| $P'_{i,in}$ | Concentration of Intracellular *byproduct* in the *ith* population ($i$=1, 2). | | $p'_{i,in}=P'_{i,in}/K_1$ | $0 \sim 10^2$ |

| | | | |
|---|---|---|---|
| $P'_{out}$ | Concentration of extracellular *byproduct*. | $p'_{out}=P'_{out}/K_1$ | $0 \sim 10^2$ |

**Table S2** Definitions, dimensionless methods and default values of the parameters involved in the model regarding a two-step MDOL community

| Parameter | Description | Value | Source | ND parameter | Default ND Value (Range) |
|---|---|---|---|---|---|
| **Parameters for the basic model** | | | | | |
| $K_j$ | Michaelis-Menten constant for the *jth* reaction ($j$ =1, 2). | $10^{-4}$ M | 41 | | |
| $E_j$ | Concentration of the *jth* enzyme ($j$ =1, 2). | $10^{-6}$ mol·g$^{-1}$ | 41-43 | $e_j=E_j/K_j$ | $10^1$ ($1\sim10^3$) |
| $k_j$ | Specific rate of the *jth* reaction ($j$ =1, 2). | 10 s$^{-1}$ | 36 | $\alpha_j=k_j/k_1$, $a_1=e_1$, $a_2=\alpha_2 e_2$ | |
| $r_s$, $r_i$, $r_p$ | Diffusivity of the S, I, and P. | 10 s$^{-1}$ | 44 | $\gamma_s=r_s/k_1$, $\gamma_i=r_i/k_1$, $\gamma_p=r_p/k_1$ | 1 ($10^{-2}\sim10^1$) |
| $V_c$ | The volume of cells per g cells | $10^{-3}$ L· g$^{-1}$ | | | |

| | | | | | |
|---|---|---|---|---|---|
| $Kg_1$, $Kg_2$ | Half-saturation constant of Monod growth. | $10^{-4}$ M | 45 | | |
| $kg_1$, $kg_2$ | Maximum consumption rate of P. | $10^{-5}$ mol·g$^{-1}$·s$^{-1}$ | 44 | $Ig_i=kg_i/(V_cKg_ik_i)$ | 10 ($10^{-1}$~$10^2$) |
| $g_i$ | Specific growth rate of *ith* population ($i$=1, 2). | s$^{-1}$ | | $\mu_i=g_i/k_1$ | |
| $D_i$ | Death rate of the *ith* population ($i$=1, 2). | $10^{-5}$ s$^{-1}$ | 45 | $d_i=D_i/k_1$ | $10^{-6}$ ($10^{-7}$~$10^{-5}$) |
| $Y$ | Yield coefficient for biomass production | $10^2$ g·mol$^{-1}$ | 45 | $y=Y\cdot K_1\cdot V_c$ | $10^{-5}$ ($10^{-6}$~$10^{-4}$) |
| $N_m$ | Carrying capacity of the whole populations | 10 g· L$^{-1}$ | 44 | $\rho=N_m\cdot V_c$ | $10^{-2}$ ($10^{-4}$ ~1) |
| $c_j$ | Metabolic burden for the *ith* population ($i$=1, 2). | | 46 | $c_j$ | 1 ($10^{-2}$~1) |
| $X_{i,0}$ | Initial biomass of the *ith* population, ($i$=1, 2). | 0.1 g· L$^{-1}$ | 45 | $x_{i,0}=X_{i,0}\cdot V_c$ | $10^{-4}$ ($10^{-6}$~$10^{-5}$) |

**Additional parameters (For the models generalizing the basic rule)**

| | | | | | |
|---|---|---|---|---|---|
| $r_{i1}$, $r_{i2}$, $r_{p1}$, $r_{p2}$ | Diffusivity of the S, I, and P in the cells of the *ith* population (*i*=1, 2). | 10 $s^{-1}$ | 36 | $\gamma_{i1}=r_{i1}/k_1$, $\gamma_{p1}=r_{p1}/k_1$, $\gamma_{i2}=r_{i2}/k_1$, $\gamma_{p2}=r_{p2}/k_1$ | |
| $r^a_{i1}$, $r^a_{i2}$, $r^a_{p1}$, $r^a_{p2}$ | Active transport rate of the S, I, and P in the cells of the *ith* population (*i*=1, 2). | 10 $s^{-1}$ | 36 | $\gamma^a_{i1}=r_{i1}/k_1$, $\gamma^a_{p1}=r_{p1}/k_1$, $\gamma^a_{i2}=r_{i2}/k_1$, $\gamma^a_{p2}=r_{p2}/k_1$ | 1 ($10^{-2}$~$10^1$) |
| $L_i$, $L_p$ | Abiotic degradation rate of I, and P | 0.05 $s^{-1}$ | 35 | $l_i=L_i/k_1$, $l_p=L_p/k_1$ | $5\times10^{-3}$ (0~$10^{-2}$) |
| $S_0$ | Initial concentration of *substrate* | 1 M | | $s_0=S_0/K_1$ | 0~$10^6$ |
| $E'_2$ | Concentration of the *second* enzyme | $10^{-6}$ mol·$g^{-1}$ | 41-43 | $e'_2=E'_2/(V_cK_1)$, $\alpha_2=k_2/k_1$, $a'_2=\alpha_2 e'_2$ , $\beta_2=K'_2/K_1$ | $10^1$ 1~$10^3$ |
| $k_2$ | Specific rate of the *second* reaction. | 10 $s^{-1}$ | 36 | | |

| | | | | | |
|---|---|---|---|---|---|
| $K_2'$ | Michaelis-Menten constant for *second* reaction | | 41 | | |
| $kg_1',\ kg_2'$ | Maximum consumption rate of product for microbial growth. | $10^{-5}$ mol·g$^{-1}$·s$^{-1}$ | 44 | $Ig_1'=kg_1'/(V_cK_1k_1),\ Ig_2'=kg_2'/(V_cK_1k_2),$ $v_1=Ig_1'/a_1,\ v_2=Ig_2'/a_2$ | 1 ($10^{-1}$~10) |
| $Kg_1,\ Kg_2$ | Half-saturation constant of Monod growth for the *ith* population (*i*=1, 2). | $10^{-4}$ M | 45 | $\beta_{g1}=Kg_1/K_1,\ \beta_{g2}=Kg_2/K_1,$ $b_1=\beta_{g1}/(a_1\gamma_p),\ b_2=\beta_{g2}/(a_1\gamma_p)$ | $10^{-1}$ ($10^{-2}$~1) |
| $Bg_1,\ Bg_2$ | Consumption rate of *byproduct* for microbial growth. | 0.01 s$^{-1}$ | 45 | $bg_1=Bg_1/(V_ck_1),\ bg_2=Bg_2/(V_ck_1)$ | 1 |
| $\delta$ | Relative contribution ratio to biomass of *byproduct* to *product*. | | | $\delta$ | $10^{-5}$~$10^{-1}$ |

**Table S3** Definitions, dimensionless methods and values/ranges of different toxic coefficients

| Variables (Description) | Formula* | ND variables | ND formula* | ND Value Range |
|---|---|---|---|---|
| $ToS_i$ (Toxic coefficient of Intracellular *substrate* in the *ith* population (*i*=1, 2).) | $\frac{1}{1+\frac{S_{i,in}}{K_{s,tox}}}$ <br> $1-\frac{S_{i,in}}{K_{s,tox}}$ <br> $e^{-\frac{S_{i,in}}{K_{s,tox}}}$ | $\theta_s=\frac{K_1}{K_{s,tox}}$ | $t_i=\frac{1}{1+\theta_s s_{i,in}}$ <br> $t_i=1-\theta_s s_{i,in}$ <br> $t_i=e^{-\theta_s s_{i,in}}$ | 0 ~ 1 |
| $ToI_i$ (Toxic coefficient of Intracellular *intermediate* in the *ith* population (*i*=1, 2). | $\frac{1}{1+\frac{I_{i,in}}{K_{I,tox}}}$ <br> $1-\frac{I_{i,in}}{K_{I,tox}}$ <br> $e^{-\frac{I_{i,in}}{K_{I,tox}}}$ | $\theta_i=\frac{K_1}{K_{I,tox}}$ | $t_i=\frac{1}{1+\theta_i i_{i,in}}$ <br> $t_i=1-\theta_i i_{i,in}$ <br> $t_i=e^{-\theta_i i_{i,in}}$ | 0 ~ 1 |
| $ToP_i$ Toxic coefficient of Intracellular *product* in the *ith* population (*i*=1, 2). | $\frac{1}{1+\frac{P_{i,in}}{K_{P,tox}}}$ <br> $1-\frac{P_{i,in}}{K_{P,tox}}$ <br> $e^{-\frac{P_{i,in}}{K_{P,tox}}}$ | $\theta_s=\frac{K_1}{K_{P,tox}}$ | $t_i=\frac{1}{1+\theta_p p_{i,in}}$ <br> $t_i=1-\theta_p p_{i,in}$ <br> $t_i=e^{-\theta_p p_{i,in}}$ | 0 ~ 1 |

*Note: The three expressions in the second column show the reciprocal, linear, and logarithmic forms of toxic coefficients, respectively.

**Table S4** Definitions of variables and dimensionless methods for the model regarding a multi-step MDOL community

| Variable | Description | Units | ND variable | ND Value Range |
|---|---|---|---|---|
| **Variables for the model regarding a three-step MDOL community** | | | | |
| $S_{i,in}$ | Concentration of Intracellular *substrate* in the *ith* population (*i*=1, 2, 3). | M | $s_{i,in}=\frac{S_{i,in}}{K_1}$ | $0 \sim 10^5$ |
| $S_{out}$ | Concentration of extracellular *substrate*. | M | $s_{out}=\frac{S_{out}}{K_1}$ | $0 \sim 10^5$ |
| $I1_{i,in}$ | Intracellular concentration of *first intermediate* in the *ith* population (*i*=1, 2, 3). | M | $i1_{i,in}=\frac{I1_{i,in}}{K_1}$ | $0 \sim 10^2$ |
| $I2_{i,in}$ | Intracellular concentration of *second intermediate* in the *ith* population (*i*=1, 2, 3). | M | $i2_{i,in}=\frac{I2_{i,in}}{K_1}$ | $0 \sim 10^2$ |
| $I_{out}$ | Concentration of extracellular *intermediate*. | M | $i_{out}=\frac{I_{out}}{K_1}$ | $0 \sim 10^2$ |
| $P_{i,in}$ | Concentration of Intracellular *product* in the *ith* population (*i*=1, 2, 3). | M | $p_{i,in}=\frac{P_{i,in}}{K_1}$ | $0 \sim 10^2$ |
| $P_{out}$ | Concentration of extracellular *product*. | M | $p_{out}=\frac{P_{out}}{K_1}$ | $0 \sim 10^2$ |

| | | | | |
|---|---|---|---|---|
| $X_i$ | Total biomass of the *ith* population (*i*=1, 2, 3). | $g\cdot L^{-1}$ | $x_i=X_i\cdot V_c$ | $0\sim\rho$ |
| $t$ | Time | | $\tau=t\cdot k_1$ | |
| **Variables for the model regarding a *N*-step MDOL community** | | | | |
| $I^j_{k,in}$ | Intracellular concentration of *jth intermediate* in the *kth* population (*j*=1, 2, 3…, *N*-1, *k*=1, 2, 3…, *N*). | M | $i^j_{k,in}=\frac{I^j_{k,in}}{K_1}$ | $0\sim 10^2$ |
| $I^j_{out}$ | Extracellular concentration of *jth intermediate*. | M | $i^j_{out}=\frac{I^j_{out}}{K_1}$ | $0\sim 10^2$ |
| $P_{k,in}$ | Concentration of Intracellular *product* in the *kth* population (*k*=1, 2, 3…, *N*). | M | $p_{i,in}=\frac{P_{i,in}}{K_1}$ | $0\sim 10^2$ |
| $P_{out}$ | Concentration of extracellular *product*. | M | $p_{out}=\frac{P_{out}}{K_1}$ | $0\sim 10^2$ |
| $X_k$ | Total biomass of the *kth* population (*k*=1, 2, 3…, *N*). | $g\cdot L^{-1}$ | $x_i=X_i\cdot V_c$ | $0\sim\rho$ |
| $t$ | Time | | $\tau=t\cdot k_1$ | |

**Table S5** Definitions, dimensionless methods and default values of the parameters involved in the model regarding a multi-step MDOL community

| Parameter | Description | Value | Source | ND parameter | Default Value (Range) |
|---|---|---|---|---|---|
| **Parameters for the model regarding a three-step MDOL community** | | | | | |
| $K_j$ | Michaelis-Menten constant for the *jth* reaction. | $10^{-4}$ M | 41 | | |
| $E_j$ | Concentration of the *jth* enzyme. | $10^{-6}$ mol·g$^{-1}$ | 41-43 | $e_j=\frac{E_j}{V_cK_j}$, $\alpha_j=\frac{k_j}{k_1}$, $a_1=e_1$, $a_2=\alpha_2e_2$, $a_3=\alpha_3e_3$ | $10^1$ ($1\sim10^3$) |
| $k_j$ | Specific rate of the *jth* reaction. | 10 s$^{-1}$ | 36 | | |
| $r_s$, $r_{i1}$, $r_{i2}$, $r_p$ | Diffusivity of the S, I1, I2, and P. | 10 s$^{-1}$ | 44 | $\gamma_s=r_s/k_1$, $\gamma_{i1}=r_{i1}/k_1$, $\gamma_{i1}=r_{i1}/k_1$, $\gamma_p=r_p/k_1$ | 1 ($10^{-2}\sim10^1$) |
| $V_c$ | The volume of cells per g cells | $10^{-3}$ L· g$^{-1}$ | | | |

| | | | | | |
|---|---|---|---|---|---|
| $Kg_1$, $Kg_2$, $Kg_3$ | Half-saturation constant of Monod growth. | $10^{-4}$ M | 45 | | |
| $kg_1$, $kg_2$, $kg_3$ | Maximum consumption rate of P. | $10^{-5}$ mol·g$^{-1}$·s$^{-1}$ | 44 | $Ig_1=\frac{kg_1}{V_c Kg_1 k_1}$ , $Ig_2=\frac{kg_2}{V_c Kg_2 k_1}$ , $Ig_3=\frac{kg_3}{V_c Kg_3 k_1}$ | 10<br>($10^{-1}$~$10^{2}$) |
| $g_i$ | Specific growth rate of the *ith* population. | s$^{-1}$ | | $\mu_i=g_i/k_1$ | |
| $D_i$ | Apparent maintenance rate of the *ith* population. | $10^{-5}$ s$^{-1}$ | 45 | $d_i=D_i/k_1$ | $10^{-6}$<br>($10^{-7}$~$10^{-5}$) |
| $Y_i$ | Yield coefficient for biomass production of the *ith* population. | $10^{2}$ g·mol$^{-1}$ | 45 | $y_i=Y_i \cdot K_1 \cdot V_c$ | $10^{-5}$<br>($10^{-6}$~$10^{-4}$) |
| $N_m$ | Carrying capacity of the whole populations | 10 g· L$^{-1}$ | 44 | $\rho=N_m \cdot V_c$ | $10^{-2}$<br>($10^{-4}$ ~1) |
| $c_j$ | Coefficient of metabolic burden for the *ith* population. | | 46 | | 1<br>($10^{-2}$~1) |
| $X_{i,0}$ | Initial biomass of the *ith* population. | 0.1 g· L$^{-1}$ | 45 | $x_{i,0}=X_{i,0} \cdot V_c$ | $10^{-4}$<br>($10^{-6}$~$10^{-5}$) |

| | | | | | |
|---|---|---|---|---|---|
| **Parameters for the model regarding a *N*-step MDOL community** | | | | | |
| $K_k$ | Michaelis-Menten constant for the *jth* reaction (*j*=1-*N*). | $10^{-4}$ M | 41 | $e_k=\frac{E_k}{V_cK_k}$, $\alpha_k=\frac{k_k}{k_1}$, $a_1=e_1$, $a_k=\alpha_k e_k$ | |
| $E_k$ | Concentration of the *jth* enzyme (*j*=1-*N*). | $10^{-6}$ mol·g$^{-1}$ | 41-43 | | $10^1$ ($1\sim10^3$) |
| $k_k$ | Specific rate of the *jth* reaction (*j*=1-*N*). | 10 s$^{-1}$ | 36 | | $5\times10^{-3}$ ($0\sim10^{-2}$) |
| $r_s$, $r_{ij}$, $r_p$ | Diffusivity of the S, $I_j$, and P. | 10 s$^{-1}$ | 44 | $\gamma_s=\frac{r_s}{k_1}$, $\gamma_{ij}=\frac{r_{ij}}{k_1}$, $\gamma_p=\frac{r_p}{k_1}$ | 1 ($10^{-2}\sim10^1$) |
| $V_c$ | The volume of cells per g cells | $10^{-3}$ L· g$^{-1}$ | | | |
| $Kg_k$ | Half-saturation constant of Monod growth for the *kth* population (*k*=1-*N*). | $10^{-4}$ M | 45 | | |
| $kg_k$ | Maximum consumption rate of *product* for the growth of the *kth* population (*k*=1-*N*). | $10^{-5}$ mol·g$^{-1}$·s$^{-1}$ | 44 | $Ig_k=kg_k/(V_cKg_kk_1)$ | 10 ($10^{-1}\sim10^2$) |

| | | | | | |
|---|---|---|---|---|---|
| $g_k$ | Specific growth rate of the *kth* population (*k*=1-*N*). | $s^{-1}$ | | $\mu_k=\frac{g_k}{k_1}$ | |
| $D_k$ | Apparent maintenance rate of the *kth* population (*k*=1-*N*). | $10^{-5}$ $s^{-1}$ | 45 | $d_k=\frac{D_k}{k_1}$ | $10^{-6}$ ($10^{-7}$~$10^{-5}$) |
| $Y_k$ | Yield coefficient for biomass production of the *kth* population (*k*=1-*N*). | $10^2$ g·mol$^{-1}$ | 45 | $y_k=Y_k{\cdot}K_1{\cdot}V_c$ | $10^{-5}$ ($10^{-6}$~$10^{-4}$) |
| $N_m$ | Carrying capacity of the whole populations | 10 g· L$^{-1}$ | 44 | $\rho=N_m{\cdot}V_c$ | $10^{-2}$ ($10^{-4}$ ~1) |

**Table S6 Strains and plasmids used in this study**

| Number | Strains or plasmids | Characterizations | Source |
|---|---|---|---|
| ***E. coli* strains** | | | |
| 1 | *E.coli* Trans5α | F$^-$ φ80 *lac ZΔM15 Δ(lacZYA-arg F) U169 endA1 recA1 hsdR17(*$r_k^-$, $m_k^+$*)* *supE44λ*$^-$ *thi -1 gyrA96 relA1 phoA* | TransGen |
| 2 | *E.coli* HB101 | F$^-$ *mcrB mrr hsdS20*(rB mB) *recA13 leuB6 ara-14 proA2 lacY1 galK2 xyl-5 mtl-1 rpsL20*(Smr) *glnV44* | Lab of Professor Lin Min, CAAS |
| 3 | *E.coli* Top10F’ | F′ [*lac*I$^q$, *Tn*10(Tet$^R$)] *mcr*A Δ(*mrr-hsd*RMS-*mcr*BC) φ80*lac*ZΔM15 Δ*lac*X74 *rec*A1 *ara*D139 Δ(*ara-eu*)7697 *gal*U *gal*K *rps*L(Str$^R$) *end*A1 *nup*G | Beijing Zoman Biotechnology Co., Ltd |
| ***P. stutzeri* strains** | | | |
| 1 | *P. stutzeri* AN0001 | AN1111△*nahC::G*△*nahG::T*△*nahTH::A*△*nahA::G* | This study |
| 2 | *P. stutzeri* AN0001 | AN1111△*nahA::A*△*nahC::C*△*nahG::G* | This study |
| 3 | *P. stutzeri* AN0010 | AN1111△*nahA::T*△*nahC::A*△*nahTH::C* | This study |
| 4 | *P. stutzeri* AN0011 | AN1111△*nahA::T*△*nahC::A* | This study |
| 5 | *P. stutzeri* AN0100 | AN1111△*nahA::C*△*nahG::T*△*nahTH::A* | This study |
| 6 | *P. stutzeri* AN0101 | AN1111△*nahA::A*△*nahG::G* | This study |
| 7 | *P. stutzeri* AN0110 | AN1111△*nahA::A*△*nahTH::G* | This study |
| 8 | *P. stutzeri* AN0111 | AN1111△*nahA::A* | This study |
| 9 | *P. stutzeri* AN1000 | AN1111△*nahC::G*△*nahG::T*△*nahTH::A* | This study |

| | | | |
|---|---|---|---|
| 10 | *P. stutzeri* AN1001 | AN1111△*nahC::T*△*nahG::A* | This study |
| 11 | *P. stutzeri* AN1010 | AN1111△*nahC::A*△*nahTH::C* | This study |
| 12 | *P. stutzeri* AN1011 | AN1111△*nahC::A* | This study |
| 13 | *P. stutzeri* AN1100 | AN1111△*nahG::T*△*nahTH::A* | This study |
| 14 | *P. stutzeri* AN1101 | AN1111△*nahG::A* | This study |
| 15 | *P. stutzeri* AN1110 | AN1111△*nahTH::A* | This study |
| 16 | *P. stutzeri* AN1111 | *P. stutzeri* AN10 △*nahW*△*catA*△*Pnah*1::*Ptac*△*Pnah*2::*Ptac*△*nahR:*$lacI^Q$ | This study |
| 17 | *P. stutzeri* AN10 | Wild type strain | [12] |
| 18 | *P. stutzeri* AN0111ccdB | *P. stutzeri* AN0111 containning pJMccdB | This study |
| 19 | *P. stutzeri* AN0111x174 | *P. stutzeri* AN0111 containning pJMx174 | This study |
| 20 | *P. stutzeri* AN0011ccdB | *P. stutzeri* AN0011 containning pJMccdB | This study |
| 21 | *P. stutzeri* AN0011x174 | *P. stutzeri* AN0011 containning pJMx174 | This study |
| 22 | *P. stutzeri* AN0001ccdB | *P. stutzeri* AN0001 containning pJMccdB | This study |
| 23 | *P. stutzeri* AN0001x174 | *P. stutzeri* AN0001 containning pJMx174 | This study |
| 24 | *P. stutzeri* AN0001ccdAB | *P. stutzeri* AN0001 containning pJMccdAB | This study |
| 25 | *P. stutzeri* AN1000mBeRFP | *P. stutzeri* AN1000 containning pmmPc-mBeRFP | This study |
| 26 | *P. stutzeri* AN1100mBeRFP | *P. stutzeri* AN1100 containning pmmPc-mBeRFP | This study |
| 27 | *P. stutzeri* AN1110mBeRFP | *P. stutzeri* AN1110 containning pmmPc-mBeRFP | This study |
| 29 | *P. stutzeri* AN0001egfp | *P. stutzeri* AN0001 containning pmmPc-egfp | This study |

| | | | |
|---|---|---|---|
| 30 | *P. stutzeri* AN0111ccdBegfp | *P. stutzeri* AN0111ccdB containning pmmPc-egfp | This study |
| 31 | *P. stutzeri* AN0011x174egfp | *P. stutzeri* AN0111x174 containning pmmPc-egfp | This study |
| 32 | *P. stutzeri* AN0011ccdBegfp | *P. stutzeri* AN0011ccdB containning pmmPc-egfp | This study |
| 33 | *P. stutzeri* AN0011x174egfp | *P. stutzeri* AN0011x174 containning pmmPc-egfp | This study |
| 34 | *P. stutzeri* AN0001ccdBegfp | *P. stutzeri* AN0001ccdB containning pmmPc-egfp | This study |
| 35 | *P. stutzeri* AN0001x174egfp | *P. stutzeri* AN0001x174 containning pmmPc-egfp | This study |
| 36 | *P. stutzeri* AN0001ccdABegfp | *P. stutzeri* AN0001ccdAB containning pmmPc-egfp | This study |
| 37 | *P. stutzeri* AN1000ecfp | *P. stutzeri* AN1000 containning pmmPc-ecfp | This study |
| 38 | *P. stutzeri* AN0100-dsred | *P. stutzeri* AN0100 containning pmmPc-dsred | This study |
| 39 | *P. stutzeri* AN0010-mBeRFP | *P. stutzeri* AN0100 containning pmmPc-mBeRFP | This study |
| **Plasmids** | | | |
| 1 | pRK2013 | Helper plasmid for conjugation; $Km^R$ | Lab of Professor Lin Min, CAAS |
| 2 | pJM220 | pUC18T-miniTn7T-gm-rhaSR-PrhaBAD; $Gm^R$ | Addgene[27] |
| 3 | pJMccdB | *ccdB* gene on pJM220 | This study |
| 4 | pJMx174 | *x174* gene on pJM220 | This study |
| 6 | pTNS3 | Helper plasmid; $Amp^R$ | Addgene[28] |

| | | | |
|---|---|---|---|
| 7 | pFLP3 | Containing the recombinase structural gene *flp* and *SacB* selection marker; $Tet^R$ | Addgene[28] |
| 8 | pMMPc-Gm | Shuttle vector between *E. coli* and *P. stutzeri* | Lab of Professor Ping Xu, SJTU |
| 9 | pmmPc-ecfp | *ecfp* gene on pmmPc-Gm | This study |
| 10 | pmmPc-dsred | *dsred* gene on pmmPc-Gm | This study |
| 11 | pmmPc-mBeRFP | *mBeRFP* gene on pmmPc-Gm | This study |
| 12 | pMMPc-egfp | *egfp* gene on pMMPc-Gm | This study |

Table S7 Relationship between death rate and rhamnose concentration

| **System** | **Fitted formula** | $R^2$ |
| --- | --- | --- |
| ccdB | $\boldsymbol{D_{ccdB}}$=0.081 lg $\boldsymbol{R_c}$ + 0.3406 | 0.9501 |
| X174 | $\boldsymbol{D_{x174}}$=0.0338 lg $\boldsymbol{R_c}$ + 0.2555 | 0.9818 |

**Table S8 The dimensionless methods and values of the parameters used in the model for predicting the experimental results**

| Parameter | Input Value | Source | ND parameter | ND Value |
|---|---|---|---|---|
| $K_1$ | $9.2\times10^{-7}$ M | 47 | | |
| $k_1$ | $1.82$ s$^{-1}$ | 47 | $e_1=\frac{E_1}{V_cK_1}$, $\alpha_1=\frac{k_1}{k_1}$, $a_1=e_1$ | 2174 |
| $E_1$ | $2\times10^{-6}$ mol·g$^{-1}$ | 43 | | |
| $K_2$ | $3.4\times10^{-5}$ M | 48 | | |
| $k_2$ | $22.7$ s$^{-1}$ | 48 | $e_2=\frac{E_2}{V_cK_2}$, $\alpha_2=\frac{k_2}{k_1}$, $a_2=\alpha_2e_2$ | 733.6 |
| $E_2$ | $2\times10^{-6}$ mol·g$^{-1}$ | 43 | | |
| $K_3$ | $9.6\times10^{-6}$ M | 48 | | |
| $k_3$ | $4.6$ s$^{-1}$ | 48 | $e_3=\frac{E_3}{V_cK_3}$, $\alpha_3=\frac{k_3}{k_1}$, $a_3=\alpha_3e_3$ | 526.6 |
| $E_3$ | $2\times10^{-6}$ mol·g$^{-1}$ | 43 | | |
| $K_4$ | $1.5\times10^{-6}$ M | 49 | $e_4=\frac{E_4}{V_cK_4}$, $\alpha_3=\frac{k_4}{k_1}$, $a_3=\alpha_4e_4$ | 146500 |
| $k_4$ | 200 | 49 | | |
| $E_4$ | $2\times10^{-6}$ mol·g$^{-1}$ | 43 | | |

| | | | | |
|---|---|---|---|---|
| $r_s$ | $3.35\ s^{-1}$ | 36 | $\gamma_s=\frac{r_s}{k_1}$ | 1.84 |
| $r_i$ | $3020\ s^{-1}$, $26.3\ s^{-1}$, $4.27\ s^{-1}$ | 36 | $\gamma_i=\frac{r_i}{k_1}$ | 1659, 14.45, 2.35 |
| $r_p$ | $25.1\ s^{-1}$ | 36 | $\gamma_p=\frac{r_p}{k_1}$ | 13.79 |
| $r_p{}'$ | $25.1\ s^{-1}$ | 36 | $\gamma_p{}'=\frac{r_p{}'}{k_1}$ | 13.79 |
| $V_c$ | $10^{-3}\ L\cdot g^{-1}$ | 37 | | |
| $K_g$ | $2\times10^{-5}$ M | 50 | | |
| $kg_1,\ kg_2$ | $6.37\times10^{-7}\ mol\cdot g^{-1}\cdot s^{-1}$ | Experimental measure | $Ig_i=kg_i/(V_cK_gk_i)$ | 35.0 |
| $D_k$ ($k$=1~3) | $4.92\times10^{-6}\ s^{-1}$ | Experimental measure | $d_k=D_k/k_k$ | $2.7\times10^{-6}$ |
| $D_4$ | Table S7 | Experimental measure | $d_4=D_4/k_4$ | |
| $Y$ | $100\ g\cdot mol^{-1}$ | 50 | $y=Y\cdot K_1\cdot V_c$ | $7.36\times10^{-8}$ |

| | | | | |
|---|---|---|---|---|
| $N_m$ | 1.6 g· $L^{-1}$ | Experimental measure | $\rho=N_m\cdot V_c$ | $1.6\times10^{-3}$ |
| $c_k$ ($k$=1~3) | 1 | Experimental measure | $c_k$ | 1 |
| $X_{i,0}$ | 0.04 g· $L^{-1}$ | Experimental measure | $x_{i,0}=X_{i,0}\cdot V_c$ | $4\times10^{-5}$ |
| $L_i$ | $10^{-4}$ $s^{-1}$ | [51] | $l_i=\frac{L_i}{k_1}$ | $5.49\times10^{-5}$ |
| $Bg_1$, $Bg_2$ | $1.76\times10^{-6}$ mol·$g^{-1}$·$s^{-1}$ | Experimental measure | $bg_i=Bg_i/(V_cK_gk_1)$ | 96.7 |
| $\delta$ | 0.1 | Experimental measure | $\delta$ | $10^{-5}$~$10^{-1}$ |
| $S_0$ | $1.25\times10^{-3}$ M | [40] | $s_0=S_0/K_1$ | 1359 |
| $K_{s,tox}$ | 0.0043 | Experimental measure | $\theta_s=K_1/K_{s,tox}$ | 0.00021 |
| $K_{i,tox}$ | 0.00057, 0.00031, 0.0028 | Experimental measure | $\theta_i=K_1/K_{i,tox}$ | 0.0016, 0.0032, 0.00033 |

### S6 Supplementary Figures

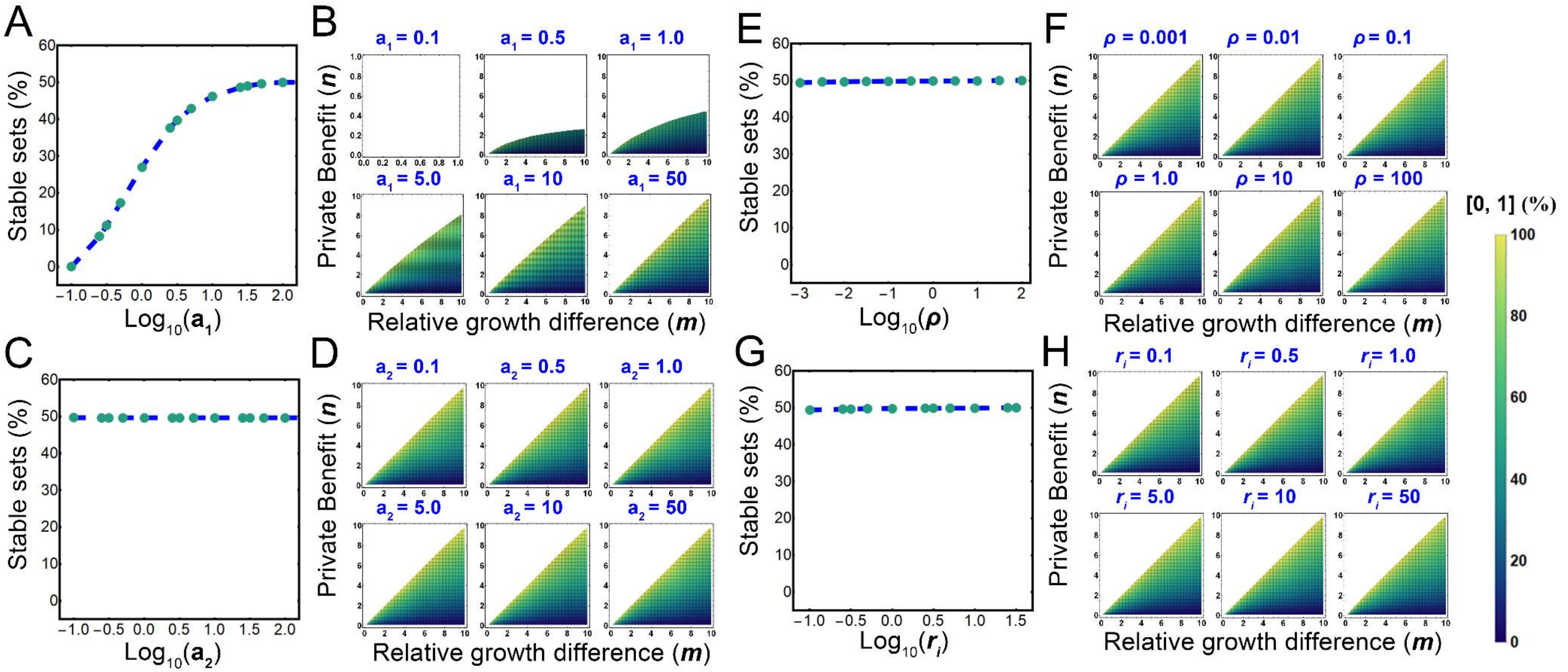

**Figure S1** Sensitive analyses of the basic assembly rule of a two-step MDOL community. We tested the effects of the rates of the first ($a_1$; A-B) and the second reaction ($a_2$; C-D), maximum biomass capacity ($\rho$; E-F), as well as the diffusive coefficient of I ($\gamma_i$; G-H) on our basic assembly rule. Plots A, C, E, G shows the percentage of the parameter sets that can leads to the co-exsitstence of [1, 0] and [0, 1] accounting for the total parameter set (10100) under different settings. The density maps in Plots B, D, F, H shows the range of stable parameter sets characterized by privatization benefit ($n$) and relative growth difference ($m$), as well as relative abundance of [0, 1] under different settings. Values of other parameters are given in Table S2.

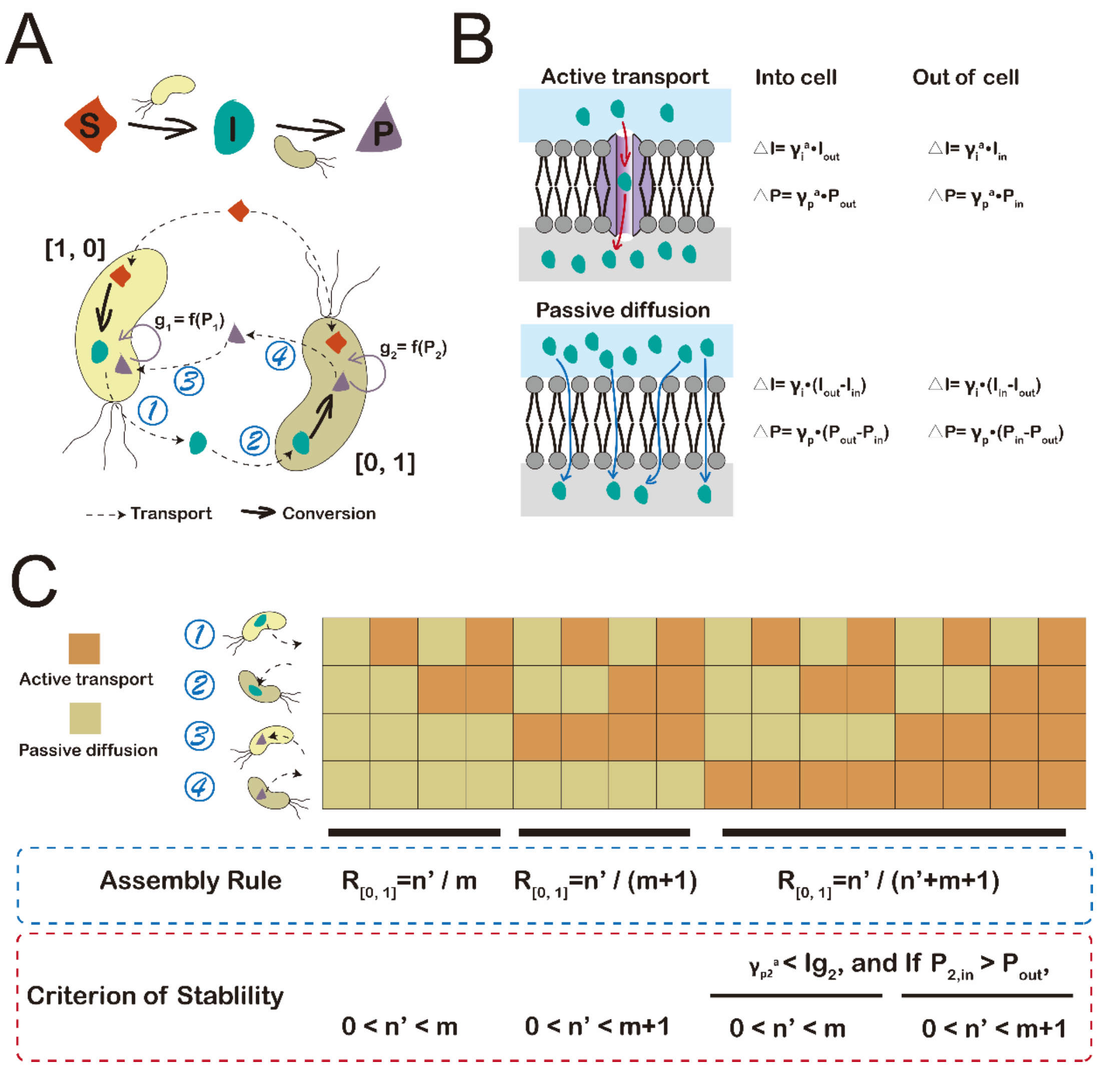

**Figure S2** Assembly rules when the transport of the intermediate (I) and product (P) is mediated by different mechanisms. (A) Schematic diagram shows the assumptions of our basic model considering differetnt transport mechanisms of I and P. Symbols ① to ④ indicate four transport processes of I and P. (B) Schematic diagram shows our assumptions on modelling the transporting processes. (C) Assembly rules of a two-step MDOL community under 16 different configurations of I and P transport.

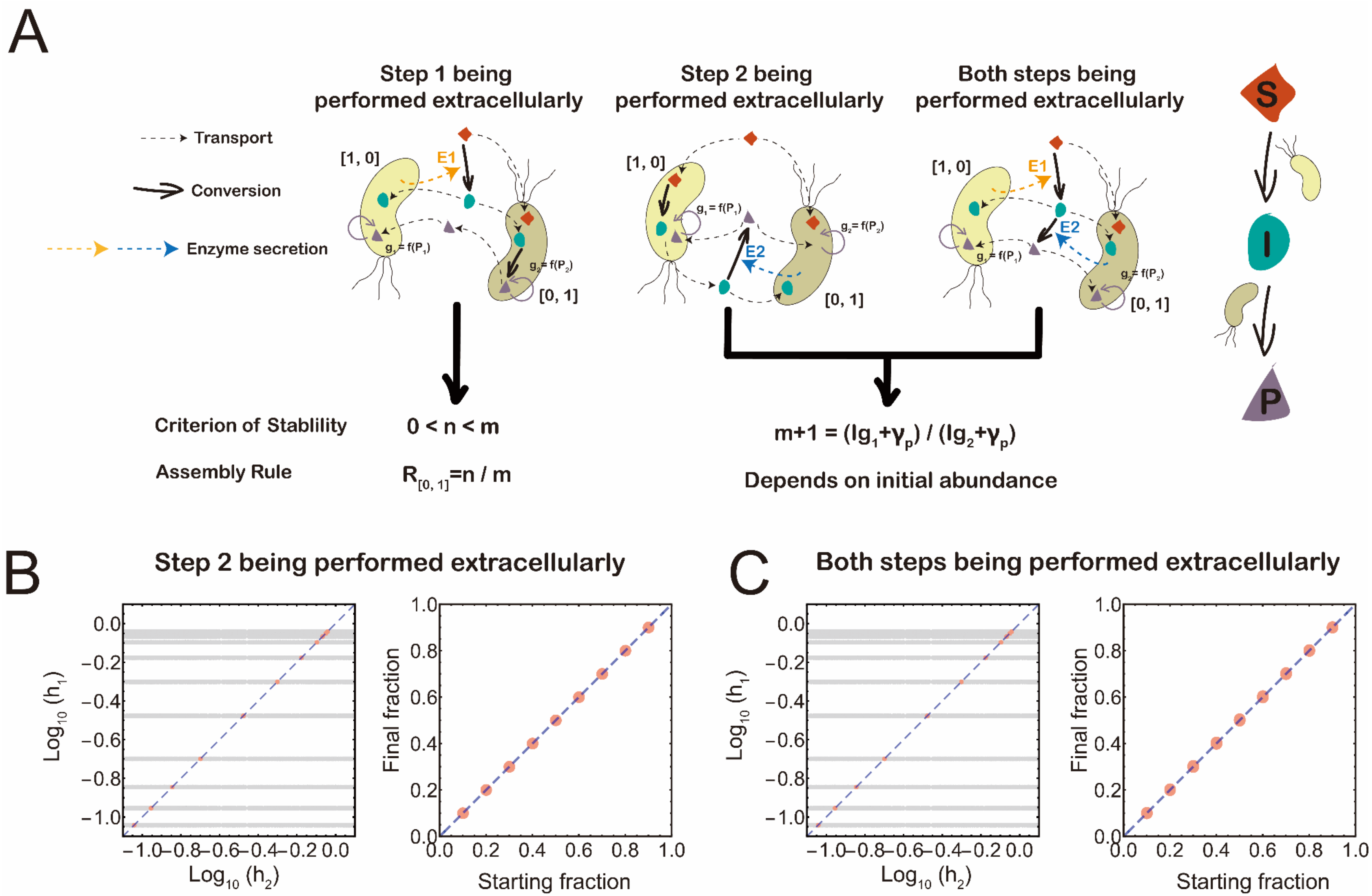
A
Step 1 being performed extracellularly
Step 2 being performed extracellularly
Both steps being performed extracellularly
Transport
Conversion
Enzyme secretion
[1, 0]
[0, 1]
E1
E2
S
I
P
Criterion of Stablility
0 < n < m
Assembly Rule
R[0, 1]=n / m
m+1 = (lg1+γp) / (lg2+γp)
Depends on initial abundance
B
Step 2 being performed extracellularly
C
Both steps being performed extracellularly
Log10 (h1)
Log10 (h2)
Final fraction
Starting fraction

**Figure S3** Assembly rules when alternative, or both metabolic reactions are catalyzed extracellularly. (A) Schematic diagram shows the assumptions of the three considered scenarios: the first reaction was performed extracellularly while the second was performed intracellularly (left); the second reaction was performed extracellularly while the first was performed intracellularly (middle); both reactions were performed extracellularly (right). Summary of the assembly rules in these scenarios is shown at the bottom of the corresponding diagram. (B-C) Derivations of the assembly rules when the second reaction are catalyzed extracellularly. The left graph depicts the distribution of 14461 designed parameter sets to test the condition required for the stability for a two-step MDOL community when the second reaction are catalyzed extracellularly (Eqn. [S2.35]. The grey dots indicate the distribution of all the sets; the orange dots indicate the parameter sets that lead to stable community dynamics. In addition, in the graph $h_1=\frac{c_1 Ig_1 y_1}{(Ig_1+\gamma_p)d_1}$ and $h_2=\frac{c_2 Ig_2 y_2}{(Ig_2+\gamma_p)d_2}$, so the blue line indicate the curve where the parameter sets satisfying Eqn. [S2.35] should locate in. The data derived from the simulations assuming the first reaction was performed intracellularly are shown in (B), while the data derived from the simulations assuming the first reaction was also performed extracellularly are shown in (C).

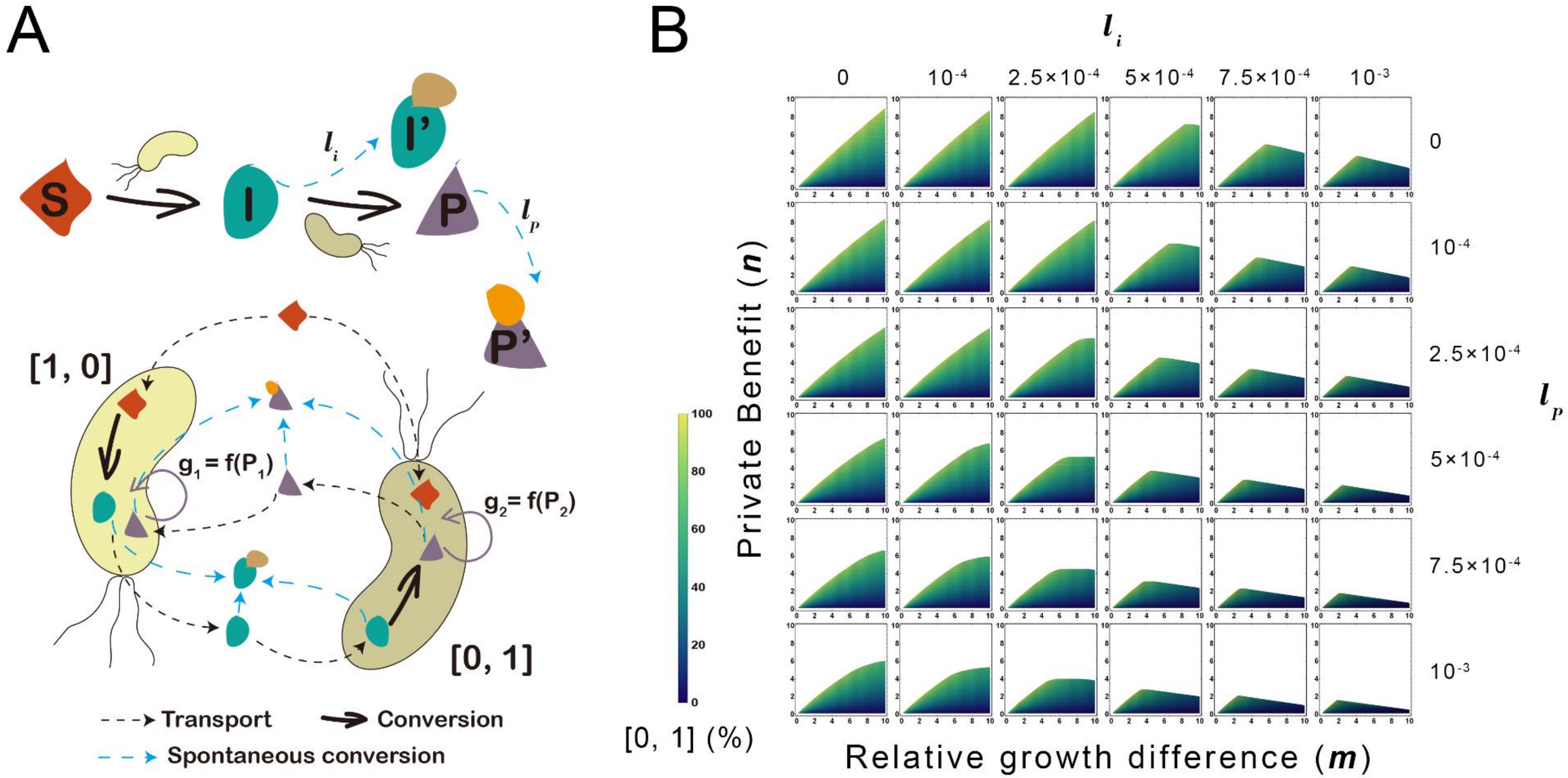

**Figure S4** Assembly rules when spontaneous conversion of the intermediate (I) and product (P) are involved. (A) Schematic diagram shows the assumptions of our basic model considering spontaneous conversion of I and P. We hypothesized that I can be converted to a derivant I', while P can be converted to a derivant P’. Both I’ and P’ cannot be undergone biotransformation subsequently. (B) The density maps show the parameter spaces characterized by private benefit $n$ and relative growth difference $m$, as well as relative abundance of [0, 1] in the stable communities, under different abiotic degradation rates of intermediate ($l_i$) and product ($l_p$). These plots are generated by mathematical simulations. Values of other parameters are given in Table S2.

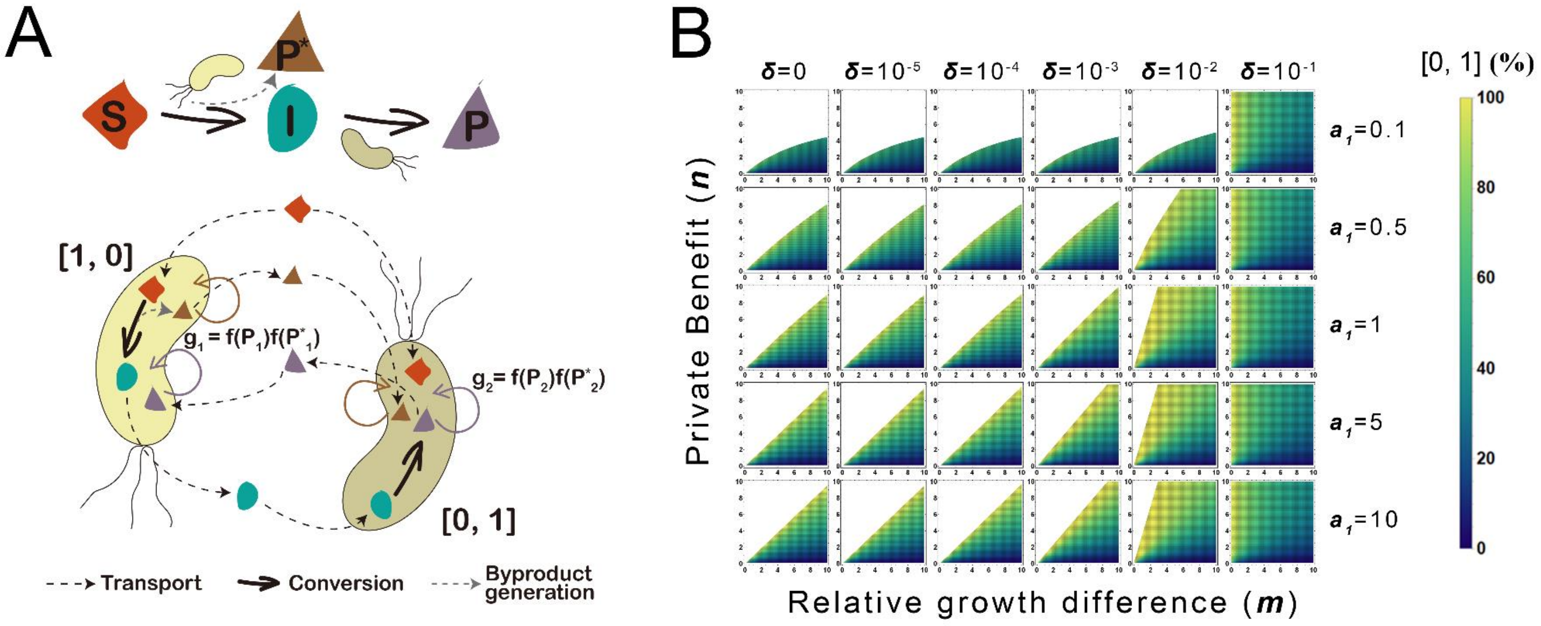

**Figure S5** Assembly rules when a byproduct (P*) is generated from the first reaction. (A) Schematic diagram shows the assumptions of our basic model considering the generation of P*. P* can be used for supporting the growth of the both populations, thus affecting their growth rate ($g_1$ and $g_2$). (B) The density maps show the parameter spaces characterized by private benefit $n$ and relative growth difference $m$, as well as relative abundance of [0, 1] in the stable communities, under different relative contributions of by-products ($\delta$) and different rate of first reaction ($a_1$). These plots are generated by mathematical simulations. Values of other parameters are given in Table S2.

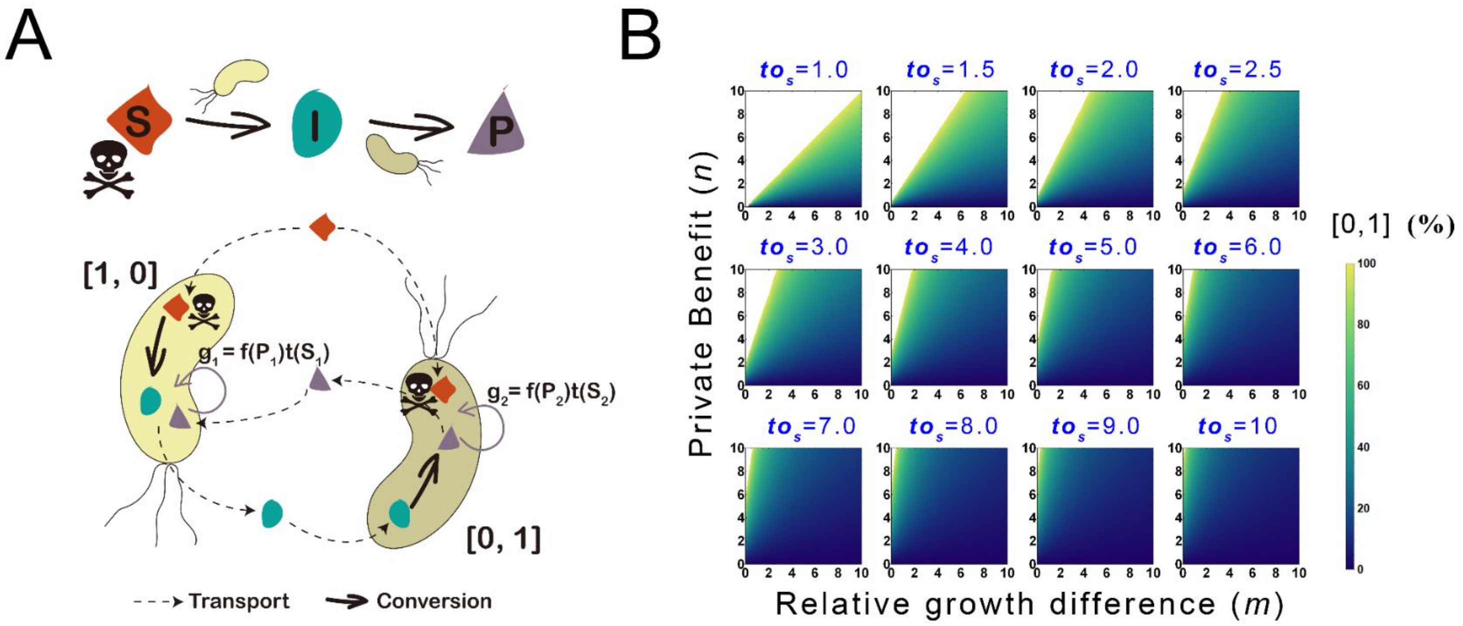

**Figure S6** Assembly rules when the substrate (S) is bio-toxic to both populations. (A) Schematic diagram shows the assumptions of our basic model considering substrate bio-toxicity. We hypothesize that higher intracelluar concentration of S decreases the growth rates of the two populations, which is characterized by a term *t(S)*. (B) The density maps show the parameter spaces characterized by private benefit $n$ and relative growth difference $m$, as well as relative abundance of [0, 1] in the stable communities, under different toxic index $to_s$. These plots are generated by visualizing Eqn [S2.74].

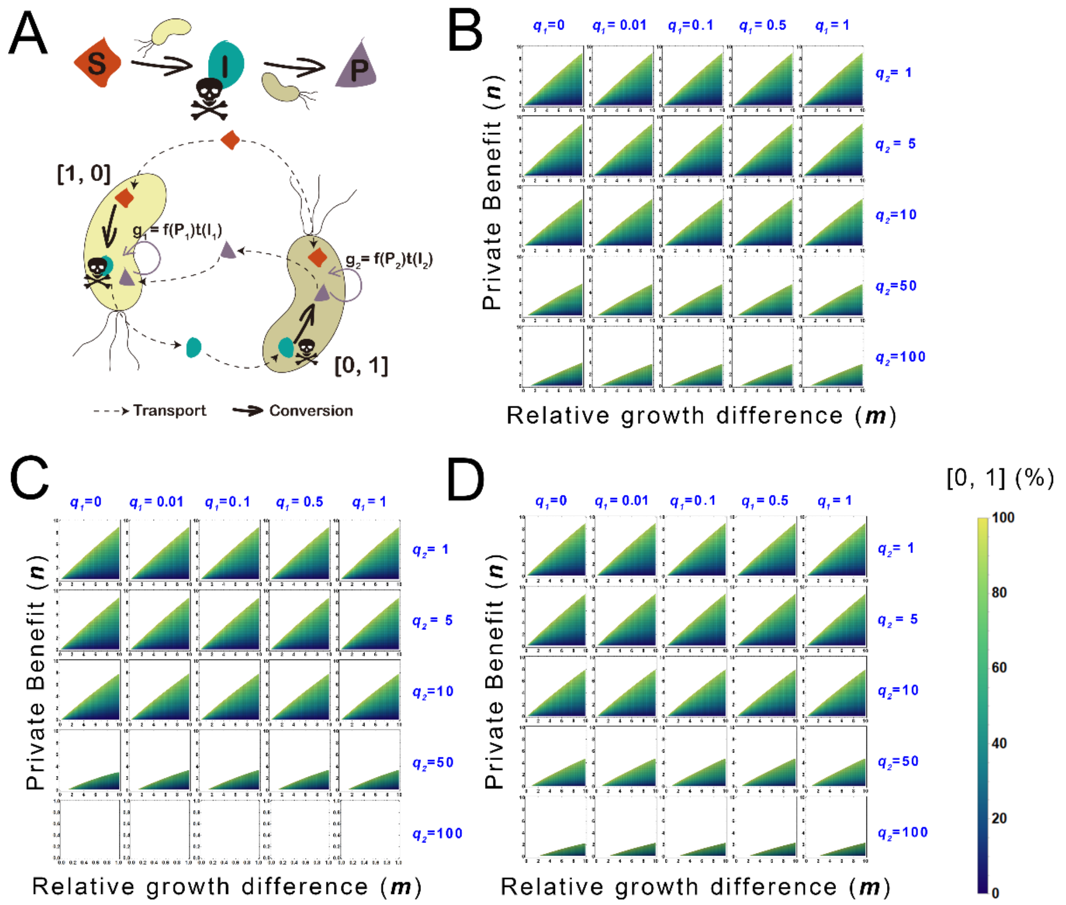

**Figure S7** Assembly rules when the intermediate (I) is bio-toxic to both populations. (A) Schematic diagram shows the assumptions of our basic model considering intermediate bio-toxicity. We hypothesize that higher intracelluar concentration of I decreases the growth rates of the two populations, which is characterized by a term *t(I)*. (B-D) Density maps show the parameter spaces, as well as relative abundance of [0, 1] in the stable communities, when the toxicity coefficients of I meet the reciprocal (B), linear (C) and exponential (D) forms and under different under different parameters $q_1$ and $q_2$ (see Section S2.7.2 for the definitions). These plots are generated by mathematical simulations. Values of other parameters are given in Table S2.

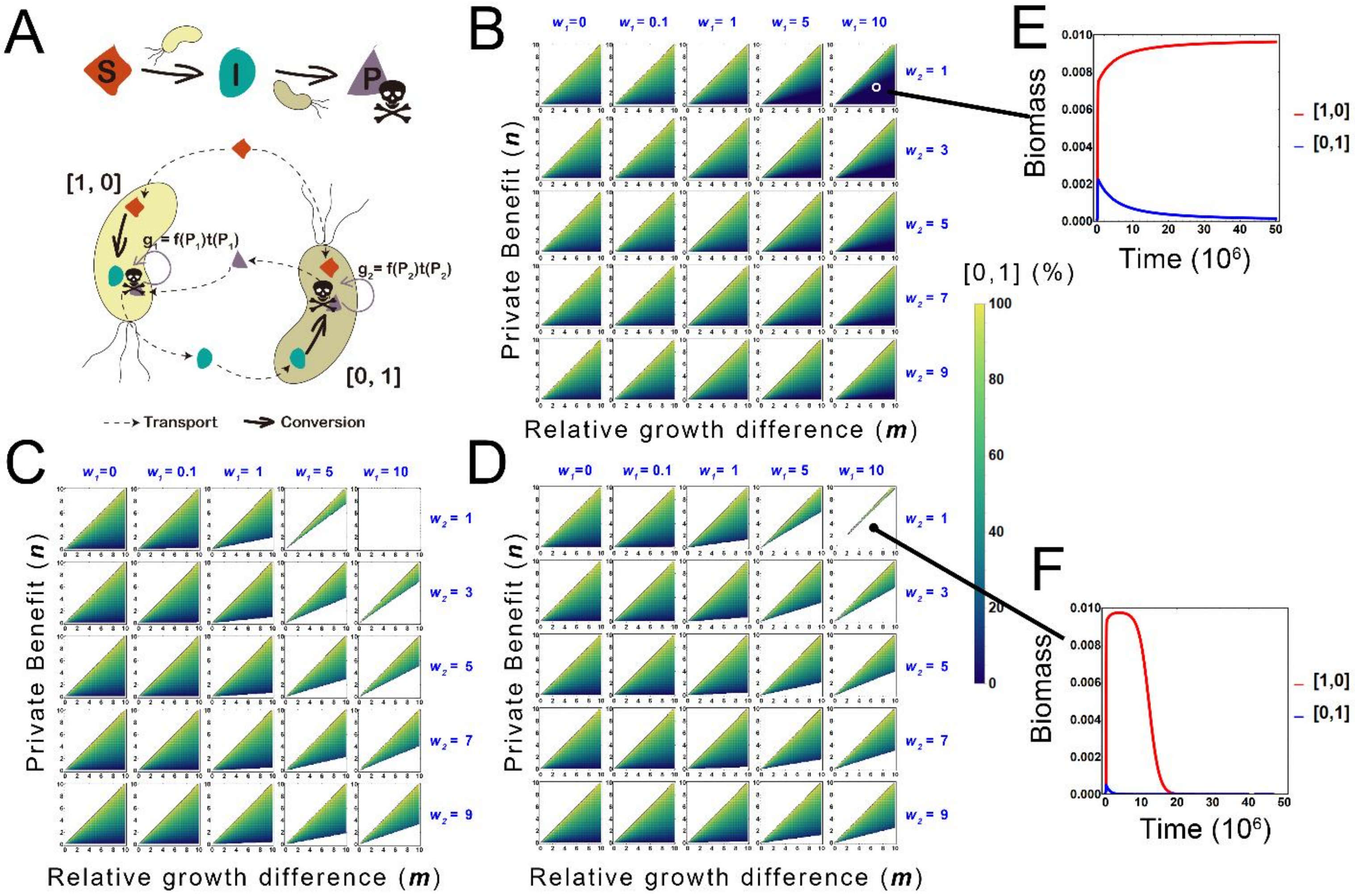
A
S
I
P
[1, 0]
[0, 1]
$g_1 = f(P_1)t(P_1)$
$g_2 = f(P_2)t(P_2)$
Transport
Conversion
B
C
D
$w_1=0$
$w_1 = 0.1$
$w_1 = 1$
$w_1 = 5$
$w_1 = 10$
$w_2 = 1$
$w_2 = 3$
$w_2 = 5$
$w_2 = 7$
$w_2 = 9$
Private Benefit (*n*)
Relative growth difference (*m*)
[0,1] (%)
E
F
Biomass
Time ($10^6$)
[1,0]
[0,1]

**Figure S8** Assembly rules when the end product (P) is bio-toxic to both populations. (A) Schematic diagram shows the assumptions of our basic model considering product bio-toxicity. We hypothesize that higher intracelluar concentration of P decreases the growth rates of the two populations, which is characterized by a term *t(P)*. (B-D) Density maps show the parameter spaces, as well as relative abundance of [0, 1] in the stable communities, when the toxicity coefficients of P meet the reciprocal (B), linear (C) and exponential (D) forms and under different under different parameters $w_1$ and $w_2$ (see Section S2.7.3 for the definitions). These plots are generated by mathematical simulations. Values of other parameters are given in Table S2.

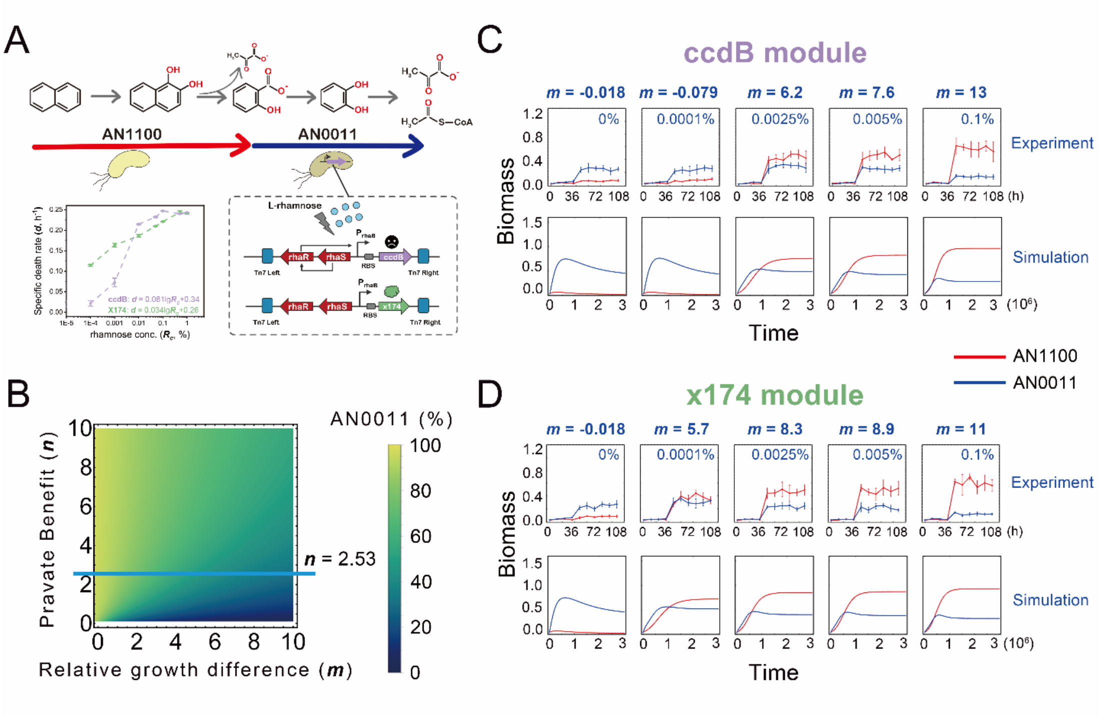

**Figure S9** Dynamics of the synthetic community composed of *P. stutzeri* AN1100 and *P. stutzeri* AN0011ccdB or *P. stutzeri* AN0011x174. (A) Schematic diagram of the construction of the synthetic consortium. To experimentally modify *m*, two genetic modules were introduced into strain *P. stutzeri* AN0011, generating *P. stutzeri* AN0011ccdB and *P. stutzeri* AN0011x174, in which the expression of toxic protein, CcdB or X174, are controllably induced by rhamnose. Therefore, the death rate of the modified strain can be quantitatively modulated by adjusting the rhamnose concentration, as shown in the bottom left. (B) Predicting the assembly of the synthetic community by mathematical modelling. The density map shows the parameter space in which the community can maintain (the colorized space), as well as the relative abundance of *P. stutzeri* AN0011ccdB or *P. stutzeri* AN0011x174 in the stable community (the color gradient). The blue line suggests that in our synthetic community *n* = 2.53, which is the region we performed experimental verification. (C-D) The dynamics of the synthetic community composed of *P. stutzeri* AN1100 and *P. stutzeri* AN0011ccdB (C), or *P. stutzeri* AN1100 and *P. stutzeri* AN0011x174 (D) from co-culture experiments and mathematical modelling under different rhamnose concentrations.

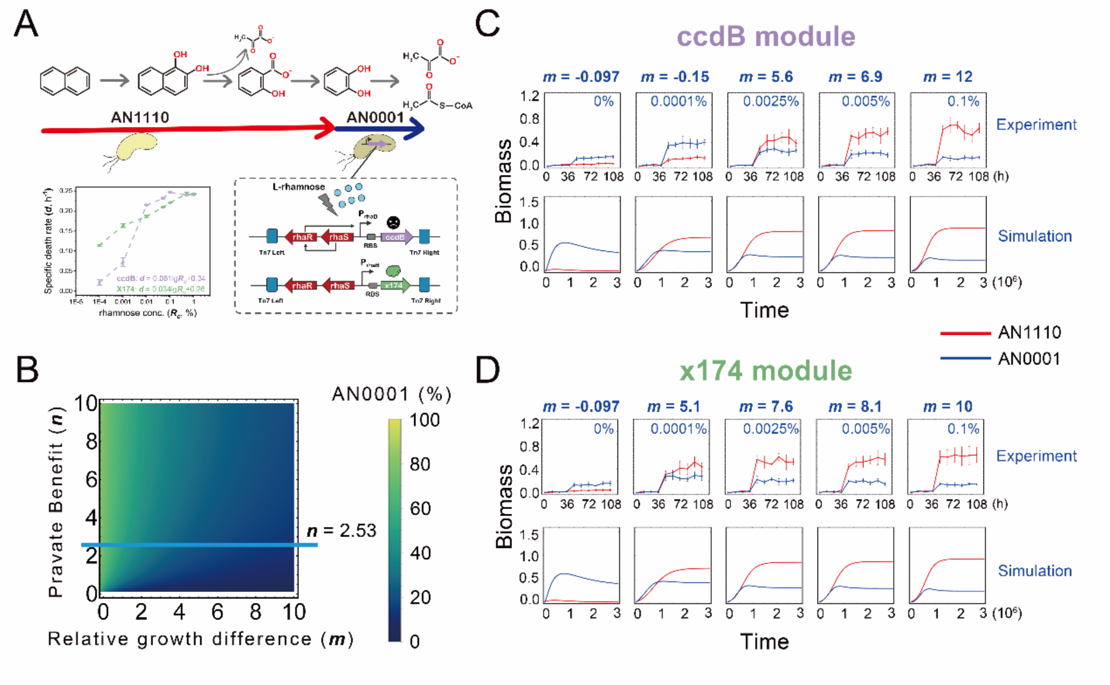

**Figure S10** Dynamics of the synthetic community composed of *P. stutzeri* AN1110 and *P. stutzeri* AN0001ccdB or *P. stutzeri* AN0001x174. (A) Schematic diagram of the construction of the synthetic consortium. To experimentally modify *m*, two genetic modules were introduced into strain *P. stutzeri* AN0001, generating *P. stutzeri* AN0001ccdB and *P. stutzeri* AN0001x174, in which the expression of toxic protein, CcdB or X174, are controllably induced by rhamnose. Therefore, the death rate of the modified strain can be quantitatively modulated by adjusting the rhamnose concentration, as shown in the bottom left. (B) Predicting the assembly of the synthetic community by mathematical modelling. The density map shows the parameter space in which the community can maintain (the colorized space), as well as the relative abundance of *P. stutzeri* AN0001ccdB or *P. stutzeri* AN0001x174 in the stable community (the color gradient). The blue line suggests that in our synthetic community $n$ = 2.53, which is the region we performed experimental verification. (C-D) The dynamics of the synthetic community composed of *P. stutzeri* AN1110 and *P. stutzeri* AN0001ccdB (C), or *P. stutzeri* AN1110 and *P. stutzeri* AN0001x174 (D) from co-culture experiments and mathematical modelling under different rhamnose concentrations.

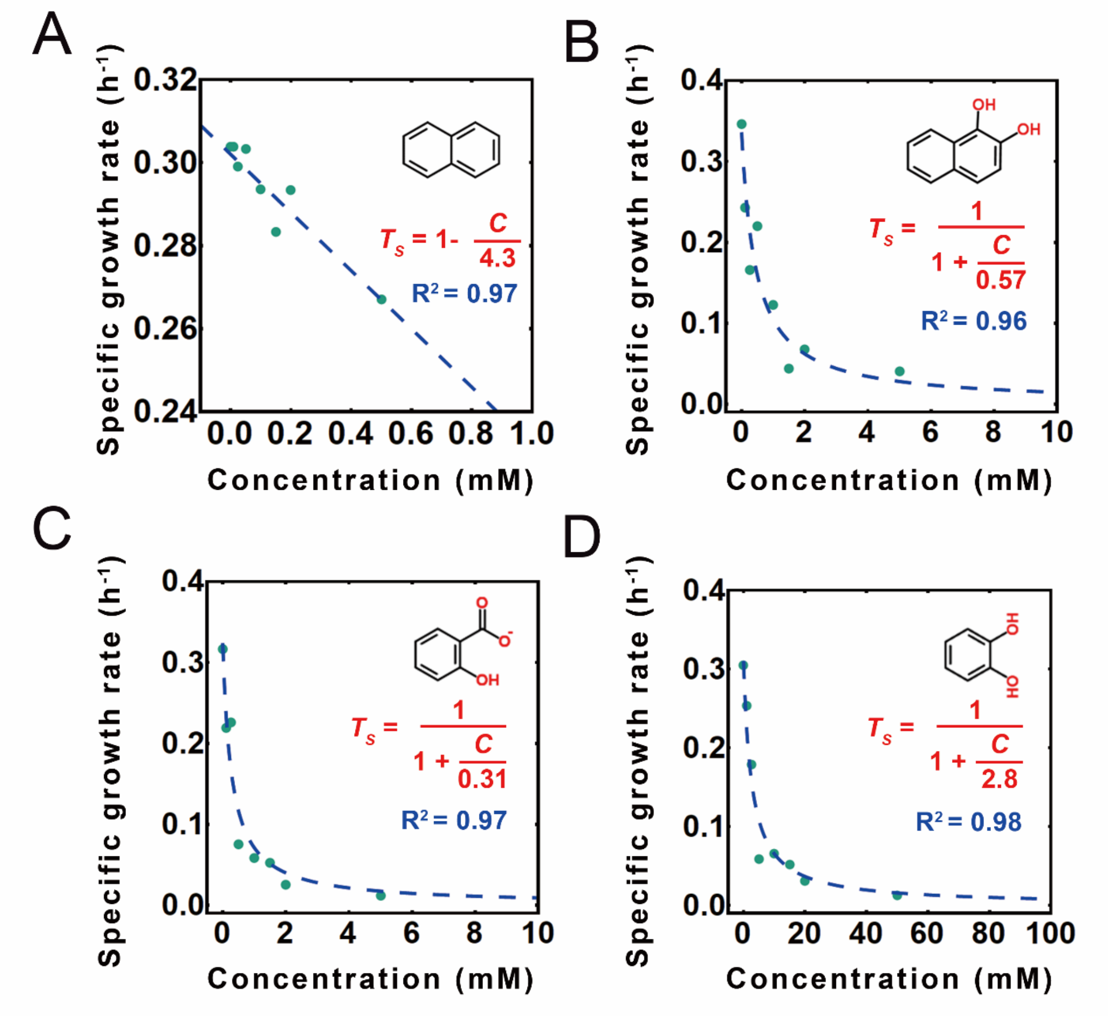

**Figure S11** Measurements of the bio-toxic coefficients of the substrate naphthalene (A), and the main intermediates, 1, 2-hydroxynaphthalene (B), salicylate (C) and catechol (D).

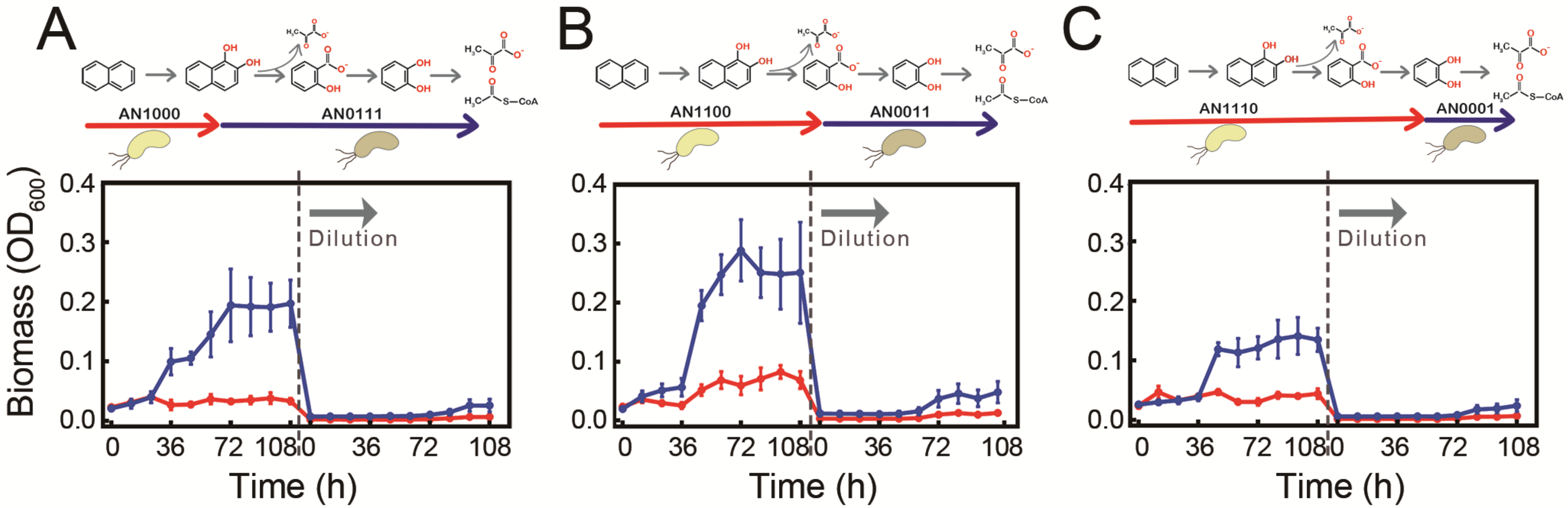

**Figure S12** Directly co-culturing the engineered strains to execute two-step MDOL leads to collapsed community dynamics. Dynamics of synthetic community composed of strains *P. stutzeri* AN1000 and *P. stutzeri* AN0111 (A), *P. stutzeri* AN1100 and *P. stutzeri* AN0011(B), as well as *P. stutzeri* AN1110 and *P. stutzeri* AN0001 (C). Naphthalene was used as the sole carbon souce in these experiments, and the cultures were diluted by a factor of 20 after 108 h, and transferred into a new fresh medium, as indicated by the dash line and the arrow.

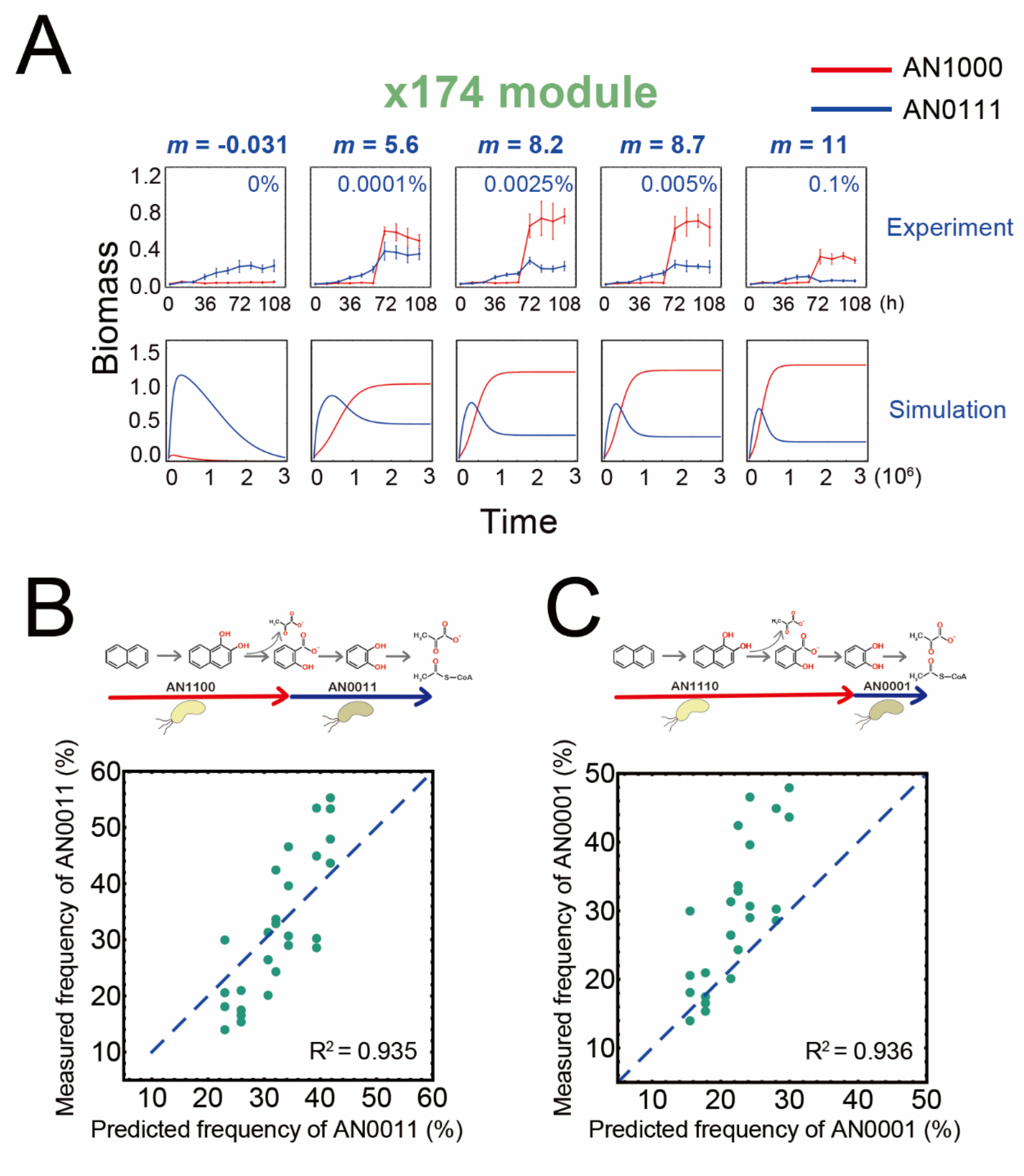

**Figure S13** Dynamics of the two-step synthetic community, as well as testing the predicting power of our mathematical frameworks. (A) The dynamics of the synthetic community composed of *P. stutzeri* AN1000 and *P. stutzeri* AN0111x174 from co-culture experiments and mathematical modelling under different rhamnose concentrations (that is, a gradient of *m* values). To measure their relative abundance, the strain performing the first step was labeled with MCherry, while the other strain was labeled with EGFP. (B-C) Testing the predicting power of our mathematical frameworks. The experimental measured frequency of the strains executing the second metabolic step are compared with the predicted frequency from our mathematical framework. The data used for these plots are summarized from those stable communities shown in Figure S14 and Figure S15, in which the values of relative abundance at the end of the third transfer were recorded. Note that the strains executing the second metabolic step in these experiments is the derived strains containing the death-rate modulating modules, that is, strains *P. stutzeri* AN0011ccdB and *P. stutzeri* AN0011x174 (B), as well as strains *P. stutzeri* AN0001ccdB and *P. stutzeri* AN0001x174 (C). Each green dot indicates one experimental replicate. The blue dashed line indicates the line in which the experimental results and predicted results are totally same. The adjust $R^2$ are aquired from statistical fit using NonlinearModelfit function of *Worfram Mathematica*.

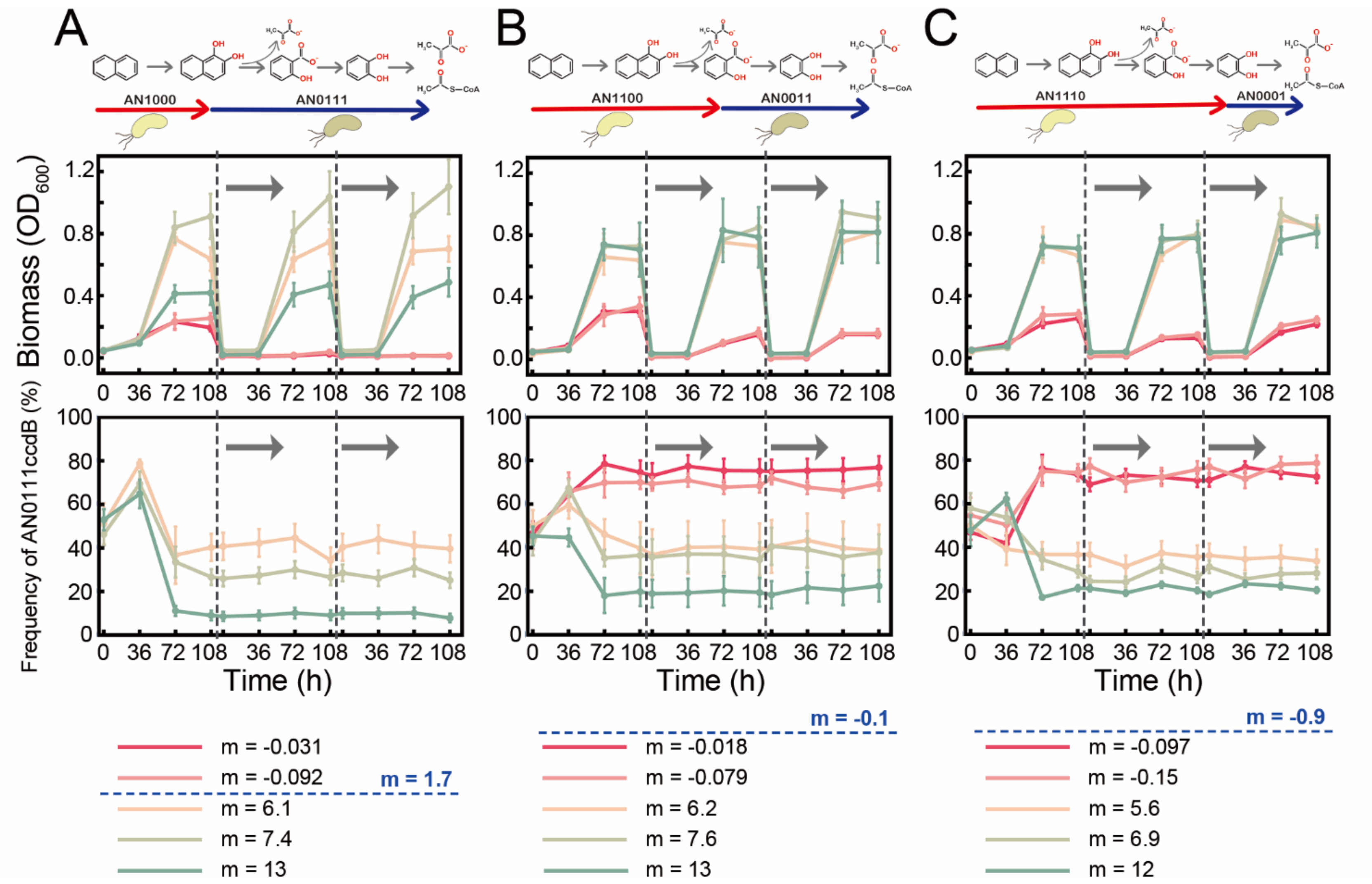
A
B
C
AN1000
AN0111
AN1100
AN0011
AN1110
AN0001
Biomass ($OD_{600}$)
Frequency of AN0111ccdB (%)
Time (h)
m = 1.7
m = -0.1
m = -0.9
m = -0.031
m = -0.092
m = 6.1
m = 7.4
m = 13
m = -0.018
m = -0.079
m = 6.2
m = 7.6
m = 13
m = -0.097
m = -0.15
m = 5.6
m = 6.9
m = 12

**Figure S14** Dynamics of our synthetic communities engaged in two-step MDOL undergoing three dilution-growth cycles. Communities composed of *P. stutzeri* AN1110 and *P. stutzeri* AN0001ccdB (A), *P. stutzeri* AN1100 and *P. stutzeri* AN0011ccdB (B), as well as *P. stutzeri* AN1110 and *P. stutzeri* AN0001ccdB (C) are shown. Naphthalene was used as the sole carbon souce in these experiments and the cultures were diluted by a factor of 20 after 108 h, and transferred into a new fresh medium, as indicated by the dash line and the arrow. The upper curve plots indicate the biomass of the communities across three cycles, while the lower curve plots indicate the relative abundance of strains executing the second metabolic step. Different colors indicate the experiments with different rhamnose concentrations, reflecting different $m$ values, as denoted in the bottom.

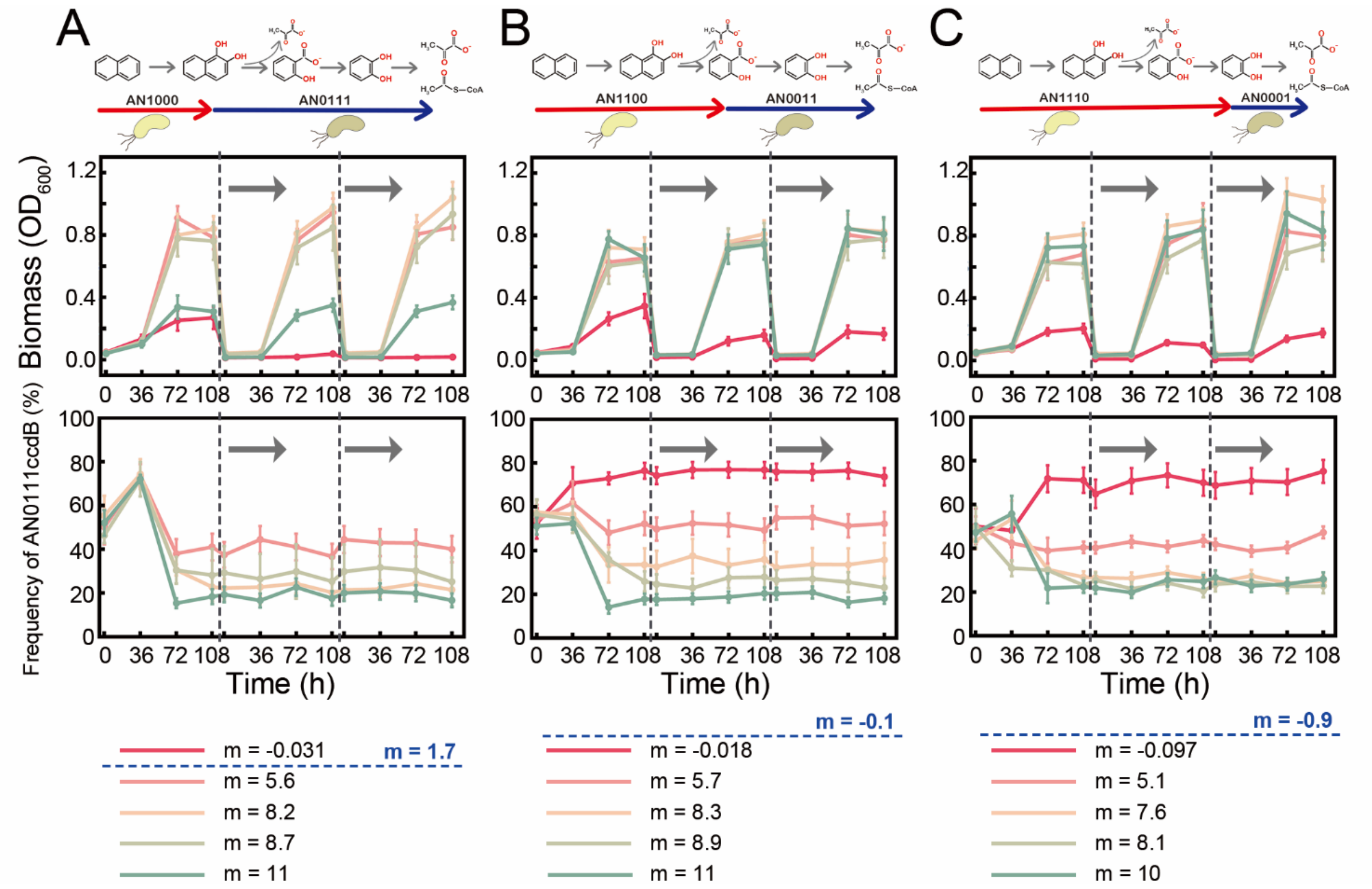
A
B
C
AN1000
AN0111
AN1100
AN0011
AN1110
AN0001
Biomass ($OD_{600}$)
Frequency of AN0111ccdB (%)
Time (h)
m = -0.031
m = 1.7
m = 5.6
m = 8.2
m = 8.7
m = 11
m = -0.1
m = -0.018
m = 5.7
m = 8.3
m = 8.9
m = 11
m = -0.9
m = -0.097
m = 5.1
m = 7.6
m = 8.1
m = 10

**Figure S15** Dynamics of our synthetic communities engaged in two-step MDOL undergoing three dilution-growth cycles. Communities composed of *P. stutzeri* AN1110 and *P. stutzeri* AN0001x174 (A), *P. stutzeri* AN1100 and *P. stutzeri* AN0011x174 (B), as well as *P. stutzeri* AN1110 and *P. stutzeri* AN0001x174 (C) are shown. Naphthalene was used as the sole carbon souce in these experiments and the cultures were diluted by a factor of 20 after 108 h, and transferred into a new fresh medium, as indicated by the dash line and the arrow. The upper curve plots indicate the biomass of the communities across three cycles, while the lower curve plots indicate the relative abundance of strains executing the second metabolic step. Different colors indicate the experiments with different rhamnose concentrations, resulted in different *m* values, as denoted in the bottom.

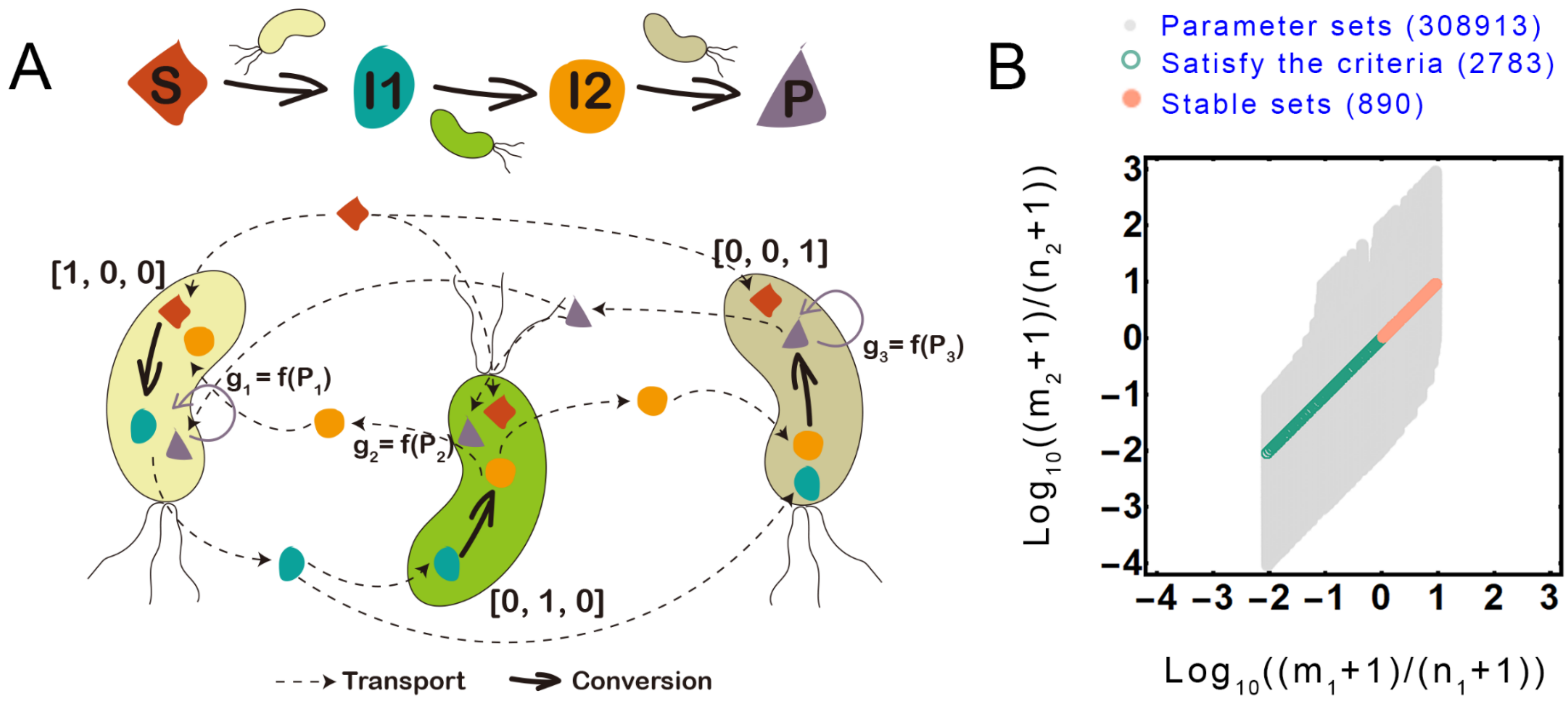

**Figure S16** The first condition required for the stability for the community engaged in three-step MDOL. (A) Schematic diagram shows the assumptions of the model regarding the assembly of a three-step MDOL community. We assumed a conceptualized organic substrate (S) was degraded into the first intermediate (I1) by a population [1, 0, 0], which is then converted to the second intermediate (I2) by the second population [0, 1, 0], and I2 is finally transferred to the end product (P) by the last population named [0, 0, 1]. All the reactions occurred intracellularly, while

S, I1, I2, and P were passive diffused across the cell membrane. Importantly, the growth of all the populations are dependent on the intracellular concentration of P, which is the sole limited resource of this system. (B) The graph depicts the distribution of 308913 designed parameter sets to test the first condition required for the stability of a three-step MDOL community. The grey dots indicate the distribution of all the sets; The green dots and orange dots indicate the parameter sets that satisfy the proposed condition defined by Eqn. [S3.43]; the orange dots indicate the parameter sets that lead to stable community dynamics.

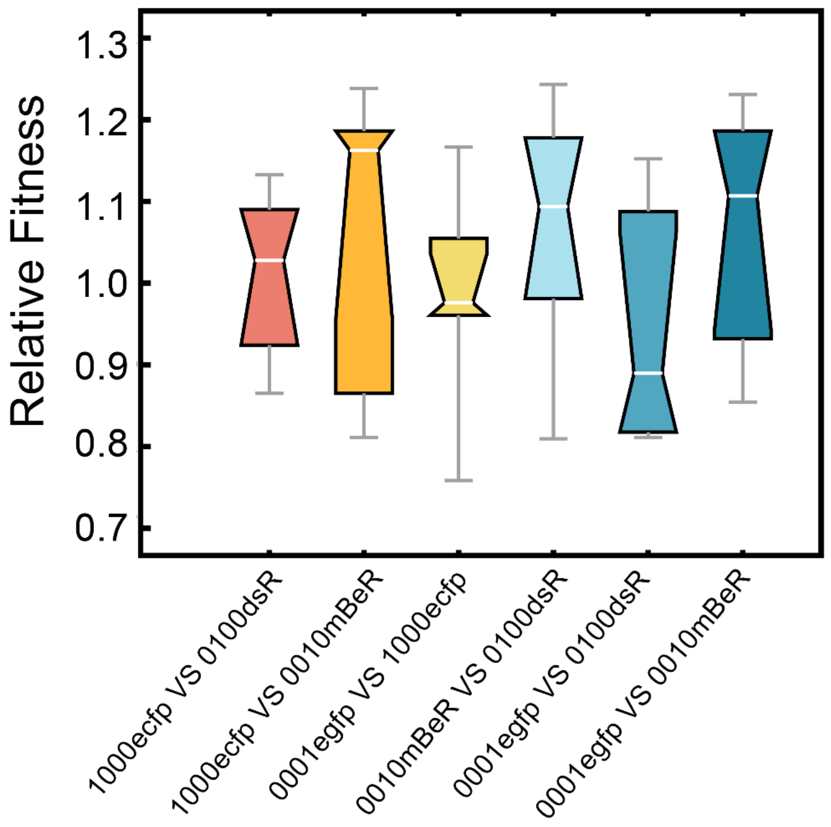

**Figure S17** Relative fitness between the strains involved in our synthetic community engaged in a four-step MDOL community. Here, strain *P. stutzeri* AN1000 was labeled with ECFP, strain *P. stutzeri* AN0100 was labeled with DsRed, strain *P. stutzeri* AN0010 was labeled with mBeRFP, and strain *P. stutzeri* AN0001 was labeled with EGFP, same as the labelling used in the co-culture experiments. The results were obtained by competitive fitness assays using minimum medium supplemented with 2 mM IPTG, 50 μg/mL gentamicin, as well as pyruvate (0.5 % w/v) as the sole carbon source. In this condition, the strains can grow on the end product (pyruvate) and still synthesize the enzymes for naphthalene degradation (since their expressions are induced by IPTG). Six replicates were performed for each condition. The measurements of relative fitness followed the standard protocol reported in Reference[38] and [39].

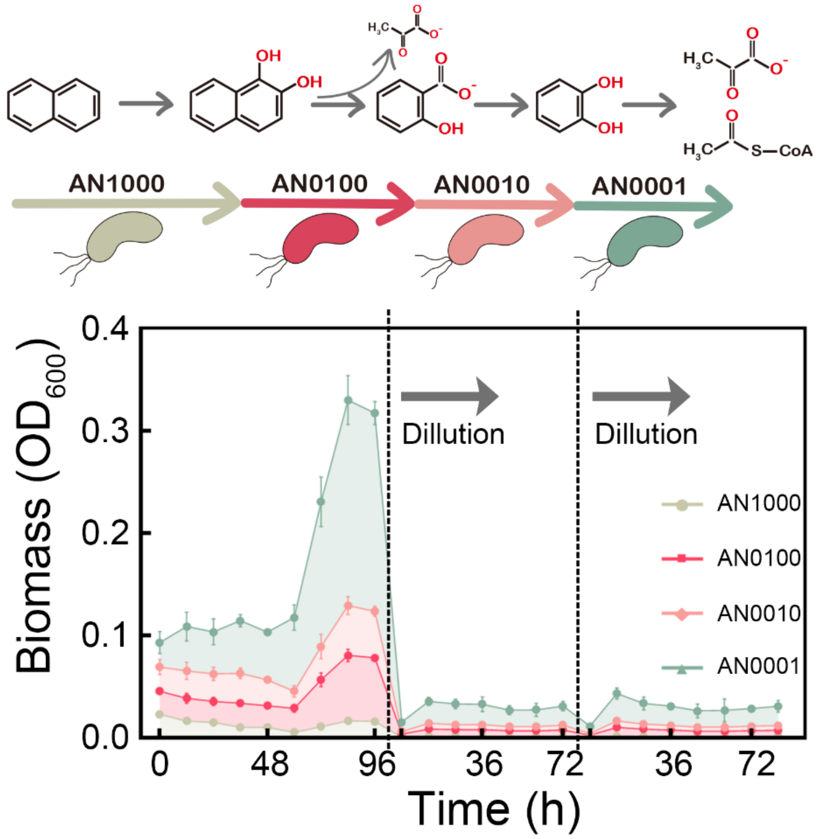

**Figure S18** Directly co-culture of the engineered strains that execute four-step MDOL leads to collapsed community dynamics. Dynamics of synthetic community composed of strains *P. stutzeri* AN1000 and *P. stutzeri* AN01000, *P. stutzeri* AN0010 and *P. stutzeri* AN0001. Naphthalene was used as the sole carbon souce in these experiments. The cultures were diluted by a factor of 20 after 96 h or 72h, and transferred into a new fresh medium, as indicated by the dash line and the arrow.

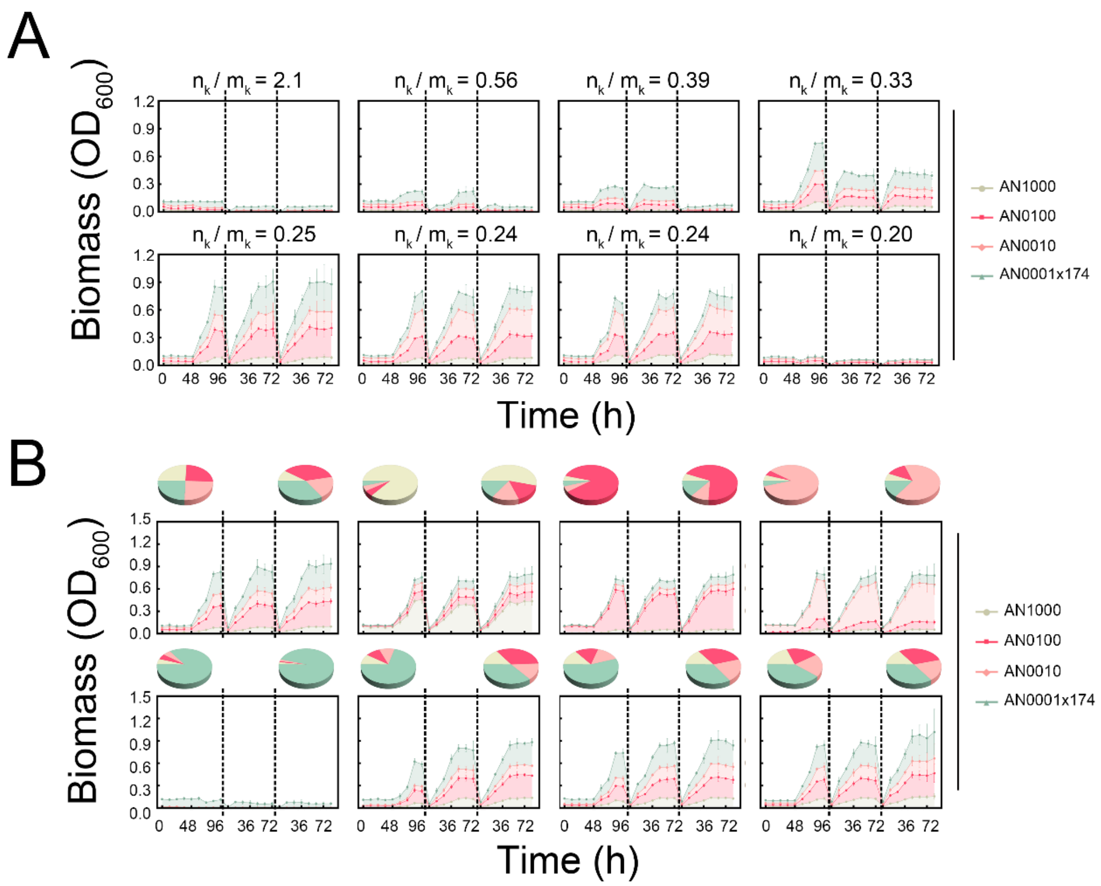

**Figure S19** Dynamics of the synthetic community composed of *P. stutzeri* AN1000, *P. stutzeri* AN0100, *P. stutzeri* AN0010 and *P. stutzeri* AN0001x174. (A) community dynamics under a gradient of $n_k/m_k$ value conditions (that is, using medium with different rhamnose concentrations). The cultures were diluted by a factor of 20 after 96 h or 72h (indicated by the dash line), and transferred into a new fresh medium. To measure their relative abundance, strain *P. stutzeri* AN1000 was labeled with ECFP, strain *P. stutzeri* AN0100 was labeled with DsRed, strain *P. stutzeri* AN0010 was labeled with mBeRFP, and strain *P. stutzeri* AN0001x174 was labeled with EGFP. (B) Growth dynamics of the synthetic community at eight different initial ratios. in these experiments, the concentration of rhamnose was set to 0.005% ($n_k/m_k$ value equals to 0.25). Each color region shows the relative abundance of each strain. The pie charts denote the community structure at the starting and end time points. Three independent replicates were performed for each condition.

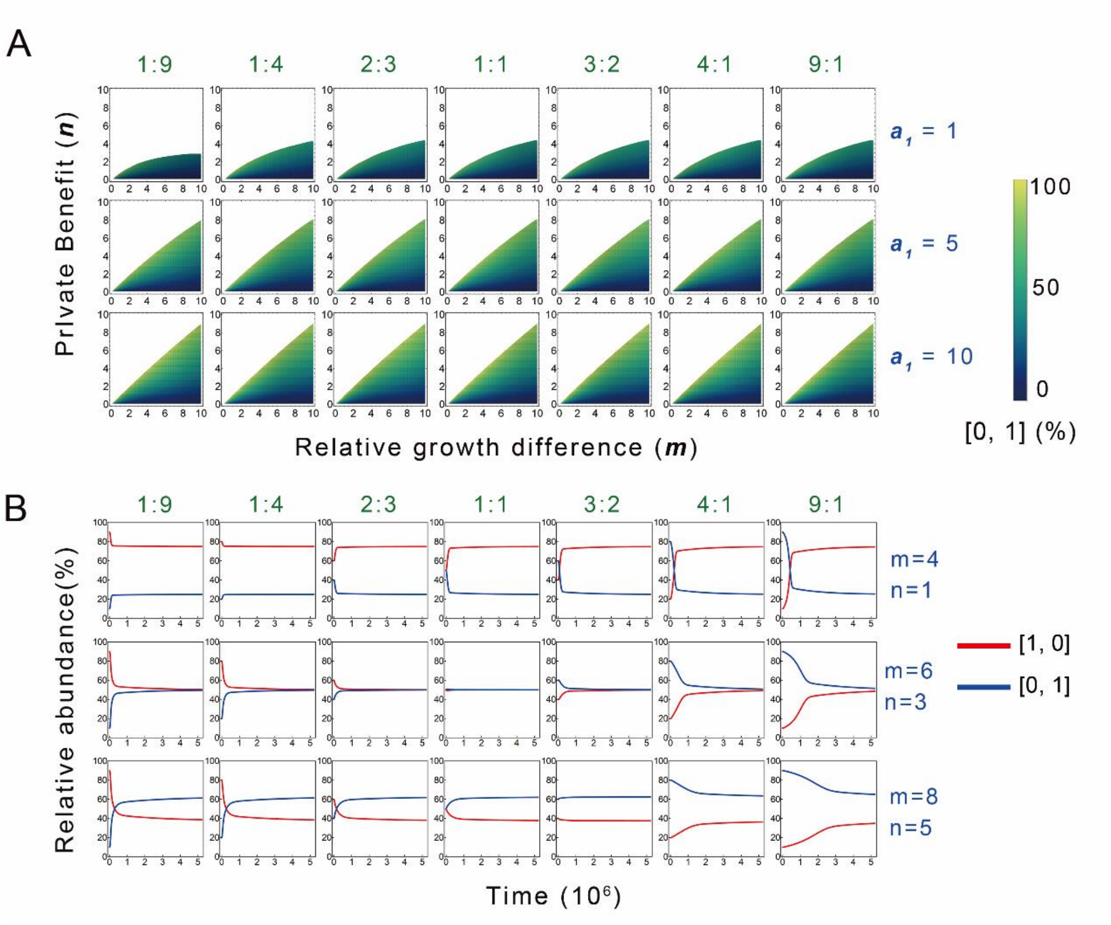

**Figure S20** The basic model predicts that the community engaged in two-step MDOL is robust to the change of initial inoculation ratio. (A) The density maps show the parameter spaces characterized by private benefit *n* and relative growth difference *m*, as well as relative abundance of [0, 1] in the stable communities, under different initial inoculation ratio and different rates of the first metabolic step ($a_1$). These plots are generated by mathematical simulations. Values of other parameters are given in Table S2. (B) The representative simulation dynamics under some specific *n* and *m* combinations. Note that the final stable state of a two-step MDOL community was independent of the initial inoculation ratio.

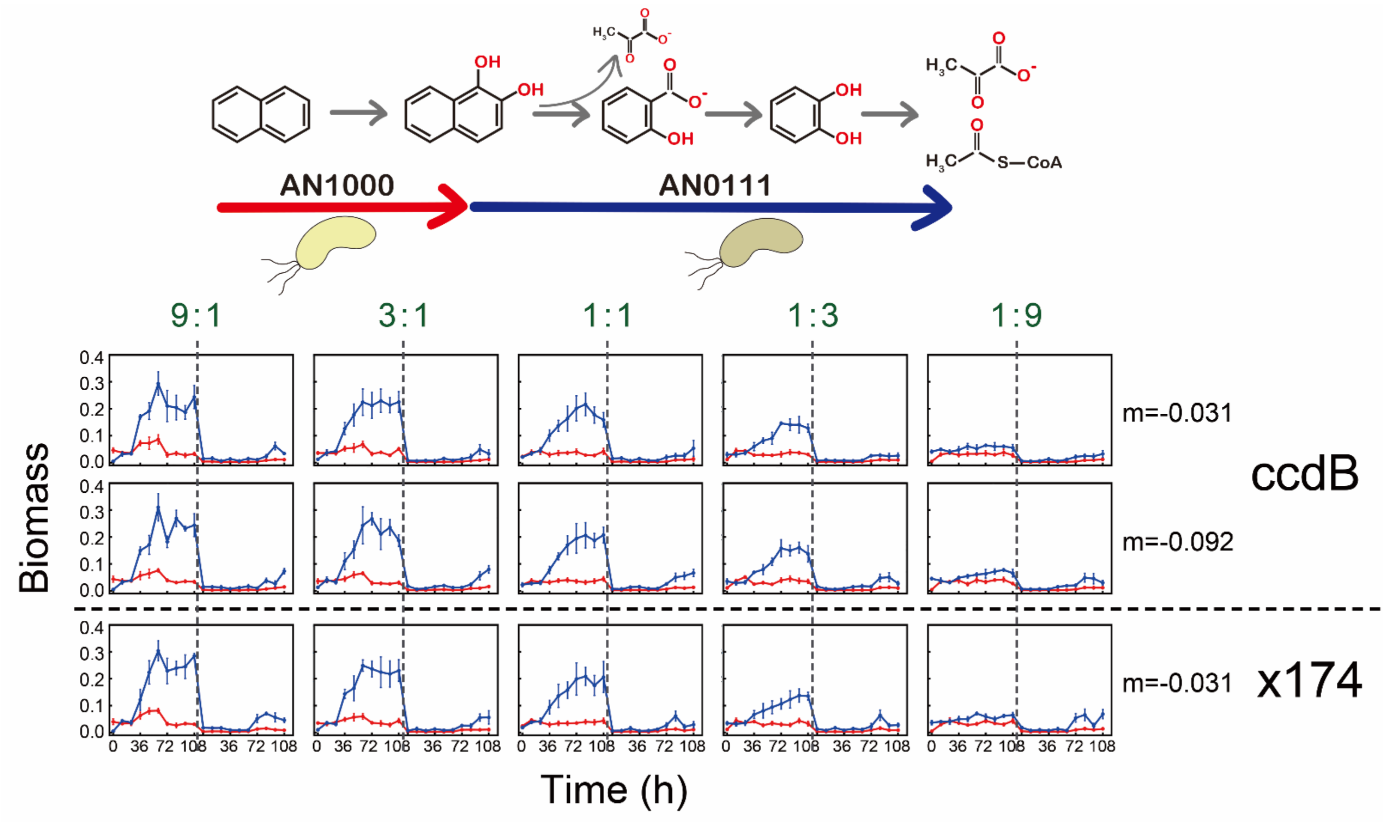

**Figure S21** Growth dynamics of the synthetic community composed of *P. stutzeri* AN1000 and *P. stutzeri* AN0111ccdB (or *P. stutzeri* AN0111x174) initialized with five different starting ratios when the concentration of rhamnose was set to a low level (that is, our rule predicts that the community should be unstable).

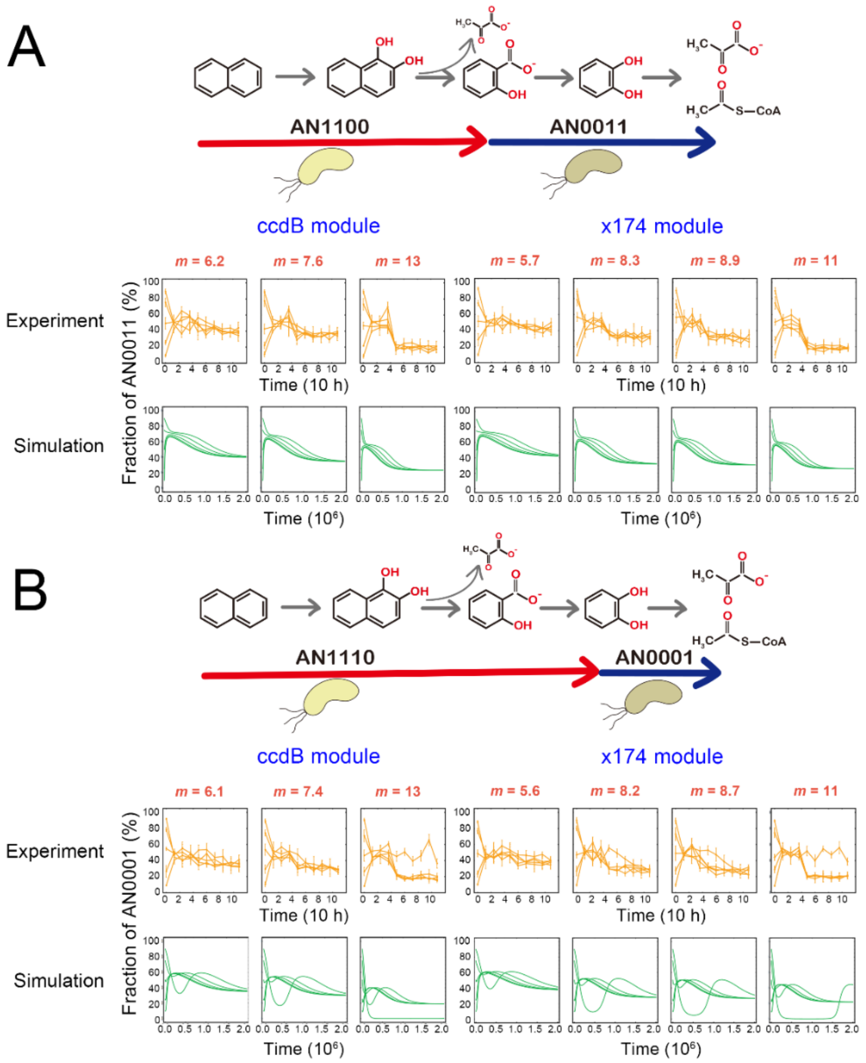

**Figure S22** Assembly rule of a two-step MDOL community is robust to the initial strain ratio. Culture experiments and mathematical modelling of two synthetic communities engaged in two-step MDOL were performed initialized with seven different *m* values, as well as five different initial ratios. The results from the community composed of (A) *P. stutzeri* AN1100 and *P. stutzeri* AN0011ccdB (or *P. stutzeri* AN0011x174), and that composed of (B) *P. stutzeri* AN1110 and *P. stutzeri* AN0001ccdB (or *P. stutzeri* AN0001x174) were shown. The relative abundances of the population executing the second metabolic step from experiments (orange, first row) and mathematical modelling (green, second row) were shown.

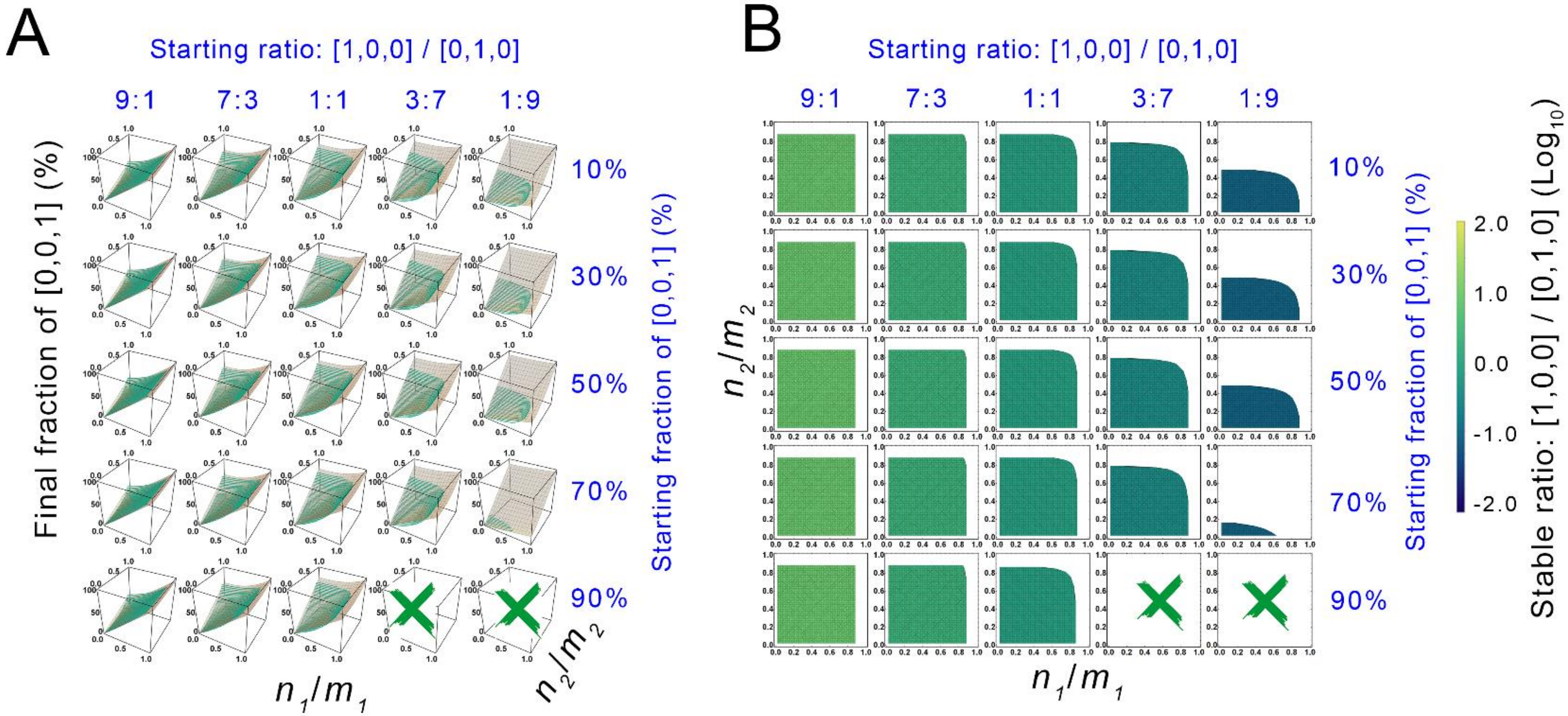

**Figure S23** The effects of initial strain ratio on the assembly of microbial community engaged in three-step MDOL. Computational simulations were performed using the model regarding the assembly of a three-step MDOL community and initialized with 25 different starting ratios. The parameter sets that leads to stable community dynamics were collected. Then the relationship between the relative abundance of the [0, 0, 1] and the ratio $n_1/m_1$ and $n_2/m_2$ were collected, shown by green dots in (A) and color gradients in (B). In Figure (A), the surface diagram shows distribution of the relative abundance of [0, 0, 1] predicted by Eqn. [7]. The green crosses suggests that in the correponding condition, no parameter sets leads to stable community dynamics. Details of these simulations are decribed in Supplementary Information S3.6.

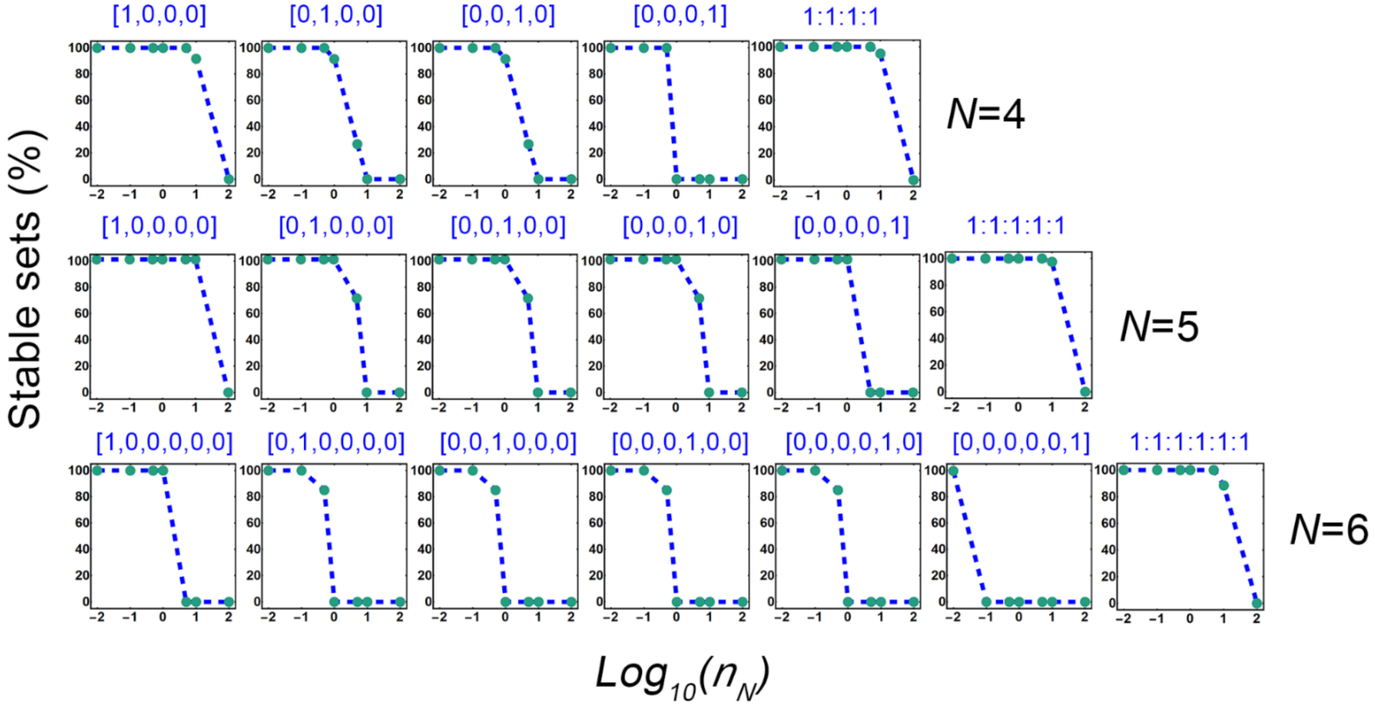

**Figure S24** Initial strain ratios affect the possibility of the size of the parameter sets leading to the stable community dynamics. We quantifiy this possibility by calculating the ratio of the parameter sets that can stabilize the community accounting for the total number of the parameter sets. The shifts of this ratio along $n_N$ values under different initial strain ratios of the community engaged in four-, five-, and six-step MDOL were shown. Details of these simulations are decribed in Supplementary Information S3.6.

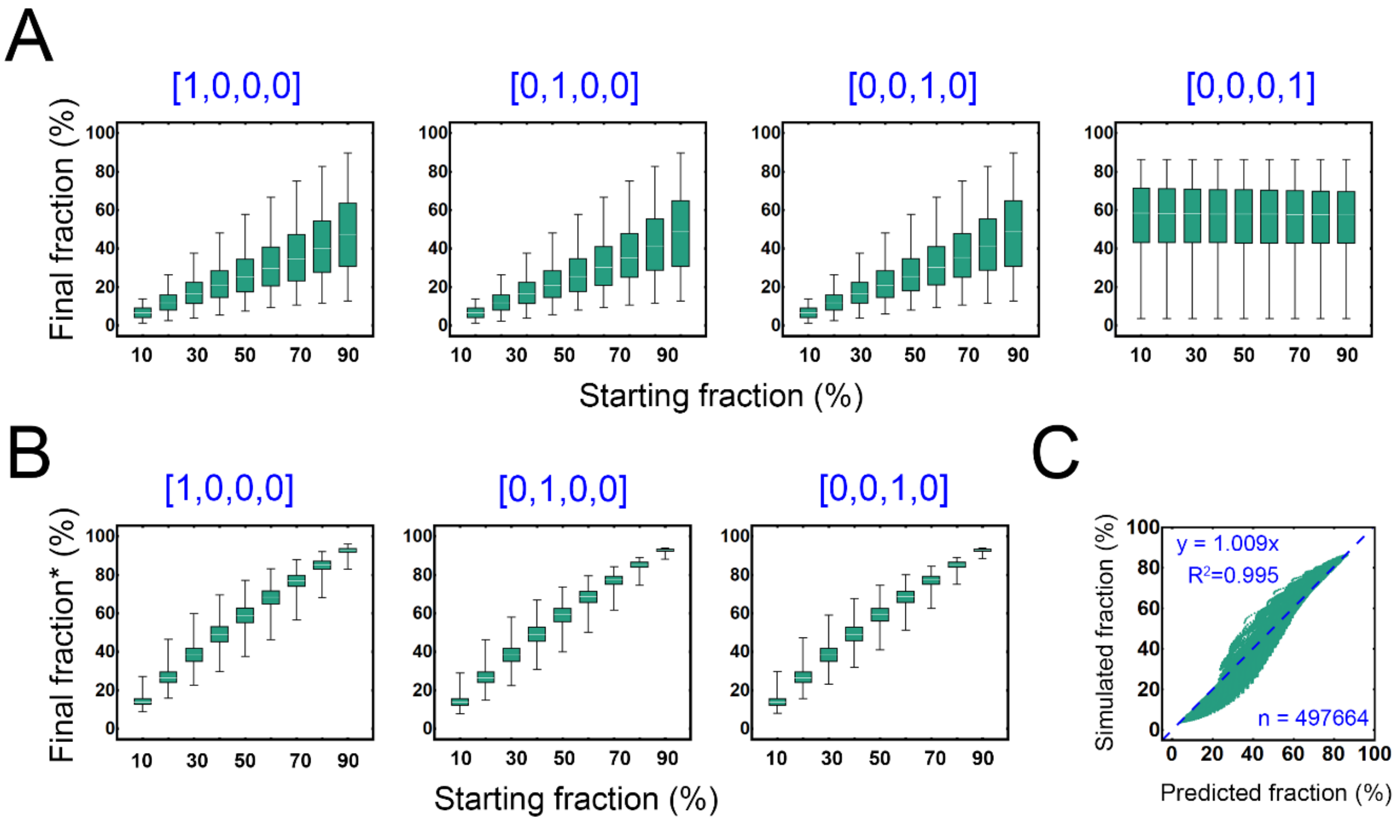

**Figure S25** The effects of initial strain ratio on the stability of a four-step MDOL community. Simulations were performed by setting a gradient of initial fraction of one population, while the initial fraction of other populations was controlled to be equal. The results that lead to stable community dynamics were collected. (A) The relationship between the initial fraction and final fraction of each population. (B) The relationship between the initial fraction and final fraction accounting for all the formmer populations (defined as final fraction*) of each population. (C) The linear correlation between the fraction of the last population [0, 0, 0, 1] predicted by Eqn. [7] and its fraction obtained by simulations regarding a four-step MDOL community under different initial strain ratio. The green dots show the distribution of simulated fraction and the corresponding predicted values from Eqn. [7]. The dashed line shows the fitted correlation curve. Details of these simulations are decribed in Supplementary Information S3.6.

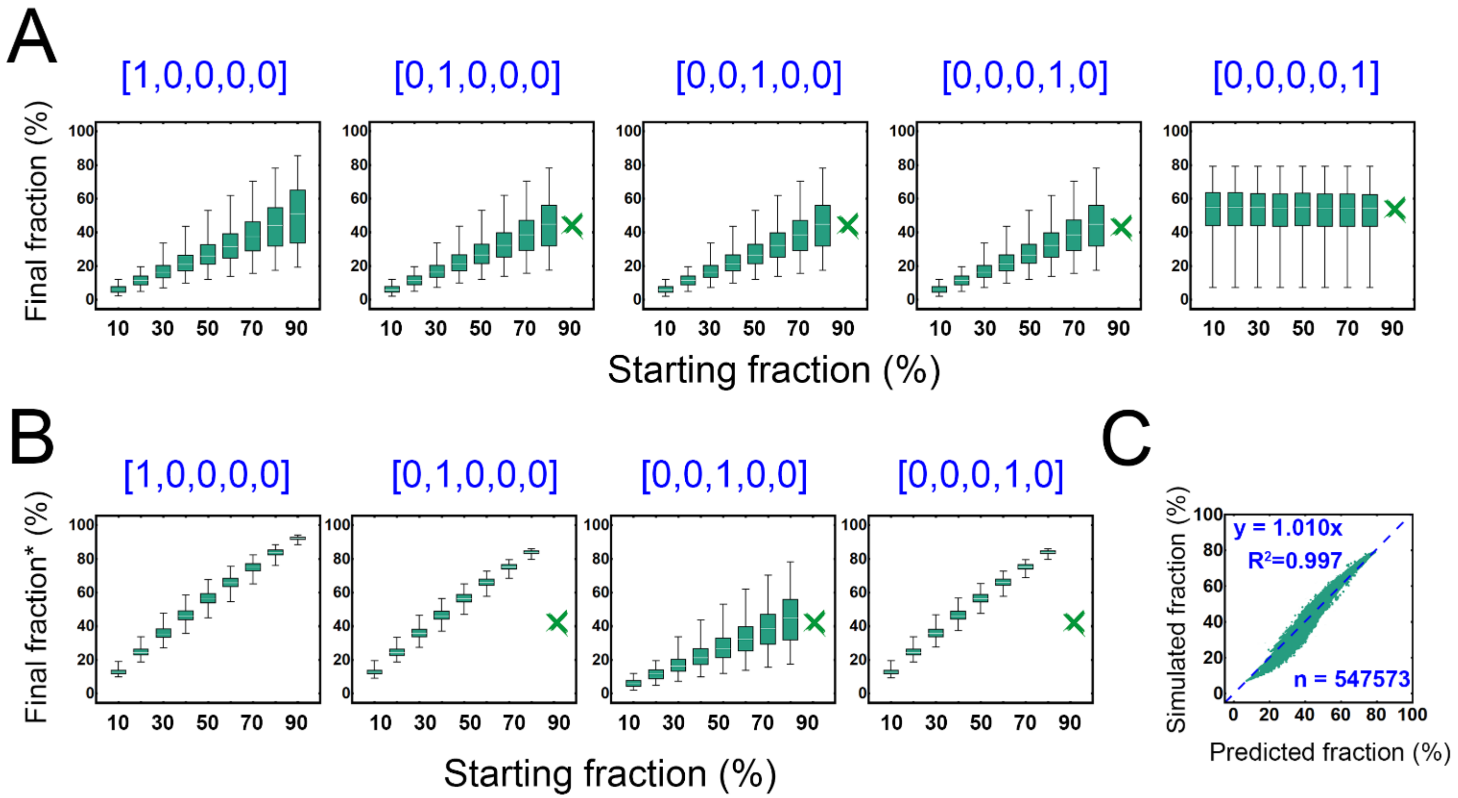

**Figure S26** The effects of initial strain ratio on the stability of a five-step MDOL community. Simulations were performed by setting a gradient of initial fraction of one population, while the initial fraction of other populations was controlled to be equal. The results that lead to stable community dynamics were collected. (A) The relationship between the initial fraction and final fraction of each population. (B) The relationship between the initial fraction and final fraction accounting for all the formmer populations (defined as final fraction*) of each population. (C) The linear correlation between the fraction of the last population [0, 0, 0, 0, 1] predicted by Eqn. [7] and its fraction obtained by simulations regarding a four-step MDOL community under different initial strain ratio. The green dots show the distribution of simulated fraction and the corresponding predicted values from Eqn. [7]. The dashed line shows the fitted correlation curve. Details of these simulations are decribed in Supplementary Information S3.6.

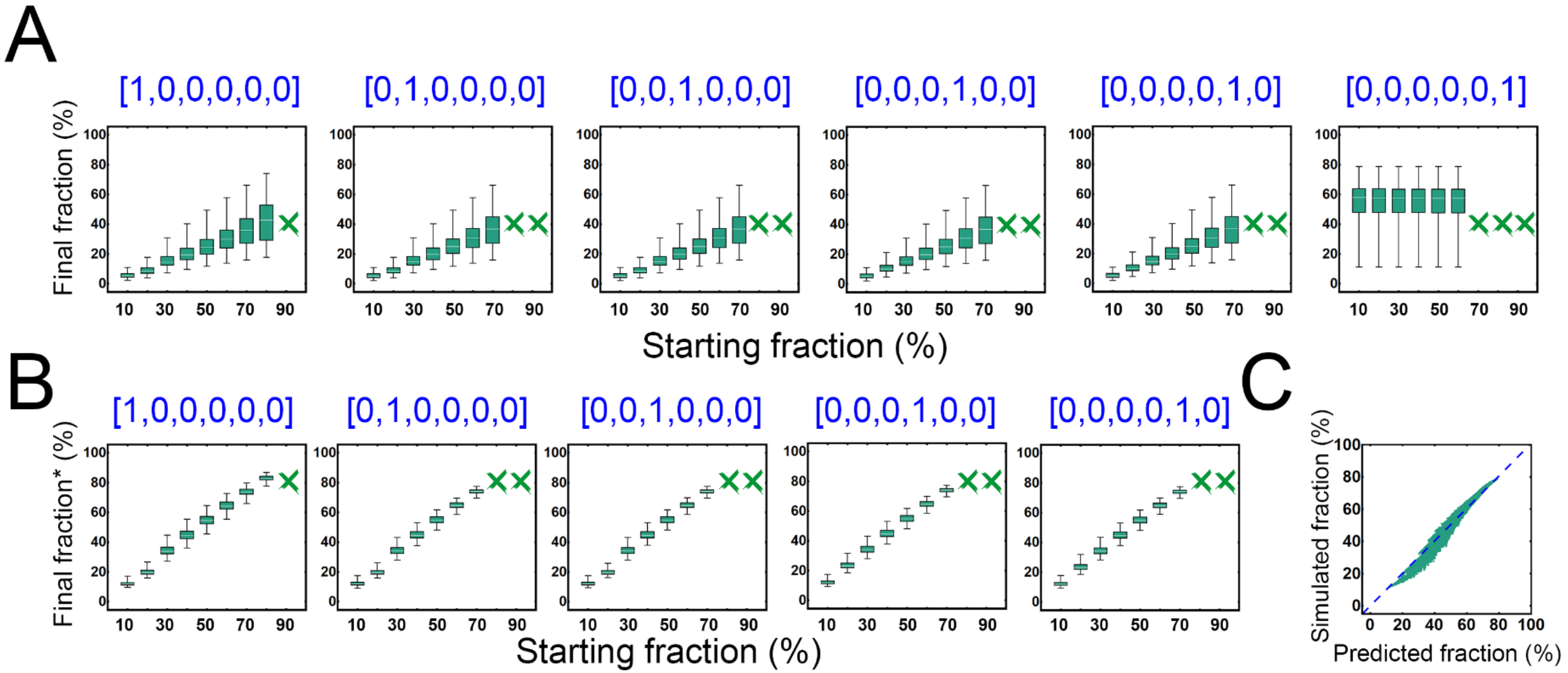

**Figure S27** The effects of initial strain ratio on the stability of a six-step MDOL community. Simulations were performed by setting a gradient of initial fraction of one population, while the initial fraction of other populations was controlled to be equal. The results that lead to stable community dynamics were collected. (A) The relationship between the initial fraction and final fraction of each population. (B) The relationship between the initial fraction and final fraction accounting for all the formmer populations (defined as final fraction*) of each population. (C) The linear correlation between the fraction of the last population [0, 0, 0, 0, 0, 1] predicted by Eqn. [7] and its fraction obtained by simulations regarding a four-step MDOL community under different initial strain ratio. The green dots show the distribution of simulated fraction and the corresponding predicted values from Eqn. [7]. The dashed line shows the fitted correlation curve. Details of these simulations are decribed in Supplementary Information S3.6.

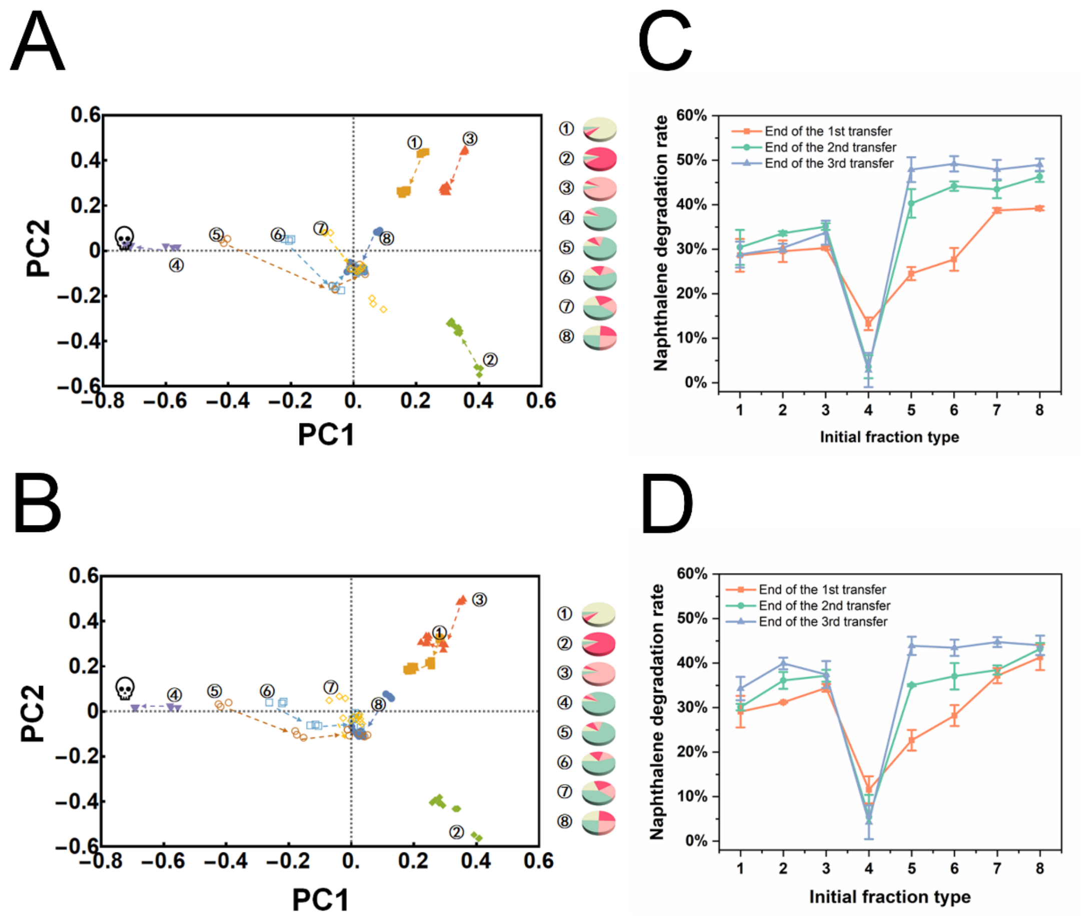

**Figure S28** The effects of initial strain ratio on the assembly and efficiency of our four-step MDOL community. PCA analyses of the dynamics of the synthetic consortium composed of *P. stutzeri* AN1000, *P. stutzeri* AN0100, *P. stutzeri* AN0010, and *P. stutzeri* AN0001ccdB (A), or *P. stutzeri* AN0001x174 (B), at eight different initial ratios when the concentration of rhamnose was set to 0.005% ($n_k/m_k$ value equals to 0.29 (A) or 0.25 (B)). The pie charts on the right denote the eight initial inoculation ratios, and the color marks are consistent with Figure 6. The community structure data used for PCA analysis is the same as that in Figure 6B and Figure S19B. Three independent experimental replicates were performed. The arrows show the direction of community succession. (C-D) Naphthene degradation rate of the consortia containing *P. stutzeri*

AN0001ccdB (C) and *P. stutzeri* AN0001x174 (D). The degradation rates were measured at the end of each transfer, using a GC-MS (Agilent 7890A and Agilent 5975C) following the standard protocol[52-54]. Three independent experimental replicates were performed.
